# Targeting the MR1-MAIT Cell Axis Improves Vaccine Efficacy and Affords Protection against Viral Pathogens

**DOI:** 10.1101/2023.02.20.529311

**Authors:** Rasheduzzaman Rashu, Marina Ninkov, Christine M. Wardell, Jenna M. Benoit, Nicole I. Wang, Michael R. D’Agostino, Ali Zhang, Emily Feng, Nasrin Saeedian, Gillian I. Bell, Fatemeh Vahedi, David A. Hess, Ryan M. Troyer, C. Yong Kang, Ali A. Ashkar, Matthew S. Miller, S.M. Mansour Haeryfar

**Affiliations:** Department of Microbiology and Immunology, Western University, London, Ontario, Canada; McMaster Immunology Research Centre, McMaster University, Hamilton, Ontario, Canada; Krembil Centre for Stem Cell Biology, Molecular Medicine Research Laboratories, Robarts Research Institute, London, Ontario, Canada; Department of Physiology and Pharmacology, Western University, London, Ontario, Canada; Department of Medicine, McMaster University, Hamilton, Ontario, Canada; Department of Biochemistry and Biomedical Sciences, McMaster University, Hamilton, Ontario, Canada; Michael G. DeGroote Institute for Infectious Disease Research, McMaster University, Hamilton, Ontario, Canada; Division of Clinical Immunology and Allergy, Department of Medicine, Western University, London, Ontario, Canada; Division of General Surgery, Department of Surgery, Western University, London, Ontario, Canada; Lawson Health Research Institute, London, Ontario, Canada

**Keywords:** MAIT cells, MR1, 5-OP-RU, adjuvant, vaccine, cytokine, interferon, influenza, SARS-CoV-2, COVID-19, vaccinia virus, viral infection, heterologous immunity

## Abstract

Mucosa-associated invariant T (MAIT) cells are MR1-restricted, innate-like T lymphocytes with tremendous antibacterial and immunomodulatory functions. MAIT cells also sense and respond to viral infections in an MR1-independent fashion. However, whether they can be directly targeted in immunization strategies against viral pathogens is unknown. We addressed this question in multiple wild-type and genetically altered but clinically relevant mouse strains using several vaccine platforms against influenza viruses, poxviruses and severe acute respiratory syndrome coronavirus 2 (SARS-CoV-2). We demonstrate that 5-(2-oxopropylideneamino)-6-D-ribitylaminouracil (5-OP-RU), a riboflavin-based MR1 ligand of bacterial origin, can synergize with viral vaccines to expand MAIT cells in multiple tissues, reprogram them towards a pro-inflammatory MAIT1 phenotype, license them to bolster mainstream virus-specific CD8^+^ T cell responses, and potentiate heterosubtypic antiviral protection. Repeated 5-OP-RU administration did not render MAIT cells anergic, thus allowing for its inclusion in prime-boost immunization protocols. Mechanistically, tissue MAIT cell accumulation was due to their robust proliferation, as opposed to altered migratory behavior, and required viral vaccine replication competency, Toll-like receptor 3 (TLR3) and cell-autonomous type I interferon receptor signaling. Furthermore, the observed phenomenon was manifest in young and old female and male mice, and could also be recapitulated in a human cell culture system in which peripheral blood mononuclear cells were exposed to replicating virions and 5-OP-RU. In conclusion, although viruses and virus-based vaccines are devoid of the riboflavin biosynthesis machinery that supplies MR1 ligands, targeting MR1 enhances the efficacy of vaccine-elicited antiviral immunity. We propose 5-OP-RU as a non-classic but potent and versatile vaccine adjuvant against respiratory viruses.

## Introduction

Infection- and vaccine-elicited protection against viral invaders relies heavily on primary and anamnestic T and B cell responses, which are controlled and shaped by the innate arm of immunity. While many classic adjuvants prolong and enhance peptide antigen (Ag) presentation by the highly polymorphic major histocompatibility complex (MHC) molecules, targeting innate immune effectors is considered an attractive strategy to bolster antiviral defense mechanisms across genetically diverse populations.

Mucosa-associated invariant T (MAIT) cells are an evolutionarily conserved subset of innate-like T lymphocytes with impressive antimicrobial, cytotoxic and immunomodulatory properties^1,2^. The canonically rearranged invariant T cell receptor (*i*TCR) α chain of MAIT cells (typically Vα19-Jα33 in mice and Vα7.2-Jα33 in humans) is paired with one of only a limited number of available β chains^3–6^ to enable recognition of riboflavin derivatives of bacterial origin displayed by a monomorphic MHC class Ib molecule called MHC-related protein 1 (MR1) ^7–9^. Such derivatives are best exemplified by 5-(2-oxopropylideneamino)-6-D-ribitylaminouracil (5-OP-RU).

Viruses are devoid of the riboflavin synthesis machinery and consequently unable to activate MAIT cells in an MR1/*i*TCR-dependent manner. However, many viral infections lead to MAIT cell activation, primarily driven by inflammatory cytokines such as interleukin (IL)-7, -12, -15, -18 and type I interferons (IFNs) ^10–13^. The importance of MAIT cells in antiviral immune surveillance is evidenced by the heightened morbidity and mortality of influenza A virus (IAV)- infected *Mr1*^-/-^ mice that lack MAIT cells ^14^. In addition, a history of bacterial pneumonia after infection with varicella-zoster virus and persistent tattoo-associated human papilloma virus-positive warts was recently reported in an individual carrying a homozygous point mutation in *MR1*^15^. Finally, certain viruses impair MR1 expression or destroy MAIT cells as immune evasion tactics whose *in vivo* relevance remains to be better understood ^16–19^.

MAIT cells should be regarded as emergency responders to viral infections and their consequences and as pivotal players in vaccine-induced immunity. First, they are abundant in peripheral blood and enriched in mucosal tissues that provide entry points, transport systems or propagation sites for many viruses or vaccines. For instance, MAIT cells can comprise >2% of all T cells in the human lung where they safeguard against respiratory pathogens^20^, including but not limited to IAVs and severe acute respiratory syndrome coronavirus 2 (SARS-CoV-2) ^21,22^. Second, MAIT cells have a memory-like phenotype and poised effector functions, including rapid cytokine production ^23,24^. Once activated, they release T helper (T_H_)-1-, T_H_2- and/or T_H_17-type cytokines that transregulate the biological behaviors of myriad cell types, including dendritic cells (DCs) ^25^, natural killer (NK) cells ^25^, CD4^+^ and CD8^+^ conventional T (T_conv_) cells ^26,27^, and B cells^28–30^, all of which participate in antiviral defense. Third, MAIT cells constitutively express or acquire cytolytic effector molecules, including perforin (PFN), granzymes (GZMs) and granulysin, during viral infections, which can be deployed to eliminate MR1^+^ target cells and extracellular bacteria encountered during secondary infections ^2,31,32^. Furthermore, certain viral diseases and the cytokines they evoke (*e.g.*, hepatitis A and IL-15) upregulate natural killer group 2, member D (NKG2D) on MAIT cells, thus potentiating MR1-independent MAIT cell-mediated cytotoxicity against stressed or damaged target cells ^33^.

The inherent immunomodulatory capacity of MAIT cells has been demonstrated in the contexts of IAV infection and adenovirus-based immunization for coronavirus disease 2019 (COVID-19)^14,27^. However, whether MR1 ligands can be employed as vaccine adjuvants for viral diseases has not been explored. In this study, we used wild-type (WT) and MAIT cell-sufficient mouse strains, multiple vaccination models and routes, and a human cell culture system to demonstrate that 5-OP-RU can indeed serve as an efficacious adjuvant to enlarge tissue MAIT cell compartments, to generate MAIT cells with pro-inflammatory and immunoprotective properties, and to augment virus-specific T_conv_ cell responses.

## Materials and Methods

### Ethics statement

Animal experiments were performed following the Canadian Council on Animal Care guidelines and using animal use protocols (AUPs) approved by Animal Care and Veterinary Services at Western University (AUPs 2018-093, 2019-054 and 2020-084) or Animal Research Ethics Board at McMaster University (AUPs 19-12-32 and 21-04-12).

Human peripheral blood was collected from consenting healthy donors under protocol #5545, which was approved by the Western University Research Ethics Board for Health Sciences Research Involving Human Subjects.

### Mice

Adult WT C57BL/6 (B6), BALB/c and C3H/HeN mice were purchased from Charles River Canada (St. Constant, QC) and used at 6-14 weeks of age. Old WT B6 mice, which were provided by Dr. Tony Rupar (Western University), were housed in a pathogen-free barrier facility at Western University until they were used at 18-22 months of age. B6-MAIT^CAST^ mice ^34^ were provided by Dr. Olivier Lantz (Institut Curie, Paris, France) and bred in-house.

Adult B6(Cg)-*Ifnar1^tm1.2Ees^*/J, hereafter referred to as *Ifnar1*^-/-^ mice, were initially purchased from The Jackson Laboratory (Ann Arbor, Michigan) and bred at McMaster University. C57BL/6J-*Ptprc^em6Lutzy^*/J mice, hereafter referred to as CD45.1^+^ B6 mice, were also purchased from Jackson and used in the same facility.

Using sex- and age-matched animals was ensured in this study as appropriate.

### Viruses, vaccines and immunization routes

Several influenza A virus (IAV) strains, namely A/Puerto Rico/8/1934 (PR8) (H1N1), A/Northern Territory/60/1968 (NT60) (H3N2), A/Hong Kong/1/1968 (HK) (H3N2), and the X31 reassortant (H3N2), were propagated in embryonated chicken eggs before infectious allantoic fluid was harvested, filter-sterilized and stored at −80°C until use. X31 is a chimeric virus expressing the PR8 internal genes along with the hemagglutinin (HA) and neuraminidase (NA) of an H3N2 virus strain ^35^. Mice received an intraperitoneal (*i.p.*) inoculum approximating 600 hemagglutinating units of the above-indicated viruses. In several experiments, PR8 was inactivated at 56°C for 30 minutes before injection.

To study IAV-specific recall responses, we used a prime-boost immunization protocol in which B6-MAIT^CAST^ or B6 mice were injected *i.p.* with PR8 followed, 4 or 6 weeks later, by an *i.p.* injection of X31 as we previously described ^36^.

In a few experiments, BALB/c mice were inoculated intranasally (*i.n.*) with 20 μL of the 2021-2022 quadrivalent FluMist spray (AstraZeneca Canada Inc.), 10 μL in each nare, under light isoflurane-induced anesthesia. This vaccine contains live attenuated reassortants from A/Victoria/1/2020 (H1N1) [A/Victoria/2570/2019 (H1N1)pdm09-like virus], A/Tasmania/503/2020 (H3N2) [A/Cambodia/e0826360/2020 (H3N2)-like virus], B/Phuket/3073/2013 (Yamagata lineage), and B/Washington/02/2019 (Victoria lineage).

Separate cohorts of BALB/c mice were injected intramuscularly (*i.m.*) in the right hind limb with 5 × 10^8^ plaque-forming units (PFUs) of recombinant vesicular stomatitis viruses (rVSVs) expressing the Spike protein of the Wuhan strain of SARS-CoV-2 ^37^. Animals received either a single dose of the Indiana strain (rVSV_Ind_) or two-dose immunization with rVSV_Ind_ and the New Jersey strain (rVSV_NJ_) given 24 days apart.

To model antipoxviral immunization, B6 mice were injected *i.p.* with 10^6^ PFUs of the Western Reserve strain of vaccinia virus (VacV), which was propagated using the thymidine kinase-deficient osteosarcoma cell line 143B.

### In vivo treatments

To stimulate MAIT cells with an MR1 ligand, 5-OP-RU was administered *i.p.*, *i.m.*, or *i.n.* at estimated quantities of ∼10 nanomoles, ∼1 nanomole, or ∼76 picomoles per dose, respectively ^38,39^. To generate 5-OP-RU, 5-amino-6-D-ribitylaminouracil, which was generously provided by Dr. Olivier Lantz (Institut Curie), and methylglyoxal (Sigma) were mixed at equimolar concentrations in dimethyl sulfoxide (DMSO). After overnight incubation at room temperature, the mixture was stored at −80°C until use, at which point it was thawed and further diluted in sterile phosphate-buffered saline (PBS) for injection. Where indicated, parallel animal cohorts received the same amounts of methylglyoxal and DMSO in PBS as vehicle control.

To prevent tissue trafficking by immune cells, each mouse was injected twice *i.p.* with 1 mg/kg of the sphingosine-1-phosphate receptor 1 (S1PR1) antagonist FTY720 (Sigma) 2 hours before and 22 hours after a combination of PR8 and 5-OP-RU was administered ^40,41^. FTY720 was reconstituted in water and diluted in PBS before injection, and control animals received PBS only.

In a few experiments, mice were injected *i.p.* with 50 μg of the Toll-like receptor (TLR)3 agonist polyinosinic-polycytidylic acid [poly (I:C)] (InvivoGen), 50 μg of the TLR7 agonist imiquimod (InvivoGen), or 1 μg of recombinant mouse interferon-β (rmIFNβ) (R&D Systems), all in PBS.

To sufficiently block IFN a/β receptor subunit 1 (IFNAR-1) *in vivo*, multiple *i.p.* injections of an anti-IFNAR-1 monoclonal antibody (mAb) (clone MAR1-5A3) (Bio X Cell) were given. Accordingly, each mouse received 0.5 mg of this mAb 1 hour and 24 hours before immunization with PR8 and 5-OP-RU. This was followed by two additional doses, 0.25 mg each, of MAR1-5A3 at 24 and 48 hours post-immunization. Control cohorts received a mouse IgG1к isotype control (clone MOPC-21) from Bio X Cell.

### Viral challenge

B6 mice were injected *i.p.* with 5-OP-RU, vehicle, the HK virus, or a combination of HK and 5-OP-RU. After 3 days, under isoflurane anaesthesia, mice were challenged *i.n.* with 3 × 10^3^ PFUs of the mouse-adapted PR8 virus in 40 µL of saline. Animals were closely monitored for weight loss and mortality for the following 14 days.

### Tissue and specimen processing

At indicated endpoints, mice were euthanized by cervical dislocation. Animals were terminally bled by cardiac puncture using manually heparinized syringes with 26G needles. Peritoneal lavage was performed using 10-mL syringes with 18G needles and sterile PBS to obtain peritoneal fluid.

Spleens and livers were harvested aseptically before they were mechanically homogenized. Lungs were minced into small pieces, which were then subjected to enzymatic digestion for 1 hour at 37°C in RPMI-1640 containing 0.5 mg/mL of collagenase type IV (Sigma) and 25 µg/mL of DNase I (Sigma). Hepatic parenchymal cells were removed by density gradient centrifugation at 700 × *g* in 33.75% Percoll PLUS (GE Healthcare).

The resulting splenic, pulmonary and hepatic cell preparations were washed, filtered, exposed to ACK (Ammonium-Chlorine-Potassium) Lysing Buffer to eliminate erythrocytes, washed and filtered again. Trypan blue dye exclusion was used to ensure high cellular viability.

Non-terminal mouse bleeding was conducted via saphenous venipuncture, and isolated serum samples were frozen at −80°C until they were used for bead-based cytokine multiplexing by Eve Technologies (Calgary, AB).

To obtain human peripheral blood mononuclear cells (PBMCs), uncoagulated whole blood from male and female donors (age range: 21-66) was spun at 1,200 × *g* in 50-mL SepMate tubes (STEMCELL Technologies) containing Ficoll-Paque PLUS (Cytiva, Uppsala, Sweden).

### Quantitative polymerase chain reaction (PCR) analyses

To isolate pulmonary mouse MAIT cells, non-parenchymal lung mononuclear cells (LMNCs) from indicated B6-MAIT^CAST^ cohorts were subjected to magnetic cell sorting. Briefly, phycoerythrin (PE)-conjugated, 5-OP-RU-loaded mouse MR1 tetramers were used to stain MAIT cells ^7,42^, which were then purified using Anti-PE MicroBeads UltraPure, LS Columns and a QuadroMACS Separator (Miltenyi Biotec). Isolated MAIT cells were always 90-99% pure after two separation rounds. Cells were washed in PBS containing 10% BSA and 50 mM ethylenediaminetetraacetic acid before pellets were flash-frozen and stored at −80°C.

Total RNA was extracted using a PicoPure RNA Isolation Kit (Thermo Scientific) and converted to cDNA using SuperScript IV VILO Master Mix with ezDNase (Thermo Scientific). cDNA and TaqMan Fast Advanced Master Mix were added to each well of a custom-made, 96-well TaqMan Array Fast Plate (Thermo Scientific) containing lyophilized primer/probe sets listed in Supplementary Table 1. cDNA was amplified per manufacturer’s instructions, and cycle threshold (Ct) values were generated using a StepOne Plus Real-Time PCR System (Applied Biosystems). Normalized ΔCt values were determined by subtracting each Ct value by that of *Gapdh*, and the following formula was employed to calculate the relative mRNA content of MAIT cells for indicated genes: Fold Change = 2^-(ΔΔCt)^.

### Adoptive transfer experiments

LMNCs and hepatic MNCs (HMNCs) from CD45.2^+^ *Ifnar1*^-/-^ mice were isolated and pooled before ∼10^7^ cells were injected intravenously (*i.v.*), via tail vein, into CD45.1^+^ B6 mice. Twenty-four hours later, the recipients were injected *i.p.* either with PBS or with PR8 plus 5-OP-RU, followed 3 days later by cytofluorimetric enumeration of MAIT cells. The expression of the congenic marker CD45.1, or lack thereof, enabled us to distinguish between endogenous and transferred cells.

To investigate the proliferative capacity of mouse MAIT cells *in vivo*, we isolated naïve B6-MAIT^CAST^ LMNCs and HMNCs, which were immediately pooled and subjected to magnetic MAIT cell sorting. Isolated MAIT cells were then expanded *ex vivo* following a recently published protocol by Parrot *et al.* ^43^. MAIT cell-depleted fraction was irradiated at 35 Gy and used as a source of feeder cells, which were seeded at 2 × 10^5^ cells/well along with 2 × 10^4^ MAIT cells/well of a U-bottom plate. Cells were maintained in ImmunoCult T Cell Expansion Medium (STEMCELL Technologies) supplemented with 20 ng/mL of recombinant mouse interleukin (IL)- 2 (R&D systems), 8% CTS Immune Cell Serum Replacement (Thermo Fisher), 100 µg/mL of Normocin (InvivoGen), and 100 U/mL Penicillin/Streptomycin. Cultures were split twice a week and replenished with fresh medium and feeder cells. After 3 weeks, expanded MAIT cells were washed, labeled with 1 µM CellTrace Far Red dye (Thermo Fisher), and injected *i.v.* at 5 × 10^5^ cells/B6-MAIT^CAST^ mouse. Twenty-four hours after adoptive transfer, the recipients were left untreated or given PR8 and 5-OP-RU *i.p.* before they were sacrificed, 3 days later, for several organs in which MAIT cell proliferation was assessed by flow cytometry.

### Ex vivo cell stimulation protocols

Mouse splenocytes, peritoneal cells, LMNCs and/or HMNCs were resuspended in RPMI-1640 supplemented with 10% heat-inactivated fetal bovine serum (FBS), 2 mM GlutaMAX-I, 0.1 mM MEM nonessential amino acids, 1 mM sodium pyruvate, 10 mM HEPES, 100 U/mL penicillin and 100 µg/mL streptomycin, which is referred to as complete medium in this work.

Up to 2 × 10^6^ MNCs were seeded into each well of U-bottom plates. Cells were left untreated in complete medium, stimulated for 4 hours with 50 ng/mL of phorbol 12-myristate 13-acetate (PMA) plus 500 ng/mL of ionomycin, or stimulated for 24 hours with 5 ng/mL of recombinant mouse IL-12p70 (Peprotech) plus 5 ng/mL of recombinant mouse IL-18 (R&D Systems). To retain intracellular mediators for downstream cytofluorimetric analyses, 10 µg/mL of brefeldin A (Sigma) was added at the beginning of cultures containing PMA and ionomycin, or after 18 hours of stimulation with IL-12 and IL-18. Cultures were incubated inside a humidified incubator set at 37°C and 5% CO_2_.

In several experiments, MAIT cells were magnetically purified before stimulation. They were rested for 2 hours at 37°C before they were washed, seeded at ∼120,000 cells/well, and stimulated with PMA and ionomycin or with IL-12 and IL-18 as described above. In other experiments, MAIT cells were stimulated with plate-coated anti-CD3 (clone 17A2) (Bio X Cell) in the presence of 2 µg/mL of an anti-CD28 mAb (clone 37.51) (Bio X Cell). After 24 hours, culture supernatants were collected, aliquoted and stored at −80°C until their cytokine contents were quantified using Ready-SET-Go! ELISA Kits from eBioscience.

Human PBMCs were seeded at 5 × 10^5^ cells/well in U-bottom microplates in the absence or presence of C1R-MR1 cells that had been left unmanipulated, infected with PR8 and/or pulsed with 2 nM 5-OP-RU. The MR1-overexpresing lymphoblastoid human B cell line C1R-MR1 was provided by Dr. Jose Villadangos (University of Melbourne). To infect C1R-MR1 cells, 10^5^ TCID_50_ (50% tissue culture infectious dose) of PR8 was added to 10^7^ cells/mL in Opti-MEM I Reduced Serum Medium (Thermo Fisher). After 1 hour at 37°C, infected cells were washed, resuspended in complete medium, and co-incubated at a 1:2 ratio with PBMCs for 16, 48 or 96 hours prior to cytofluorimetric analyses of indicated T cell surface markers.

In additional experiments, 10^7^ PBMCs/mL were incubated with 5 µM carboxyfluorescein diacetate succinimidyl ester (CFDA-SE) (Invitrogen) for 20 minutes at 37°C. Labeled cells were then washed, resuspended in complete medium and seeded at 10^6^ cells/well in U-bottom plates either alone or in the presence of unmanipulated, PR8-infected, 5-OP-RU-pulsed, or PR8-infected and 5-OP-RU-pulsed C1R-MR1 cells. Four or 7 days later, CFDA-SE dye dilution was assessed by flow cytometry as a measure of cellular proliferation.

### Flow cytometry

Mouse cells were incubated for 15 minutes on ice with 2.4G2 B cell hybridoma culture supernatant containing a CD16/CD32-blocking mAb. This was to prevent false positive staining due to non-specific binding of fluorochrome-conjugated mAbs to Fcy receptors.

5-OP-RU-loaded MR1 tetramers and PBS-57-loaded CD1d tetramers were provided by the NIH Tetramer Core Facility (Atlanta, GA) for MAIT and invariant natural killer T (*i*NKT) cell staining, respectively. As negative staining controls, 6-formylpterin (6-FP)-loaded MR1 tetramers and empty CD1d tetramers, also supplied by the NIH Tetramer Core Facility, were utilized. B220^+^ events were excluded to reduce background noise during mouse MAIT cell staining.

Fluorochrome-conjugated mAbs against various cell surface markers and corresponding isotype controls are listed in Supplementary Table 2. These reagents were prepared in a staining buffer containing 2% FBS in PBS and added to cell preparations for 20 minutes at room temperature. To detect intracellular mediators, we used the Foxp3/Transcription Factor Staining Buffer Set from Thermo Fisher. After fixation and permeabilization for 20 minutes at room temperature, samples were incubated with indicated mAbs or isotype controls in permeabilization buffer for 30 minutes at room temperature in the dark. Staining with 7-aminoactinomycin D (7-AAD) or Fixable Viability Dye (eBioscience) allowed for dead cell exclusion in our analyses.

To detect IAV-specific CD8^+^ T cells on day 8 post-FluMist instillation, ∼10^6^ LMNCs and 100 µL of whole blood from vaccinated BALB/c mice were stained with an anti-CD8a mAb (clone 53-6.7) and Alexa 488-labeled H-2K^d^:TYQRTRALV tetramers (NIH Tetramer Core Facility). TYQRTRALV is a linear peptide corresponding to the immunodominant peptide epitope of the IAV nucleoprotein (NP), namely NP_147-155_, in the BALB/c strain ^44^. Erythrocytes were removed from the whole blood specimen after cell surface staining.

NP_147-155_-specofic T cells were also enumerated using intracellular cytokine staining (ICS) for IFN-γ as we previously described ^45^. In brief, 2 × 10^6^ LMNCs from vaccinated mice were left untreated, exposed to 500 nM of an H-2^d^-restricted irrelevant peptide derived from the lymphocytic choriomeningitis virus (LCMV) NP (NP_112-126_: RPQASGVYM) ^46^, or stimulated with 500 nM of TYQRTRALV for 5 hours at 37°C. Brefeldin A was present at 10 µg/mL during the final 3 hours, after which cells were washed, stained with anti-CD8α, fixed and permeabilized, incubated with an anti-IFN-γ mAb (clone XMG1.2), washed again and analyzed.

A similar ICS protocol was used to detect splenic and pulmonary CD8^+^ T cells recognizing an immunodominant epitope of the SARS-CoV-2 Spike protein, namely S_535-543_ (KNKCVNFNF) ^47^, in vaccinated BALB/c mice.

Stained cells were interrogated using a BD FACSCanto II flow cytometer, and data were analyzed using FlowJo software version 10.0.7 (Tree Star).

### Statistical analyses

Statistical comparisons were carried out using GraphPad Prism 8.0.1 software. Chi-squared tests, Student’s *t*-test, or ANOVA with relevant post-hoc tests were employed as appropriate, and differences with *p* ≤ 0.05 were considered significant. Sample sizes and the statistical methods used are detailed within figure legends.

## Results

### 5-OP-RU synergizes with a PR8-based vaccine to induce robust MAIT cell accumulation in multiple tissues

MAIT cells can be activated by viral infections or in response to anti-pathogen vaccines ^22,27^. However, whether MR1 ligands, typified by 5-OP-RU, optimize the efficacy of antiviral immunization strategies has not been explored. We began to address this question following a serendipitous observation in a mouse model of IAV vaccination in which PR8 was inoculated *i.p.* into MAIT cell-sufficient mice.

Unlike in humans, MAIT cells are scarce in conventional strains of laboratory mice. Therefore, we used a congenic strain, namely B6-MAIT^CAST^, which harbors ∼20 times more MAIT cells compared with WT B6 mice ^34^ and, as such, phenocopy human MAIT cell repertoires.

MAIT cells comprised only a tiny fraction of peritoneal TCRβ^+^ cells in naïve B6-MAIT^CAST^ mice or in animals that had received either PR8 or 5-OP-RU alone 3 days earlier (Fig. 1A-D). However, combining PR8 and 5-OP-RU resulted in massive MAIT cell accumulation inside the peritoneal cavity (Fig. 1B-D). This was not a transient response since both the frequency and the absolute number of peritoneal MAIT cells remined high on day 7 (Supplementary Fig. 1).

**Fig. 1:**
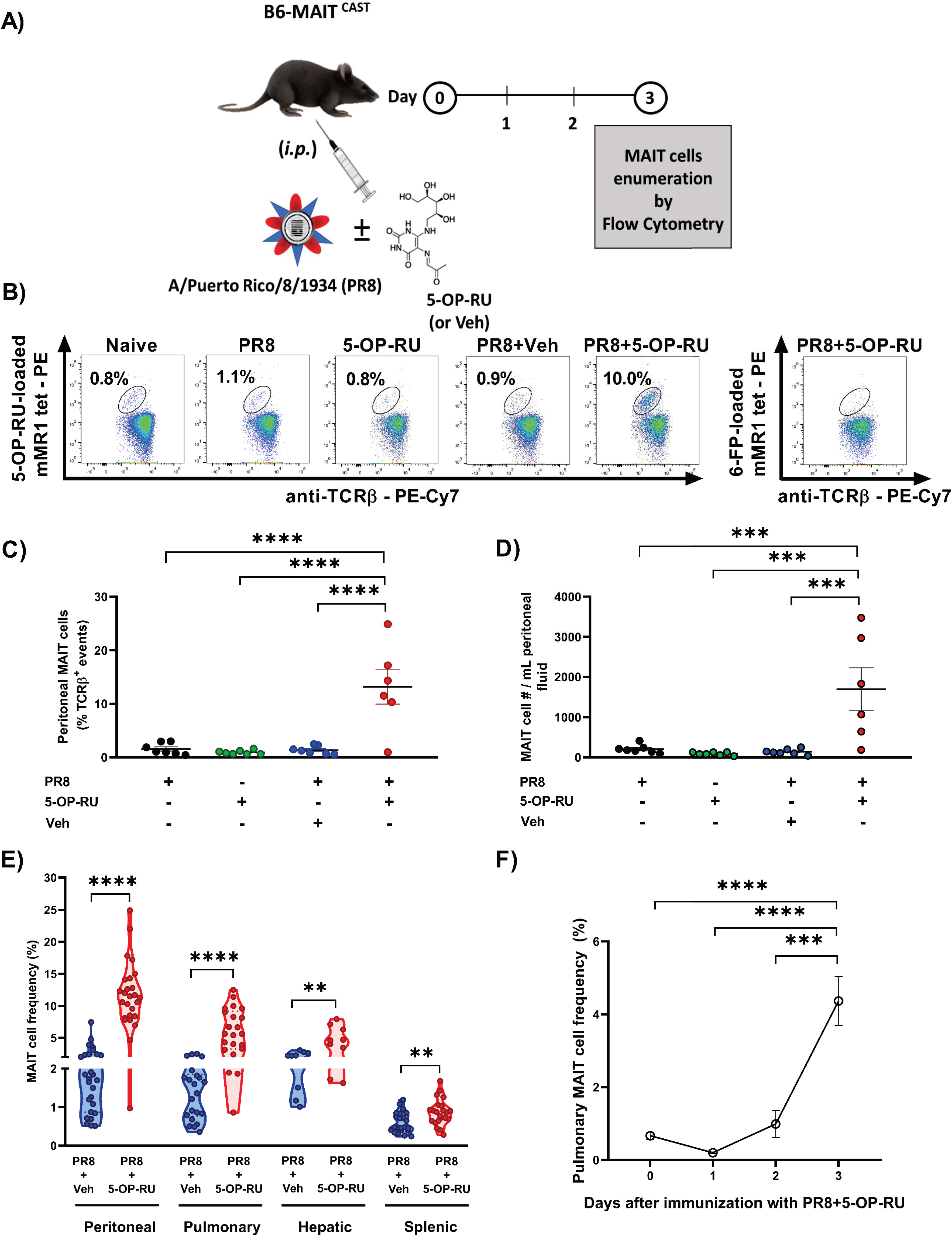
Intraperitoneal co-administration of PR8 and 5-OP-RU results in MAIT cell accumulation in multiple tissues. B6-MAIT^CAST^ mice were left unimmunized or inoculated *i.p.* with PR8, 5-OP-RU/vehicle, or both followed, 3 days later (**A-E**) or at indicated time points (**F**), by cytofluorimetric enumeration of MAIT cells within the peritoneal cavity (**B-E**), lungs (**E-F**), liver and spleen (**E**). 6-FP-loaded mouse MR1 tetramers were used as a staining control (**B**). Peritoneal MAIT cell frequencies (**B-C**) and their absolute numbers (**D**) are illustrated. Each circle (**C-E**) represents an individual mouse. For the kinetics of the pulmonary MAIT cell response to PR8 plus 5-OP-RU, data from day 0 (n=5), day 1 (n=3), day 2 (n=5) and day 3 (n=5) were analyzed, with error bars representing the standard error of the mean (SEM) (**F**). One-way ANOVA was followed by Dunnett’s post-hoc multiple comparisons (**C, D and F**). For group comparisons, Mann-Whitney *U* test and unpaired *t*-tests were employed as appropriate (**E**). **, *** and **** denote significant differences with *p* ≤ 0.01, *p* ≤ 0.001 and *p* ≤ 0.0001, respectively.

Importantly, *i.p.* inoculation of 5-OP-RU plus PR8 enlarged the MAIT cell compartment not only in the peritoneal cavity but also in the lungs, liver and spleen (Fig. 1E). The pulmonary response kinetics pointed to early MAIT cell activation since these cells were less detectable by MR1 tetramer staining on day 1 (Fig. 1F), due likely to activation-induced invariant TCR (*i*TCR) internalization.

We confirmed the specificity of 5-OP-RU, as an adjuvant candidate, for MAIT cells by enumerating other T cell types, including *i*NKT cells, a prominent invariant T cell type in rodents. As expected, pulmonary, hepatic and splenic *i*NKT cell frequencies remained unaltered post-vaccination (Supplementary Fig. 2).

### Incorporating 5-OP-RU in secondary IAV immunization leads to increased MAIT cell numbers

To test the efficacy of 5-OP-RU in a heterosubtypic prime-boost immunization model mimicking annual IAV booster vaccinations, we inoculated B6-MAIT^CAST^ mice *i.p.* with PR8 (H1N1) followed, 4 weeks later, by an *i.p.* injection of the reassortant H3N2 virus X31 ^35,36^. This strategy prevents the Abs generated through PR8 priming from quickly removing the boosting X31 virions from the system. As with the primary response, the enhancing effect of 5-OP-RU was evident when this compound was added to X31 (Fig. 2). Pulmonary MAIT cell numbers were also elevated 3 days after immunization with X31 and 5-OP-RU when WT B6 mice were used in lieu of B6-MAIT^CAST^ mice (Supplementary Fig. 3).

**Fig. 2:**
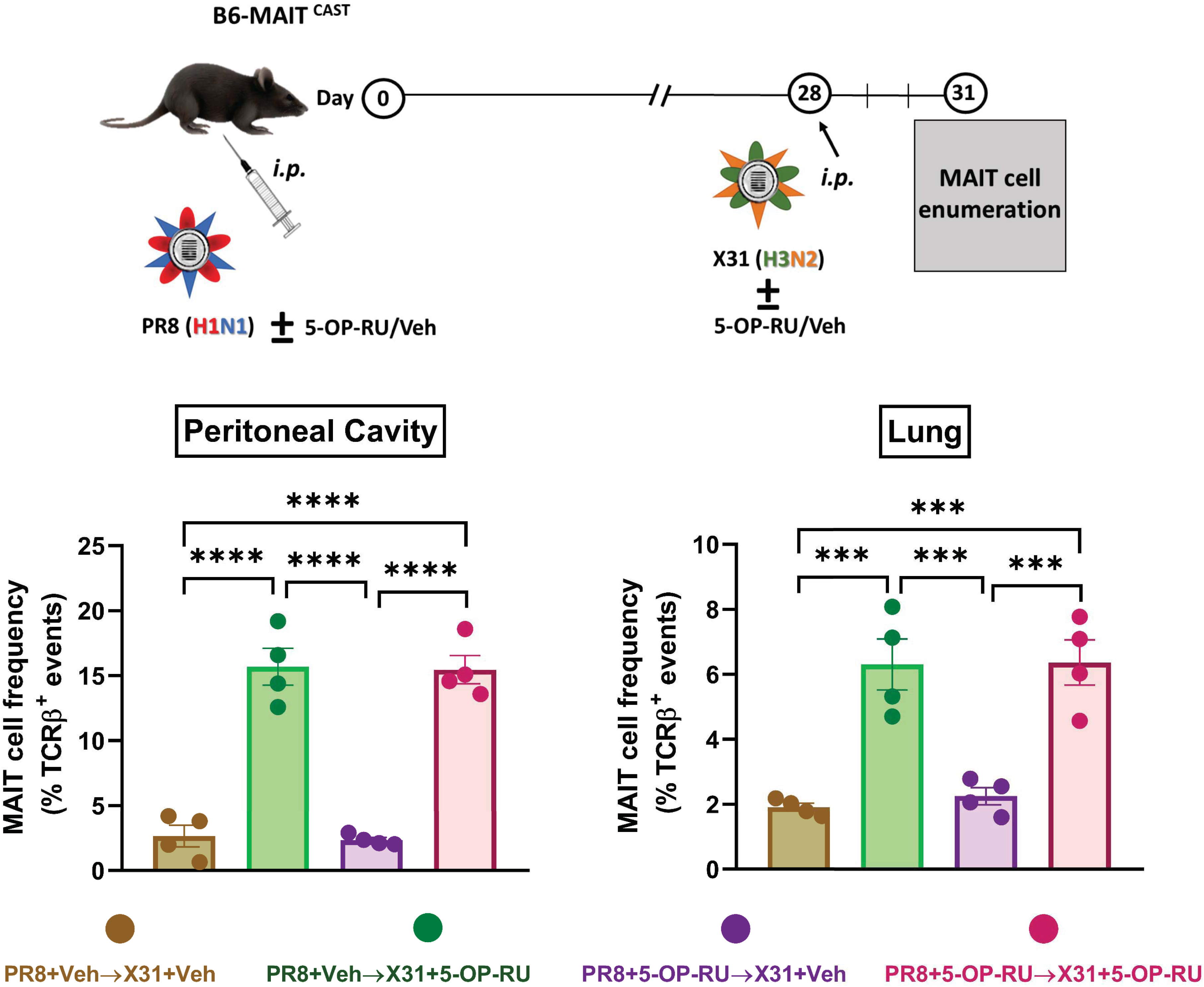
5-OP-RU synergizes with IAV in a prime-boost immunization model. B6-MAIT^CAST^ mice (n=4/cohort) were primed *i.p.* with PR8 (H1N1) plus 5-OP-RU/vehicle four weeks before they received a boosting injection of the reassortant X31 virus (H3N2) with 5-OP-RU/vehicle. Three days later, peritoneal and pulmonary MAIT cell frequencies were determined by flow cytometry. Each circle corresponds to an individual mouse, and error bars represent the SEM values. Statistical analyses were performed using one-way ANOVA followed by Tukey’s Multiple Comparisons tests. *** and **** denotes significant differences with *p* ≤ 0.001 and *p* ≤ 0.0001, respectively.

We next asked whether including 5-OP-RU in both primary and secondary immunizations further increases MAIT cell frequencies. This was clearly not the case since peritoneal and pulmonary MAIT cell percentages were similar in animals receiving 5-OP-RU during both phases and those receiving 5-OP-RU in the boosting phase only (Fig. 2). A similar pattern emerged when hepatic and splenic MAIT cells were enumerated (Supplementary Fig. 4A-B). This experiment also demonstrates that a prior exposure to 5-OP-RU does not induce MAIT cell anergy.

### 5-OP-RU-induced MAIT cell accumulation is observed in both sexes, in young and old mice, and across multiple genetic strains and vaccines

To investigate whether the enhancing effect of 5-OP-RU is manifest in both sexes, we retrospectively determined MAIT cell frequencies in closely age-matched female and male mice receiving PR8 in combination with either 5-OP-RU or vehicle. These analyses confirmed that both sexes mount similarly rigorous responses to a 5-OP-RU-adjuvanted PR8 vaccine (Fig. 3A).

**Fig. 3:**
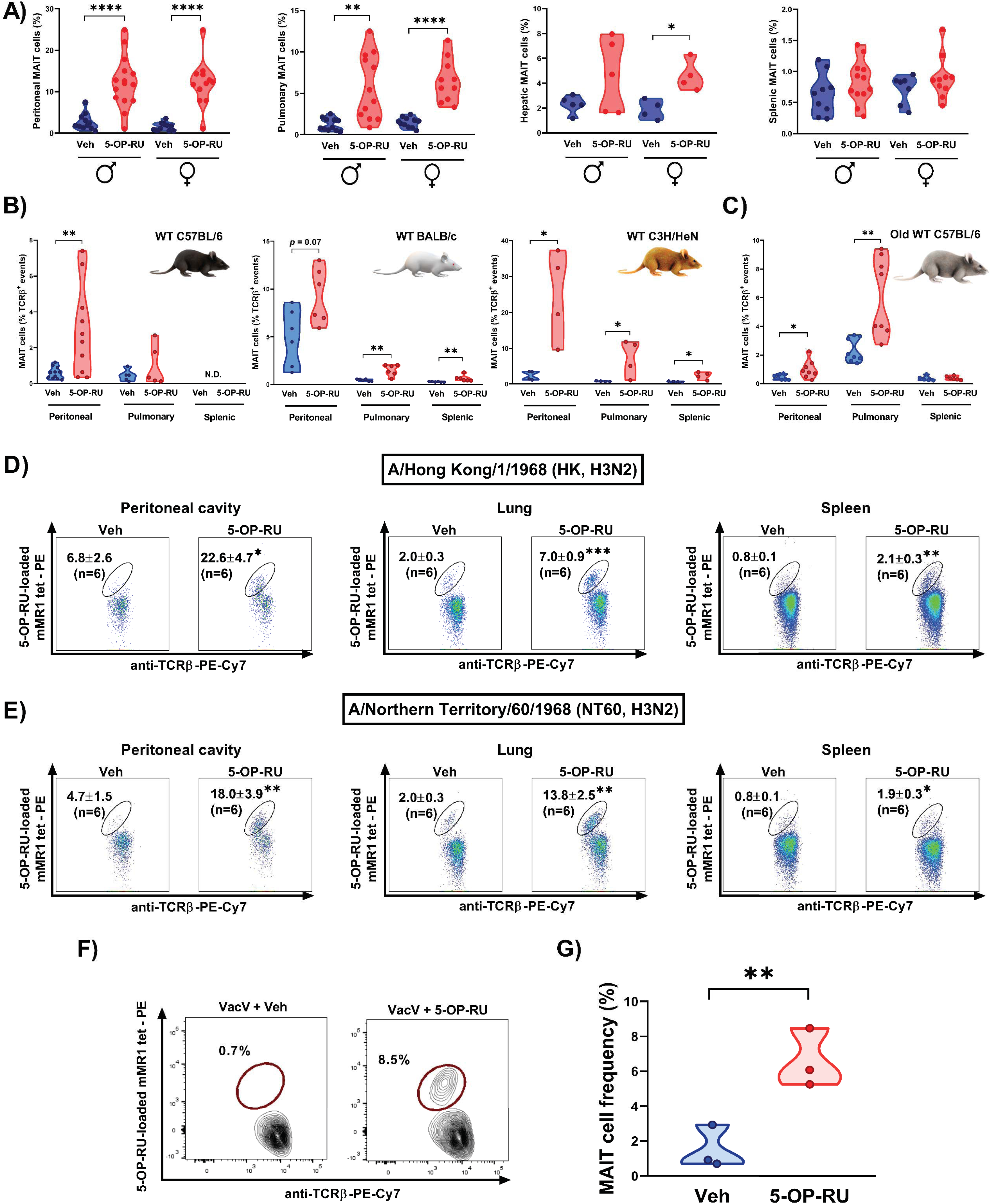
The adjuvanticity of 5-OP-RU is manifest in both sexes, in young and old animals, and in different mouse strains receiving immunization with various IAV strains or VacV. Age-matched male (♂) and female (♀) B6-MAIT^CAST^ mice (**A**), sex-matched young (6-14-week-old) and old (18-22-month-old) wild-type C57BL/6 (H-2^b^) mice (**B-C**), and age/sex-matched wild-type BALB/c (H-2^d^) and C3H/HeN (H-2^k^) mice (**B**) were inoculated *i.p.* with PR8 and 5-OP-RU (or vehicle). Separate cohorts of B6-MAIT^CAST^ mice were immunized *i.p.* with the H3N2 IAV strains HK (**D**) or NT60 (**E**) or with vaccinia virus (VacV) (**F-G**) in combination with either 5-OP-RU or vehicle as indicated. MAIT cell frequencies in specified sites were determined by flow cytometry on day 3 post-immunization. Each circle corresponds to an individual mouse in violin plots (**A-C and G**). Representative plots are also depicted (**D-F)** with mean ± SEM values indicated (**D-E**). Unpaired *t*-tests were used for group comparisons. *, **, *** and **** denote significant differences with *p* ≤ 0.05, *p* ≤ 0.01, *p* ≤ 0.001 and *p* ≤ 0.0001, respectively.

In the next series of experiments, we tested the efficacy of 5-OP-RU in several WT mouse strains, including B6 (H-2^b^), BALB/c (H-2^d^) and C3H/HeN (H-2^k^). Although MAIT cells are rare in these animals, *i.p.* co-administration of PR8 and 5-OP-RU made tissue MAIT cells readily detectable and raised their frequencies (Fig. 3B).

MAIT cells undergo a numerical decline with age ^48,49^, and an optimal vaccine should work in both young and old individuals. We found the combination of 5-OP-RU and PR8 to induce a MAIT cell surge not only in 6-14-week-old young animals but also in 18-22-month-old B6 mice (Fig. 3C).

Finally, we sought to explore the adjuvanticity of 5-OP-RU when used in conjunction with IAV strains other than the mouse-adapted PR8 strain (H1N1) or even non-IAV vaccines. We found a dramatic rise in MAIT cell numbers following *i.p.* immunization with two non-mouse-adapted H3N2 strains, namely HK (Fig. 3D) and NT60 (Fig. 3E). Also interestingly, this phenomenon was reproducible when we used VacV (Fig. 3F-G), a DNA virus with a radically different genomic structure and tissue tropism compared with IAVs. Immunization with VacV prevents smallpox and is also thought to elicit cross-protective responses against monkeypox ^50^.

### In vivo proliferation of MAIT cells accounts for their tissue accumulation after 5-OP-RU-adjuvanted anti-IAV vaccination

To begin to address the observed phenotype mechanistically, we asked whether MAIT cell accumulation was a result of their enhanced proliferative capacity or altered migratory behavior. We took several approaches to answer this question. First, we examined MAIT cells from B6-MAIT^CAST^ mice for their expression of CD69, an early activation marker that mediates tissue retention by suppressing S1PR1 ^51,52^. Animals that had received PR8 plus 5-OP-RU showed a gradual rise in both the proportion of CD69^+^ MAIT cells and the geometric mean fluorescence intensity (gMFI) of CD69 staining, indicating the higher expression level of this molecule on a per-cell basis (Fig. 4A and Supplementary Fig. 5). Second, treating mice with FTY720 (*aka.*, fingolimod), a drug that antagonizes S1PR1 to block lymphocyte trafficking^40,41^, failed to reverse MAIT cell accumulation (Fig. 4B). This was not due to absent S1PR1 expression by MAIT cells. In fact, MAIT cells from different tissues displayed stable or enhanced S1PR1 levels compared with non-MAIT T lymphocytes that consist primarily of T_conv_ cells (Fig. 4C). Third, we found vaccination with PR8 plus 5-OP-RU to upregulate Ki-67, a proliferation marker, on MAIT cells (Fig. 4D). Finally, to evaluate MAIT cell expansion more definitively, we adoptively transferred magnetically purified, *ex vivo*-expanded, CellTrace-labeled MAIT cells into B6-MAIT^CAST^ mice, which were then left unvaccinated or inoculated with PR8 plus 5-OP-RU. Three days later, substantial CellTrace dye dilution among pulmonary, hepatic and splenic MAIT cells was detectable in vaccinated animals only, indicative of *in vivo* MAIT cell proliferation (Fig. 4E).

**Fig. 4:**
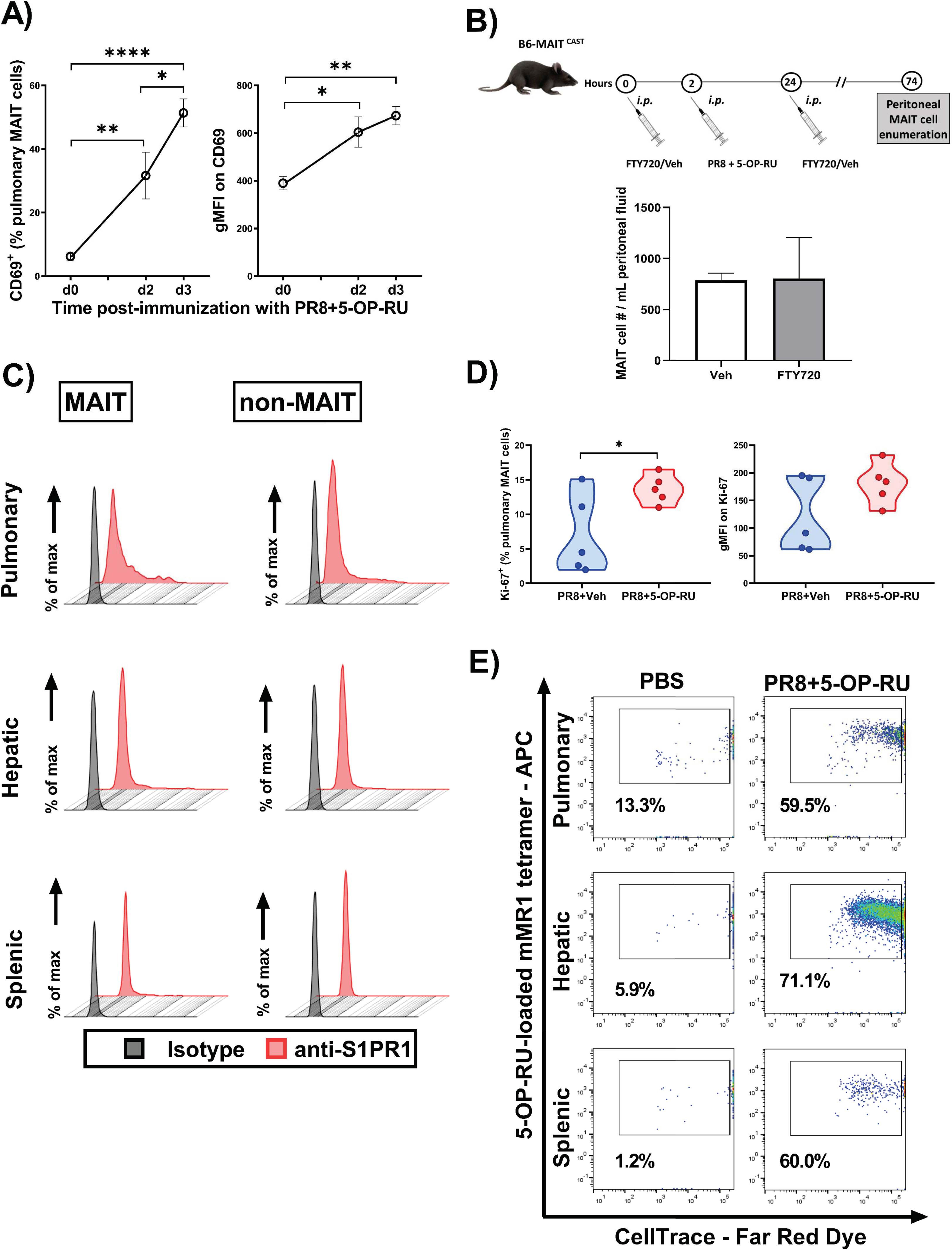
Immunization with 5-OP-RU-adjuvanted PR8 leads to CD69 and Ki-67 upregulation by MAIT cells and their *in vivo* proliferation but not migration. B6-MAIT^CAST^ mice (n=5/group) were injected *i.p.* with PR8 plus 5-OP-RU (**A** and **D**) or vehicle (**D**) 3 days before pulmonary MAIT cells were analyzed for CD69^+^ (**A**) and Ki-67^+^ (**D**) cell frequencies and for the geometric mean fluorescence intensity (gMFI) of staining for these markers (**A** and **D**). Separate cohorts (n=3/group) received *i.p.* injections of FTY720 (or vehicle) 2 hours before and 22 hours after immunization with PR8 and 5-OP-RU, and peritoneal MAIT cells were enumerated on day 3 post-immunization (**B**). The surface expression levels of S1PR1 on TCRβ^+^ MR1 tetramer^+^ tissue MAIT cells and TCRβ^+^ MR1 tetramer^-^ non-MAIT T cells were also evaluated in naïve B6-MAIT^CAST^ mice (n=2) (**C**). Representative plots from one mouse are depicted (**C**). To examine MAIT cell proliferation *in vivo*, CellTrace-labeled purified MAIT cells were adoptively transferred into B6-MAIT^CAST^ mice, which subsequently received either PBS or a combination of PR8 and 5-OP-RU *i.p.* (**E**). Three days later, CellTrace dye dilution by MAIT cells from indicated sites was analyzed by flow cytometry (**E**). Statistical differences were computed using one-way ANOVA followed by Tukey’s Multiple Comparisons test (**A**), Mann-Whitney *U* test (**B**) and unpaired *t*-test (**D**). *, ** and **** denote differences with *p* ≤ 0.05, *p* ≤ 0.01 and *p* ≤ 0.0001, respectively.

Taken together, the above results demonstrate that MAIT cell accumulation following 5-OP-RU-adjuvanted IAV immunization stems from the *in situ* expansion of these cells in indicated tissues as opposed to their recruitment from other locations.

### The adjuvant effect of 5-OP-RU depends on vaccine replication, TLR3 engagement and in vivo IFNAR signaling

To determine whether active viral propagation was required for the adjuvant effect of 5-OP-RU, we compared intact PR8 and heat-inactivated PR8 (HI-PR8). As illustrated in Fig. 5A, HI-PR8 failed to synergize with 5-OP-RU to make MAIT cells proliferate. This finding, coupled with our observation that fundamentally different viruses (*i.e.*, IAVs and VacV) could be combined with 5-OP-RU to induce MAIT cell expansion (Fig. 1 and Fig. 3D-G), suggested a role for double-stranded ribonucleic acid (dsRNA) in our model.

**Fig. 5:**
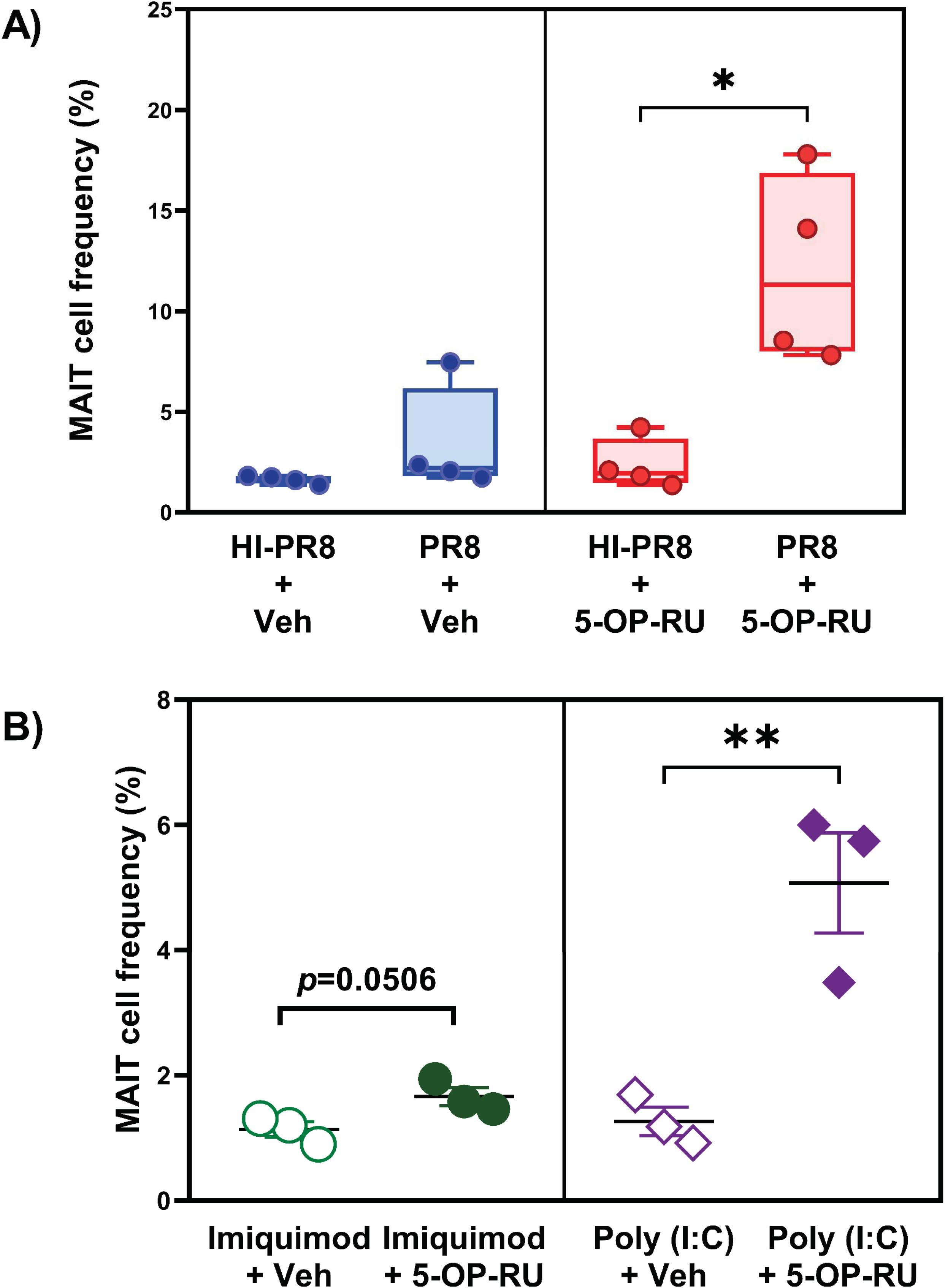
5-OP-RU can be combined with poly (I:C), but not with heat-inactivated PR8, to expend the peritoneal MAIT cell population. B6-MAIT^CAST^ mice (n=4/group) were inoculated *i.p.* with intact or heat-inactivated PR8 (HI-PR8) in combination with 5-OP-RU or vehicle as indicated (**A**). Additional cohorts (n=3/group) were injected *i.p.* with poly (I:C) or imiquimod plus 5-OP-RU or vehicle (**B**). Three days later, peritoneal MAIT cell frequencies were determined by flow cytometry. Each circle or diamond represents an individual mouse. Statistical comparisons were made using unpaired *t*-tests, with significant differences identified.

The presence of dsRNA, a by-product of viral replication, can be sensed by TLR3, resulting in a type I IFN response ^53^. We found poly (I:C), a synthetic analog of viral dsRNA and a TLR3 agonist, to expand MAIT cells when co-administered with 5-OP-RU (Fig. 5B), thus partially simulating the effect of the PR8 virions (Supplementary Fig. 4). In contrast, using imiquimod to trigger TLR7, a major IAV recognition receptor ^54,55^, did not markedly increase the frequency of MAIT cells (Fig. 5B and Supplementary Fig. 6).

Among detectable cytokines in the sera of vaccinated B6-MAIT^CAST^ mice (Supplementary Fig. 7), we focused on IL-7, IL-12, IL-15, IL-18 and type I IFNs, which are known to stimulate MAIT cells^11,22^. These analyses revealed a noticeable rise in IFN-β levels in animals receiving PR8 plus 5-OP-RU at the six-hour timepoint (Fig. 6A). Given the importance of the TLR3-IFN axis in antiviral responses, we sought to investigate the role of type I IFNs in our model. First, we injected mice with 5-OP-RU, rmIFNβ, or both before enumerating pulmonary MAIT cells. We found rmIFNβ administration to raise the frequency of MAIT cells only if it was combined with 5-OP-RU (Fig. 6B). This was a robust response although it did not reach the same magnitude as that achieved by a combination of 5-OP-RU and PR8 at least on day 3 (Fig. 6B). In subsequent experiments, mAb-mediated blockade of IFNAR-1 prevented tissue MAIT cell accumulation (Fig. 6C). Finally, to ascertain whether IFNAR signaling was required in a MAIT cell-intrinsic or - extrinsic fashion, we transferred pooled LMNCs and HMNCs from CD45.2^+^ *Ifnar1*^-/-^ mice *i.v.* into WT B6 mice expressing the congenic marker CD45.1, which were subsequently injected with PBS or with a combination of PR8 and 5-OP-RU (Fig. 6D). As anticipated, endogenous CD45.1^+^ MAIT cells underwent expansion in vaccinated animals. In stark contrast, there were very few detectable MAIT cells among *Ifnar1*^-/-^ (CD45.2^+^) cells (Fig. 6D), which contained a MAIT cell fraction before they were transferred (∼0.5% of LMNCs and ∼1.75% of HMNCs). Therefore, MAIT cell-autonomous IFNAR signaling is required for the expansion of these cells following vaccination with PR8 and 5-OP-RU.

**Fig. 6:**
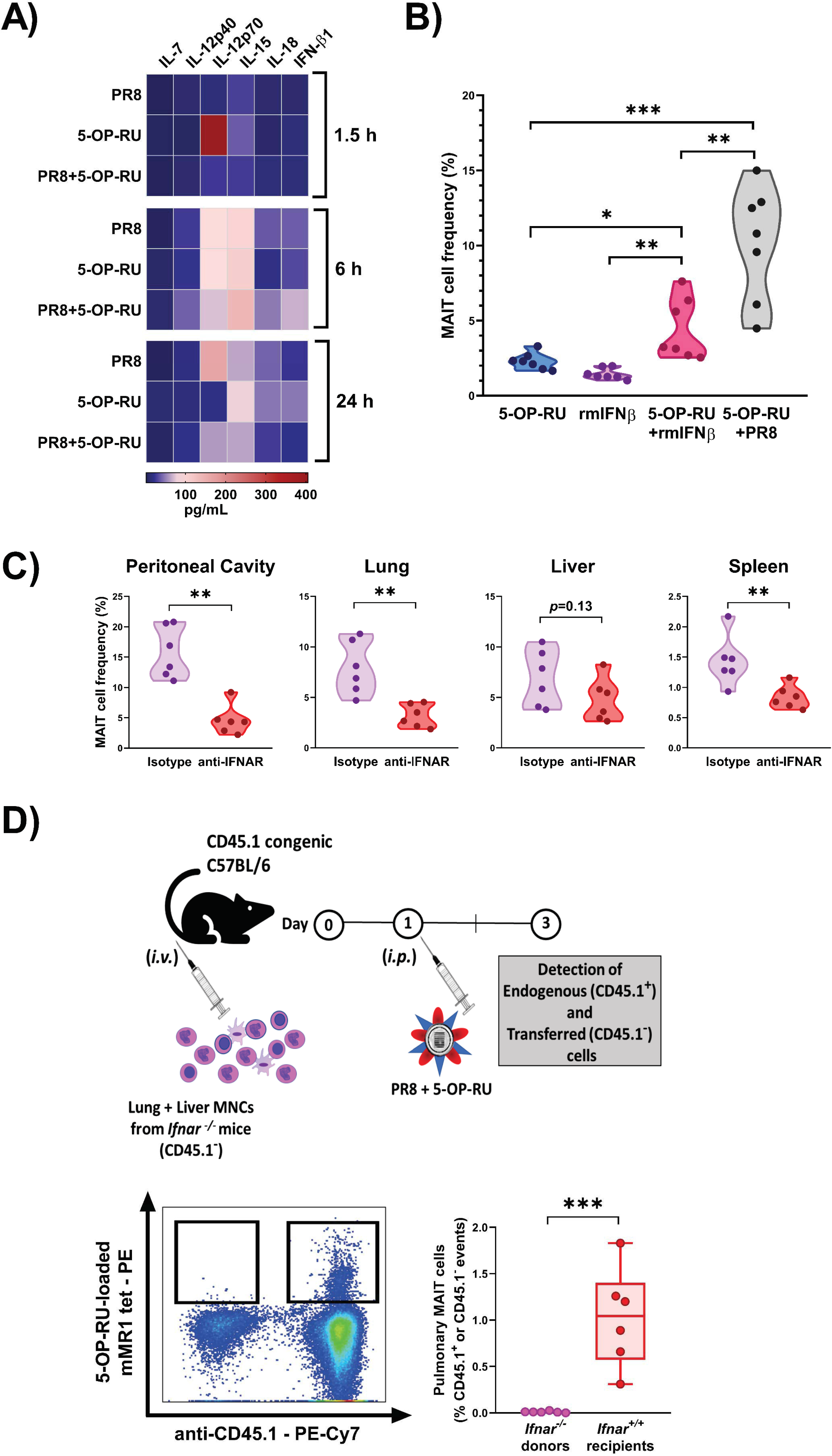
MAIT cell-intrinsic IFNAR signaling is required for MAIT cell proliferation after immunization with PR8 plus 5-OP-RU. (**A**) B6-MAIT^CAST^ mice were injected *i.p.* with PR8 (n=12), 5-OP-RU (n=12), or both (n=12). Animals (n=4/timepoint) were sacrificed 1.5, 6 or 24 hours later, and indicated cytokines were quantified in serum samples. A heatmap was generated to visualize average values at specified points. (**B**) In separate experiments, B6-MAIT^CAST^ mice (n=7/group) were injected *i.p.* with 5-OP-RU, recombinant mouse IFN-β (rmIFNβ), 5-OP-RU plus rmIFNβ, or 5-OP-RU plus PR8. (**C**) Additional cohorts (n=6/group) received an IFNAR-blocking monoclonal antibody or isotype control twice before and twice after *i.p.* immunization with PR8 and 5-OP-RU as detailed in Materials and Methods. Pulmonary (**B-C**) and peritoneal, hepatic and splenic (**C**) MAIT cells were enumerated by flow cytometry on day 3 post-immunization. (**D**) To study IFNAR signaling requirements, non-parenchymal pulmonary and hepatic mononuclear cells (MNCs) were pooled from CD45.1^-^ *Ifnar1*^-/-^ mice (n=9) and injected *i.v.* (at ∼10^7^ transferred cells/recipient) into congenic CD45.1^+^ B6 mice (n=6). Recipients were inoculated *i.p.* with PR8 plus 5-OP-RU three days before CD45.1^+^ (endogenous) and CD45.1^-^ (transferred) cells were analyzed in the lungs for their pulmonary MAIT cell component. A representative cytofluorimetric plot and summary data are depicted. Unpaired *t*-tests were employed for statistical comparisons. *, ** and *** denote differences with *p* ≤ 0.05, *p* ≤ 0.01 and *p* ≤ 0.001, respectively.

### Immunization with PR8 plus 5-OP-RU promotes the MAIT1 program

Most MAIT cells in WT B6 and B6-MAIT^CAST^ mice express retinoic acid receptor-related orphan receptor γt (RORγT) consistent with a MAIT17 phenotype ^56^. There also exists a smaller MAIT1 population expressing T-box expressed in T cells (T-bet). These transcription factors govern MAIT cells’ immunomodulatory activities and inflammatory cytokine profiles. Therefore, we were curious to know whether vaccination with PR8 and 5-OP-RU reprograms MAIT cells.

In immunophenotyping experiments, we found the vast majority of expanded MAIT cells to be low CD103 (Supplementary Fig. 8) and high CD122 expressors (Supplementary Fig. 9), suggesting a functional MAIT1 lineage bias ^57,58^. Moreover, apart from their increased *Il2rb* transcript levels, the expanded population showed diminished *Il23r* and *Ccr6* (Supplementary Fig. 10 and Supplementary Fig. 11) and stable *Icos* expression (Supplementary Fig. 12), all of which can be viewed as MAIT1 characteristics ^57,58^. In fact, *Tbx21*, which encodes T-bet, was strongly upregulated in MAIT cells following vaccination with PR8 plus 5-OP-RU (Fig. 7A).

**Fig. 7:**
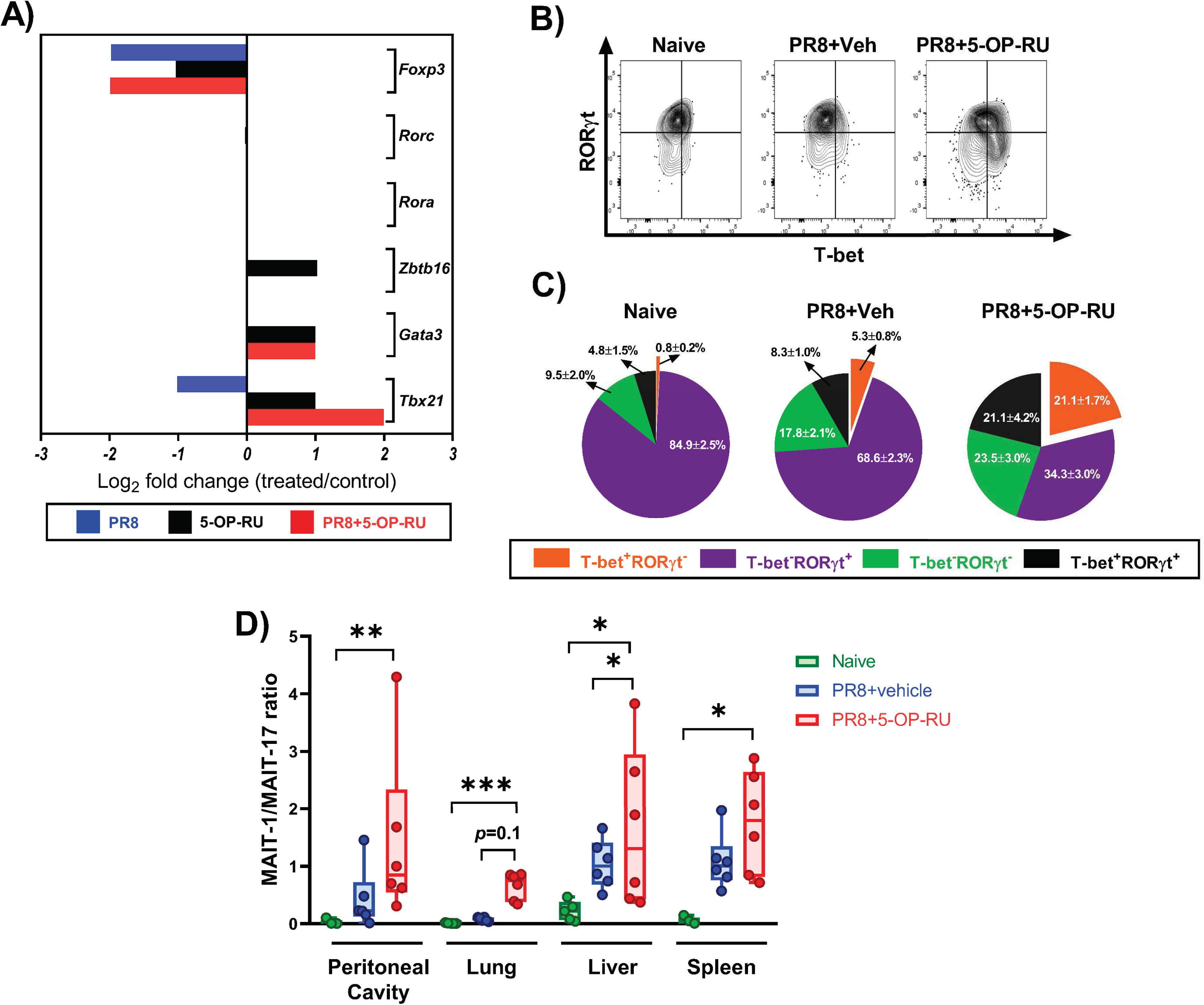
Immunization with PR8 plus 5-OP-RU promotes a MAIT1 program at transcriptional and protein levels. (**A**) B6-MAIT^CAST^ mice were injected *i.p.* with PBS (n=10), PR8 (n=10), 5-OP-RU (n=10), or PR8 plus 5-OP-RU (n=5). Three days later, pooled pulmonary MAIT cells were purified and subjected to RNA extraction, cDNA generation and quantitative PCR to assess the expression of indicated transcription factors. Gene expression fold changes in MAIT cells from each cohort relative to those isolated from control (PBS-injected) animals were calculated using the 2^-(ΔΔCt)^ method. (**B-C**) In additional experiments, B6-MAIT^CAST^ mice were left unimmunized (n=5) or inoculated *i.p.* with PR8 plus 5-OP-RU (or vehicle) (n=6/group). Three days later, pulmonary MAIT cells were interrogated by flow cytometry for their intracellular RORγt and T-bet contents. Representative plots illustrate the staining for the above transcription factors within TCRβ^+^ MR1 tetramer^+^ cells after using isotype controls to set quadrant gates (**B**), and pie charts depict the frequencies (± SEM) of T-bet^+^RORγt^-^, T-bet^-^RORγt^+^, T-bet^+^RORγt^+^ and T-bet^-^RORγt^-^ MAIT cell subsets (**C**). The percentages of T-bet^+^RORγt^-^ and T-bet^-^RORγt^+^ MAIT cells were used to calculate MAIT1:MAIT17 ratios in indicated sites for each cohort (**D**). Each circle in Box-and- Whisker plots corresponds to an individual animal (or pooled peritoneal sample). Statistical comparisons were performed using the Kruskal-Wallis test followed by the Dunn’s post-hoc test. *, ** and *** denote differences with *p* ≤ 0.05, *p* ≤ 0.01 and *p* ≤ 0.001, respectively.

To define MAIT1 and MAIT17 subsets unequivocally, we stained MAIT cells for intracellular transcription factors. As expected, T-bet^-^RORγt^+^ cells were the prominent MAIT subset in the lungs (Fig. 7B-C), liver, spleen and peritoneal cavity of B6-MAIT^CAST^ mice (Supplementary Fig. 13) whereas T-bet^+^RORγt^-^ and T-bet^+^RORγt^+^ subpopulations were minorities. Moreover, PR8 immunization without 5-OP-RU elevated T-bet^+^ cell frequencies only marginally. Remarkably, however, 5-OP-RU-adjuvanted vaccination gave rise to a large T-bet^+^ subset (Fig. 7B-C and Supplementary Fig. 13). As a result, a significantly higher MAIT1:MAIT17 ratio emerged (Fig. 7D), indicating a skewed response towards T helper-type 1 (T_H_1)-type cytokine production. We also detected a T-bet^-^RORγt^-^ MAIT cell subset at varying frequencies across different tissues and cohorts (Fig. 7B-C and Supplementary Fig. 13). However, we ruled out the possibility that GATA binding protein 3 (GATA-3)^+^ cells were a major component of this double-negative subpopulation (Supplementary Fig. 14).

Global cytokine gene expression analyses demonstrated increased *Ifng*, decreased *Il4*, and decreased or unaltered *Il17* levels in MAIT cells after immunization with PR8 and 5-OP-RU (Fig. 8A). To determine the extent to which MAIT cells were skewed, we stimulated bulk MNCs from naïve and vaccinated mice to the T_H_1-polarizing cytokines IL-12 and IL-18 before measuring MAIT cells’ IFN-γ and IL-17A contents. We found significantly higher IFN-γ^+^ cell percentages in mice that had received 5-OP-RU (Fig. 8B and Supplementary Fig. 15). By comparison, IL-17A^+^ MAIT cell frequencies were always low and comparable in the three experimental groups.

**Fig. 8:**
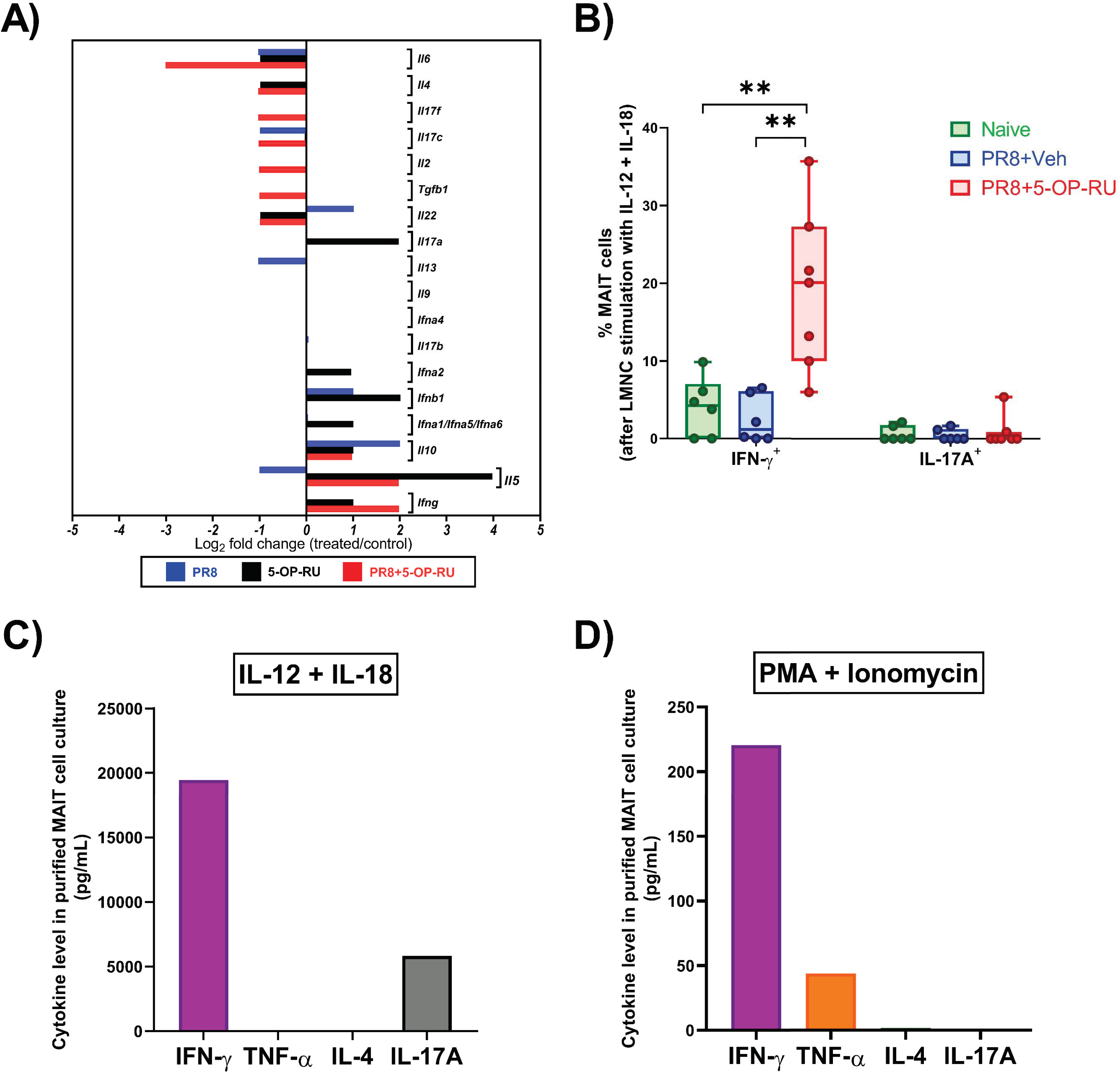
Co-administration of PR8 and 5-OP-RU confers upon MAIT cells a T_H_1-type cytokine profile. (**A**) B6-MAIT^CAST^ mice were injected *i.p.* with PBS (n=10), PR8 (n=10), 5-OP-RU (n=10), or PR8 plus 5-OP-RU (n=5). Three days later, the expression of indicated cytokines by pulmonary MAIT cells was quantified by real-time PCR for each cohort relative to PBS-injected animals. (**B**) Unfractionated non-parenchymal lung mononuclear cells (LMNCs) from naïve mice (n=6), animals immunized with PR8 plus 5-OP-RU (n=7), and animals immunized with PR8 plus vehicle (n=6) were stimulated *ex vivo* with recombinant mouse IL-12 and IL-18. After 18 hours, the frequencies of IFN-γ^+^ and IL-17A^+^ cells among TCRβ^+^ MR1 tetramer^+^ MAIT cells were determined by flow cytometry. One-way ANOVA was used, followed by the Tukey’s Multiple Comparison test, for statistical comparisons. ** denotes significant differences with *p* ≤ 0.01. (**C-D**) Pulmonary MAIT cells were purified from mice previously injected with PR8 and 5-OP-RU (n=10), pooled and stimulated with either a combination of IL-12 and IL-18 (**C**) or a combination of PMA and ionomycin (**D**). After 24 hours, cytokines levels were measured by ELISA in culture supernatants.

To assess MAIT cells’ cytokine production capacity in the absence of transactivating signals provided by other cell types, we incubated purified MAIT cells from mice that had received PR8 and 5-OP-RU with a panel of stimuli. First, exposure to IL-12 and IL-18 generated a large quantity of IFN-γ, which was ∼4 times more than IL-17A (Fig. 8C), consistent with cytofluorimetric data obtained from unfractionated MNC cultures (Fig. 8B and Supplementary Fig. 15). In contrast, using plate-coated anti-CD3 to cross-link *i*TCRs, alone or in combination with an anti-CD28 mAb resulted in IFN-γ and IL-17A secretion (Supplementary Fig. 16A-B). Therefore, a prior *in vivo* exposure to 5-OP-RU had not impeded MAIT cells’ ability to signal through their Ag receptors.

To examine MAIT cells’ polarization in an unbiased manner, without relying on *i*TCR and cytokine receptor signaling, we used a combination of PMA and ionomycin. PMA activates protein kinase C and ionomycin releases Ca^++^ from intracellular stores, thus inducing cytokine secretion^59^. In these experiments, purified MAIT cells from vaccinated animals produced large quantities of IFN-γ and some tumor necrosis factor (TNF)-a, but no detectable IL-4 or IL-17A (Fig. 8D).

Collectively, the above results indicate that vaccination with PR8 and 5-OP-RU expands, activates and skews MAIT cells towards a MAIT1 phenotype, which should favor antiviral immunity.

### Incorporating 5-OP-RU in an anti-IAV vaccination protocol affords heterosubtypic protection against IAV infection

To test the above hypothesis, we sought to explore the adjuvanticity of 5-OP-RU in a vaccination model in which *i.p.* priming with HK, which is not pathogenic in mice, precedes an *i.n.* challenge with the mouse-adapted PR8 strain (Fig. 9A).

**Fig. 9:**
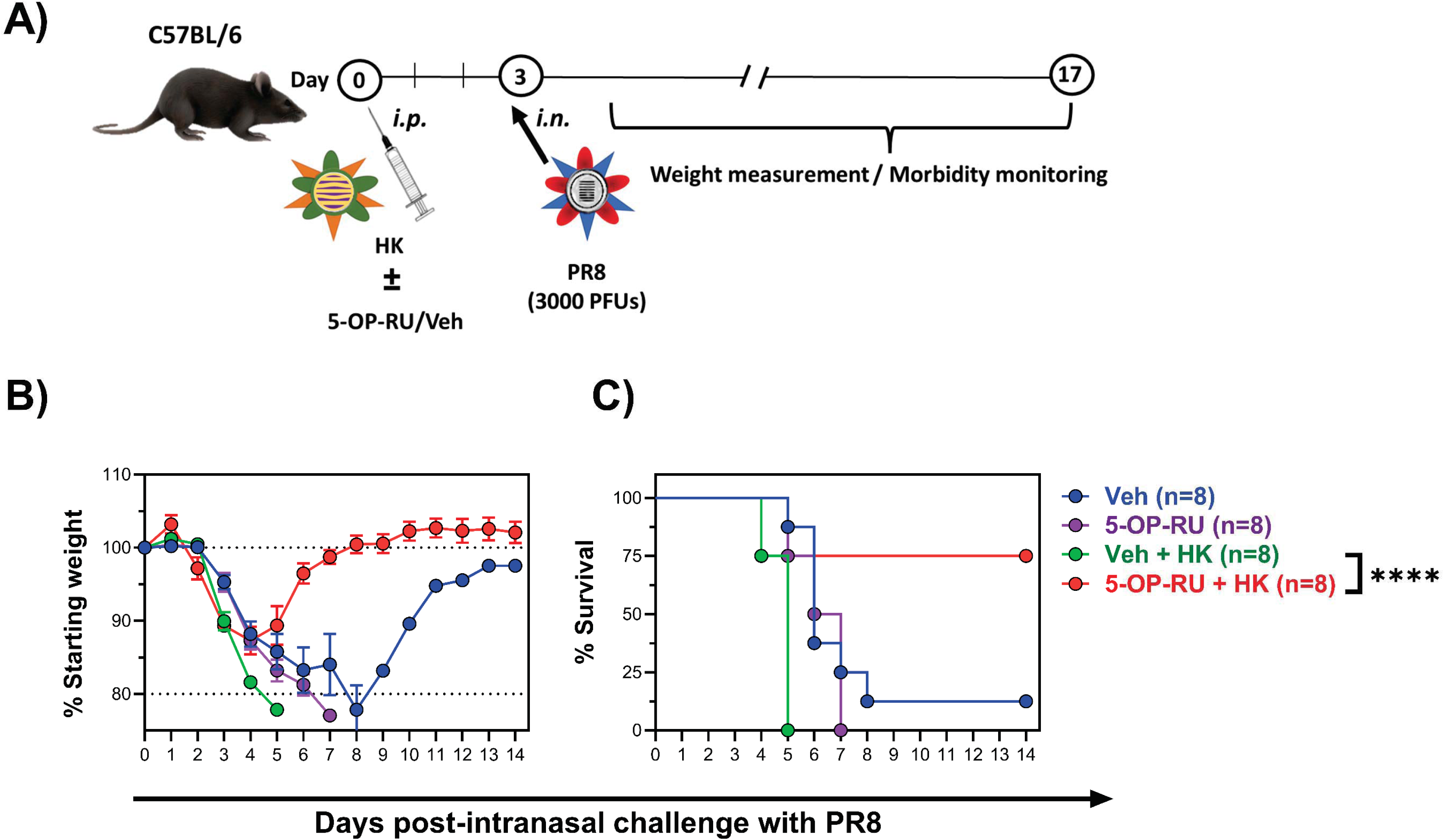
Vaccination with 5-OP-RU-adjuvanted IAV protects against a subsequent heterosubtypic challenge. (**A**) B6 mice were injected *i.p.* with the H3N2 IAV strain HK, with 5-OP-RU (or vehicle), or with a combination of HK and 5-OP-RU (or vehicle). Three days later, animals were challenged with an intranasal (*i.n.*) PR8 inoculum and monitored for weight loss (**B**) and mortality (**C**). Chi-squared tests were used in statistical analyses. **** denotes a significant difference with *p* ≤ 0.0001.

As expected, MAIT cells amassed in several tissues, including the lungs, 3 days after the *i.p.* inoculation of WT B6 mice with HK and 5-OP-RU (Supplementary Fig. 17). In these experiments, administering HK or 5-OP-RU alone failed to prevent PR8-induced weight loss and mortality (Fig. 9B-C). However, a combination of HK and 5-OP-RU protected the animals. Therefore, 5-OP-RU can be used as an adjuvant in anti-IAV immunization strategies.

### Adding 5-OP-RU to i.n. and i.m. vaccines for influenza and COVID-19 elevates MAIT cell numbers and augments CD8^+^ T_conv_ cell responses to immunodominant viral epitopes

To test the efficacy of 5-OP-RU in inducing mucosal immunity, we inoculated BALB/c mice *i.n.* with the live attenuated influenza vaccine FluMist (Fig. 10A). We used BALB/c mice in this model because the IAVs used in FluMist, similarly to the PR8 strain, harbor an immunodominant peptide called NP_147-155_, which is restricted by H-2K^d^ and therefore detectable in this mouse strain ^60^. As such, these experiments enabled us to enumerate not only MAIT cells but also NP_147_-specific CD8^+^ T_conv_ cells.

**Fig. 10:**
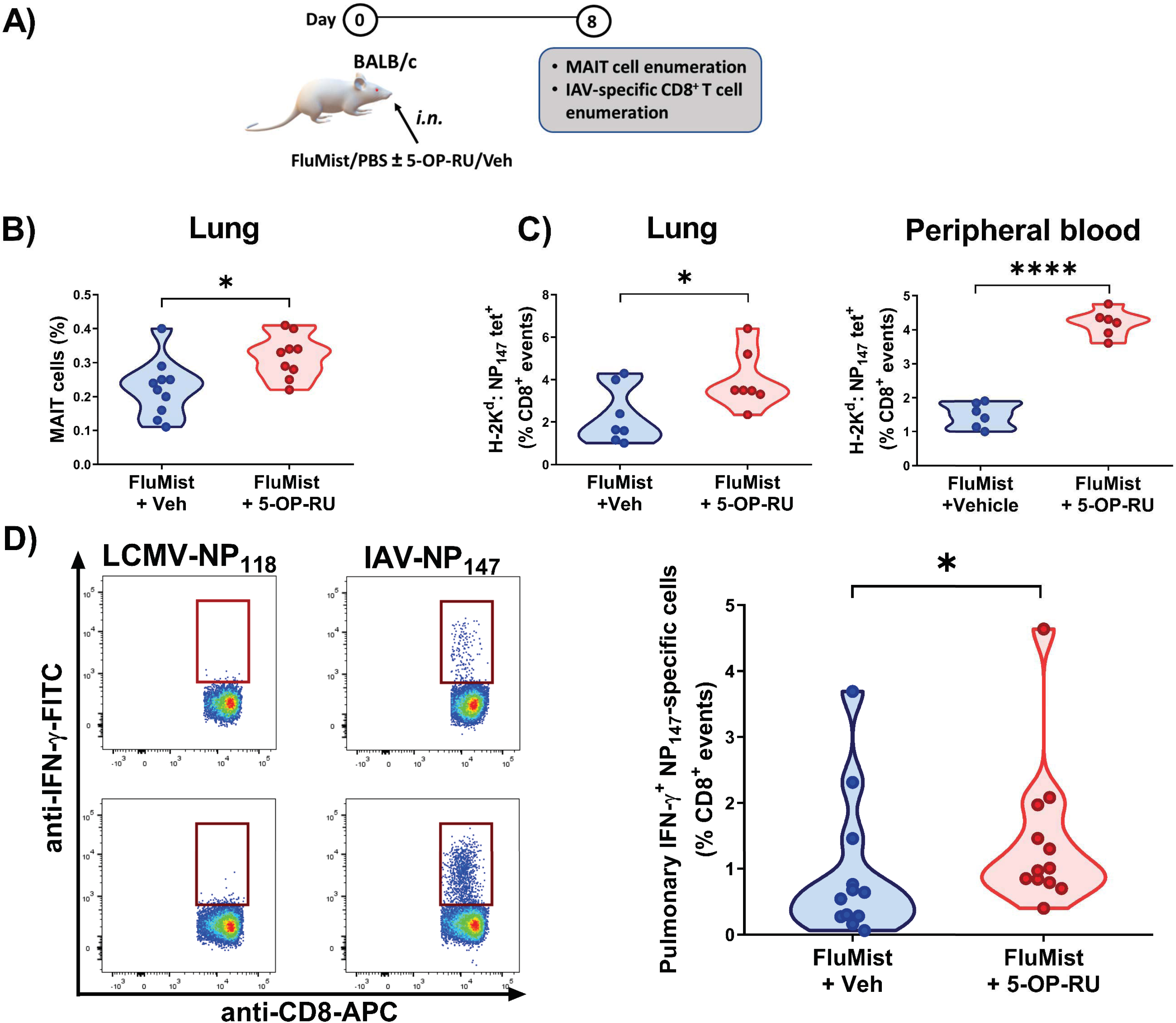
Intranasal co-administration of FluMist and 5-OP-RU to BALB/c mice increases the frequencies of pulmonary MAIT cells and immunodominant IAV-specific CD8^+^ T cells. (**A**) BALB/c mice were inoculated intranasally (*i.n.*) with the live attenuated influenza vaccine FluMist plus 5-OP-RU (or vehicle). Eight days later, pulmonary MAIT cells were enumerated by MR1 tetramer staining (**B**), and NP_147_-specific CD8^+^ T cell frequencies were determined by MHC I tetramer staining in the lungs and in the peripheral blood (**C**). (**D**) In parallel or in addition, immunodominant NP_147_-specific CD8^+^ T cells were identified by intracellular IFN-γ staining after non-parenchymal lung mononuclear cells (LMNCs) from vaccinated animals were stimulated *ex vivo* with TYQRTRALV. LMNCs were also incubated with an irrelevant peptide (RPQASGVYM) corresponding to the most immunodominant epitope of the lymphocytic choriomeningitis virus nucleoprotein (LCMV-NP_118_). Representative plots and summary data are depicted (**D**). Each circle represents an individual mouse (**B-D**). Statistical analyses were conducted using unpaired *t*-tests (**B-C**) and Mann-Whitney *U* test (**D**). * and **** denote differences with *p* ≤ 0.05 and *p* ≤ 0.0001, respectively.

As with our *i.p.* vaccination models, combining FluMist with 5-OP-RU raised MAIT cell frequencies in the lungs (Fig. 10B). Furthermore, MHC class I tetramer staining revealed a numerical increase in pulmonary and peripheral blood NP_147_-specific T_conv_ cells (Fig. 10C). These were functional, Ag-specific T cells as judged by their ability to produce IFN-γ upon brief *ex vivo* stimulation with a synthetic peptide corresponding to NP_147-155_, but not an irrelevant peptide derived from LCMV (NP_112-126_), in ICS assays (Fig. 10D).

Next, we evaluated the adjuvanticity of 5-OP-RU in *i.m.* immunization against another respiratory pathogen. To this end, BALB/c mice were injected in the hind limb with an rVSV-vectored vaccine candidate encoding the SARS-CoV-2 Spike (Fig. 11A), which we recently generated ^37^. In this setting, we again found more MAIT cells in mice receiving 5-OP-RU on days 3 and 7 post-vaccination, especially in the lungs (Fig. 11B). In addition, cognate CD8^+^ T_conv_ cells recognizing an immunodominant determinant of the SARS-CoV-2 Spike (S_535-543_) ^47^ were more abundant in animals that had received 5-OP-RU (Fig. 11C).

**Fig. 11:**
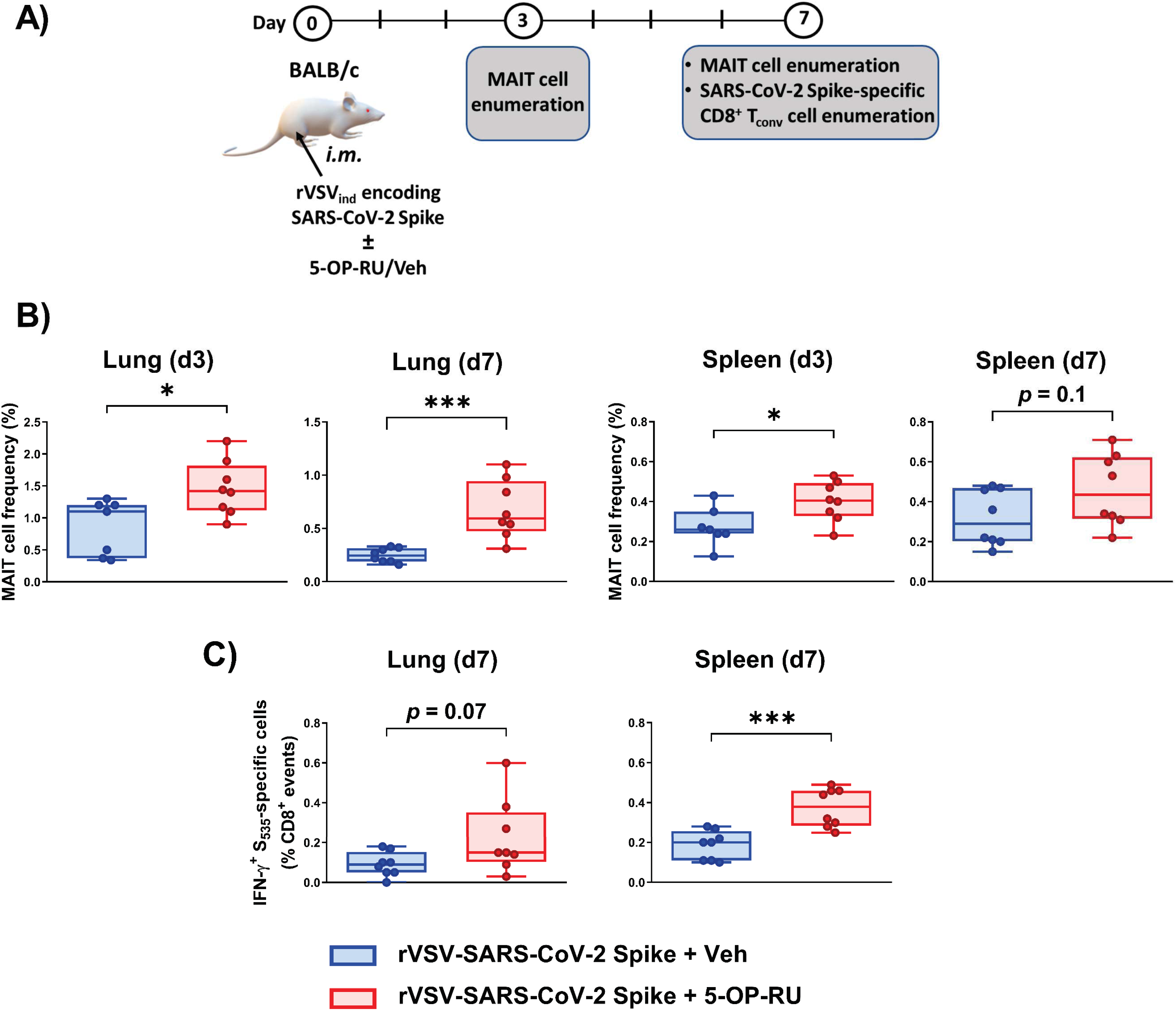
Intramuscular co-administration of an rVSV-based COVID-19 vaccine and 5-OP-RU raises the frequencies of pulmonary and splenic MAIT cells and cognate CD8^+^ T cell. BALB/c mice were injected intramuscularly (*i.m.*) with rVSV_Ind_ expressing the SARS-CoV-2 Spike (S) gene plus 5-OP-RU (or vehicle) (**A**). Three or 7 days later, pulmonary and splenic MAIT cells were enumerated by MR1 tetramer staining (**B**). On day 7 post-immunization, the percentages of S_535_-specific CD8^+^ T cells were also determined by intracellular cytokine staining for IFN-γ (**C**). Each circle in Box-and-Whisker plots represents an individual mouse. * and *** denote statistically significant differences with *p* ≤ 0.05 and *p* ≤ 0.001, respectively, by unpaired *t*-tests.

In a limited number of experiments, we included 5-OP-RU or vehicle during *i.m.* priming with rVSV_Ind_-Spike and *i.m.* boosting with rVSV_NJ_-Spike. Using this prime-boost immunization model, a significant increase in pulmonary and splenic MAIT cell numbers was noticeable in the cohort that had received 5-OP-RU during both phases (Supplementary Fig. 18).

### Exposing human PBMCs to propagating PR8 virions and 5-OP-RU results in MAIT cell activation and proliferation

To extend our studies to human MAIT cells, we cultured PBMCs with uninfected or PR8-infected C1R-MR1 cells in the absence or presence of 5-OP-RU before we determined the expression levels of several activation markers (Fig. 12A). We found significant upregulation of CD38 (Fig. 12B) and human leukocyte antigen (HLA)-DR (Fig. 12C) in cultures containing PR8-infected, 5-OP-RU-pulsed C1R-MR1 cells. This was clearly evident as early as 16 hours after the cultures were initiated and prevailed at least for 4 days (Fig. 12B-C).

**Fig. 12:**
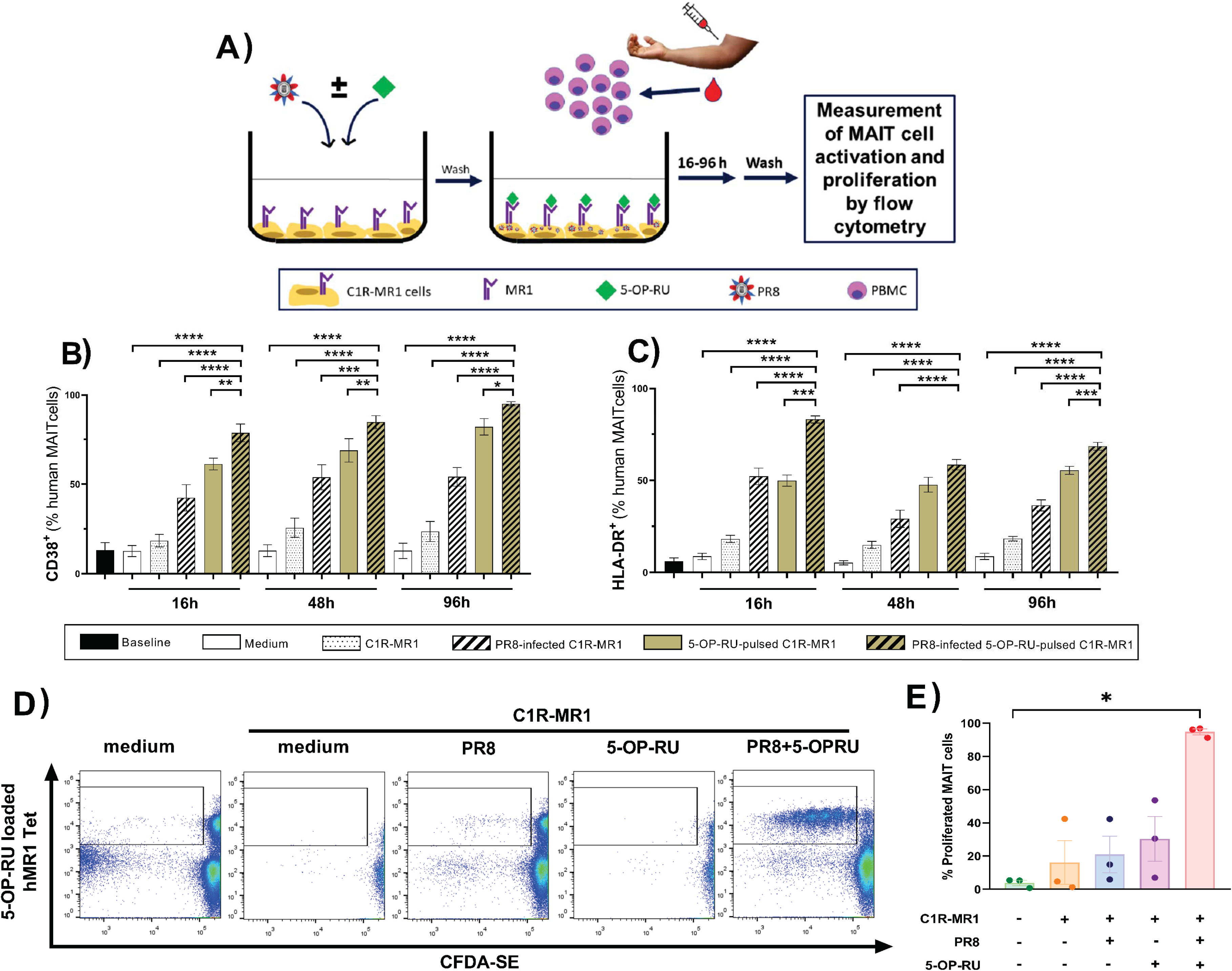
*In vitro* stimulation of human PBMCs with live PR8 and 5-OP-RU activates and expands MAIT cells. Human peripheral blood mononuclear cells (PBMCs) from healthy donors were incubated in complete medium (n=6) or stimulated with C1R-MR1 cells that had been infected with PR8 and/or pulsed with 5-OP-RU (n=10 per condition) (**A**). At indicated timepoints, the frequencies of CD38^+^ (**B**) and HLA-DR^+^ (**C**) cells among CD3^+^ hMR1 tetramer^+^ MAIT cells were determined by flow cytometry. (**D-E**) In additional experiments, PBMCs were labelled with CFDA-SE before they were stimulated as described above. Seven days later, CFDA-SE dye dilution by hMR1 tetramer-positive (MAIT) and -negative (non-MAIT T) cells was assessed after gating on CD3^+^ events. Representative plots (**D**) and summary data (n=3 donors) (**E**) are depicted. Data are shown as mean ± SEM (**B, C and E**). Statistical comparisons were made using Repeated Measures one-way ANOVA with the Dunnett’s Multiple Comparisons test (**B-C**) and Friedman tests followed by Dunn’s post-hoc analyses (**E**). *, **, *** and **** denote differences with *p* ≤ 0.05, *p* ≤ 0.01, *p* ≤ 0.001 and *p* ≤ 0.0001, respectively.

In separate cultures, we used CFDA-SE-labeled PBMCs in order to test the proliferative capacity of human MAIT cells. Seven days later, CFDA-SE dye dilution, indicating up to six cell division cycles, was detectable when PR8-infected, 5-OP-RU-pulsed C1R-MR1 cells were present in cultures (Fig. 12D-E). A similar trend was found when cultured cells were analyzed on day 4 (Supplementary Fig. 19). This response was exclusive to MAIT cells since non-MAIT T cells did not divide (Fig, 12D). The *in vitro* nature of these experiments also reinforces the idea that MAIT cell expansion can be due primarily, if not purely, to their inherent proliferative activity after they are exposed to IAVs and 5-OP-RU.

## Discussion

The presence and activation status of MAIT cells influences antiviral immune responses and the severity of viral diseases, with both protective and pathogenic roles demonstrated or proposed for these powerful lymphocytes ^21,22^. For instance, while MAIT cells are thought to contribute to anti-IAV immunity ^12,14^, their CD69 or HLA-DR levels have been reported to predict the severity or mortality of COVID-19 in most, but not all, studies conducted on the subject to date^61–65^.

MAIT cells are MR1-testricted T cells with potent antibacterial activities. Therefore, it is only fitting that administration of live bacterial vaccine strains or the MR1 ligand 5-OP-RU promotes protection against *Legionella longbeachae* ^66^, *Francisella tularensis* ^67^ and *Vibrio cholerae* ^68^. However, since viruses do not generate MR1 ligands, our initial observation that 5-OP-RU synergized with IAV vaccines to potentiate MAIT cell accumulation in multiple tissues was intriguing. Ki-67 staining, *in vivo* S1PR1 antagonism, and adoptive transfer experiments collectively demonstrated that MAIT cell proliferation, rather than their emigration from other sites, was responsible for the above phenomenon, a finding that could also be recapitulated in our human MAIT cell cultures. Of note, in a similar PBMC culture system, Lamichhane *et al.*^69^ found PR8 to augment *i*TCR-mediated MAIT cell activation and degranulation as judged by their increased CD69, PFN, GZM B and CD107a levels. However, no MAIT cell proliferation was reported. This may be due to the fact that the PR8 virions were subjected to ultraviolet radiation by these investigators. In contrast, in our system, replication-competent PR8 was used and induced MAIT cell proliferation.

Several lines of evidence point to a requirement for dsRNA-sensing pattern recognition receptors (PRRs), such as TLR3, in our system. First, HI-PR8, which is immunogenic but unable to propagate, failed to synergize with 5-OP-RU to induce peritoneal MAIT cell proliferation. Similarly, *i.m.* co-inoculation of HI-PR8 and 5-OP-RU did not increase MAIT cell numbers in draining lymph nodes (data not shown). Therefore, live viral replication, which generates dsRNA, appeared to be necessary for the observed MAIT cell response. We are cognizant of the previous reports that negative-sense RNA viruses like PR8 are not effective producers of dsRNA ^70–72^. However, the fact that MAIT cell expansion was evident when 5-OP-RU was combined with a variety of viruses and vaccines, including multiple influenza strains (*i.e.*, PR8, NT60, HK, X31 and FluMist virions), a vaccine vector based on a different negative-strand RNA virus (*i.e.*, VSV), and a DNA virus (*i.e.*, VacV), supports the notion that a common pathogen-associated molecular pattern (PAMP) was at play. We posited that viral dsRNA and TLR3 were the PAMP and the PRR involved, respectively. Indeed, poly (I:C), a synthetic mimic of dsRNA that is sensed by TLR3, induced MAIT cell proliferation when co-administered *i.p.* with 5-OP-RU. This finding is consistent with a previous report that *i.n.* inoculation of poly (I:C) before 5-OP-RU expands pulmonary MAIT cells in WT B6 mice^73^. It is also noteworthy that naked low-molecular-weight poly (I:C), but not LyoVec-complexed high-molecular-weight poly (I:C), was efficient in this capacity in our model (data not shown). Therefore, melanoma differentiation-associated gene 5 (MDA5), a cytosolic PRR that recognizes long dsRNA^74^, seems not to play a prominent role in MAIT cell expansion. We are currently studying the molecular intermediates of intracellular signaling pathways that contribute to or control MAIT cell responses, including TLR3- and possibly retinoic acid-inducible gene-I (RIG-I)-coupled pathways.

A key consequence of TLR3 engagement is the production of type I IFNs, to which MAIT cells react ^11,13,27^. The elevated blood levels of IFN-β in mice receiving PR8 plus 5-OP-RU, and the observed synergy between rmIFNβ and 5-OP-RU implicated IFNAR signaling in our experimental systems. This was confirmed by using an IFNAR-1-blocking mAb, which prevented MAIT cell proliferation. Furthermore, adoptive transfer experiments enabled head-to-head comparison between IFNAR-sufficient and -deficient cells in the same host and led us to conclude that IFNAR signaling within MAIT cells was required for their proliferation. This is in agreement with a previous study in which costimulation with IFN-a or IFN-β enhanced IFN-γ and GZM B production by human MAIT cells in response to anti-CD3/CD28-coated beads *in vitro*^69^.

How exactly *i*TCR triggering synergizes with type I IFNs to expand tissue MAIT cell populations is speculative at this point. According Stevens *et al.* ^75^, some of the early features of TCR- and IFNAR-coupled cascades, including mitogen-activated protein kinase (MAPK) phosphorylation, are identical in Jurkat cells and primary peripheral blood T_conv_ cells. However, type I IFN-induced MAPK activation is relatively short-lived and does not induce the same transcriptional responses that follow TCR stimulation. Therefore, the possibility that 5-OP-RU and type I IFNs generate cell division-promoting signals that merge propitiously and culminate in MAIT cell proliferation seems far-fetched. During LCMV infection, type I IFNs act directly on antiviral CD8^+^ T_conv_ cells to keep them alive in the Ag-specific clonal expansion phase ^76,77^. By extrapolating from these findings, one could envisage a scenario in which IFNAR signaling in MAIT cells allows them to survive in order to receive and react to a cognate proliferative signal that is transmitted after 5-OP-RU recognition. This theory may also explain the absence of adoptively transferred IFNAR^-/-^ cells in IFNAR^+/+^ recipients co-injected with PR8 plus 5-OP-RU.

While many cell types release type I IFNs during viral infections or vaccination, plasmacytoid DCs (pDCs) are particularly efficient in producing copious amounts of these cytokines ^78,79^. Our current understanding of how pDCs may communicate with MAIT cells is rudimentary. The immunogenicity of adenovirus-based vaccine vectors was recently reported to depend on MAIT cells whose activation in turn required IFN-a production by pDCs ^27^. Unlike myeloid DCs, pDCs rely on TLR7, not TLR3, to sense the presence of viruses ^80,81^, and the TLR7 agonist imiquimod did not synergize with 5-OP-RU to expand MAIT cells. Therefore, pDCs are unlikely to play a critical role in our models.

Provine *et al.* ^27^ investigated the activation status of MAIT cells and their potential targetability when using replication-incompetent vectors. In contrast, we have explored and documented the adjuvanticity of an MR1 ligand, namely 5-OP-RU, using multiple immunization modes and routes. Furthermore, we demonstrate that MAIT cell proliferation in vaccination settings requires replication-competent vaccine strains or vectors. This may be why Provine *et al.* did not report MAIT cell expansion in their study.

Importantly, 5-OP-RU-adjuvanted IAV vaccines not only expanded MAIT cells but also protected against a lethal challenge with PR8. This may be attributable to a skewed MAIT1 response characterized by T-bet expression and pro-inflammatory cytokine production, which should enable MAIT cells to transactivate a number of antiviral secondary effectors, including NK cells and CD8^+^ T_conv_ cells. In fact, in our FluMist and rVSV-based immunization models for influenza and COVID-19, respectively, we had the opportunity to enumerate immunodominant CD8^+^ T_conv_ cells, which were markedly increased in animals receiving 5-OP-RU-adjuvanted vaccines.

In our prime-boost anti-IAV immunization protocol, a secondary exposure to 5-OP-RU was sufficient to induce tissue MAIT cell accumulation (Fig. 2 and Supplementary Fig. 3). In our rVSV-based protocol for COVID-19, the optimal response was achieved when 5-OP-RU was given during both the priming and the boosting phase (Supplementary Fig. 17). These experiments clearly demonstrate that 5-OP-RU inoculation does not render MAIT cells unresponsive to subsequent 5-OP-RU administrations. Consistent with these observations, we found no transcriptional signs of overt cellular exhaustion (Supplementary Fig. 12) or proneness to apoptotic death (Supplementary Fig. 20) in MAIT cells from animals that had received 5-OP-RU as a vaccine adjuvant. These studies will be extended in the future to include additional genes. Regardless, repeated 5-OP-RU administration, when/if necessary, should be feasible in preventative or even in therapeutic vaccination strategies. This is in sharp contrast with *i*NKT cells, sometimes dubbed as the mouse cousins of human MAIT cells, which undergo long-term anergy via a programmed death-1 (PD-1)/programmed death ligand-1 (PD-L1)-dependent mechanism in animals receiving a single injection of the prototypic CD1d ligand α-galactosylceramide (α-GalCer) ^82,83^. As a result, anergic *i*NKT cells lose their responsiveness to future encounters with the same glycolipid Ag and fail to produce several cytokines, including IFN-γ. This may explain, at least partially, why repeated injections of free-floating α-GalCer have not worked optimally in clinical trials for solid tumors and viral diseases ^84^. Future work will need to address the most tolerable and productive 5-OP-RU dosages, injection frequencies and time intervals.

As a non-classical adjuvant, 5-OP-RU was able to expand MAIT cells in young and old mice, female and male, of different MHC backgrounds. The evolutionarily conserved nature of MAIT cells and their selection mode ^85^, the fact that mouse MAIT cells resemble their human counterparts more closely than previously appreciated ^86^, and given many reports, including ours, that 5-OP-RU activates both mouse and human MAIT cells, our findings should be translatable to clinical settings. Since MR1 is monomorphic, its ligands should work in genetically diverse human populations. This is unlike T_conv_ cell-based interventions that need to be tailored to individual HLA backgrounds.

Opportunistic bacterial and fungal infections are among the most catastrophic complications of viral infections. For instance, community-acquired and nosocomial bacterial pneumonia claim many lives after IAV and SARS-CoV-2 infections ^87,88^. MAIT cells should help clear such infections by at least four mechanisms. First, their stimulation bolsters other innate and adaptive effectors with antibacterial activities ^25–29^. Second, MAIT cells directly recognize and kill infected cells displaying bacterial (and fungal) MR1 ligands ^2,31^. Third, cytolytic effector molecules within the MAIT cells’ arsenal may damage free-living extracellular bacteria, and synergize with the bactericidal activities of certain antibiotics such as carbapenems ^32^. Of note, our global gene expression analyses revealed upregulated *Gzmb*, *Gzmk* and *Fasl* transcript levels in MAIT cells from mice receiving PR8 plus 5-OP-RU (Supplementary Fig. 21). These cytolytic effector molecules should help augment MAIT cells’ antibacterial functions. Finally, MAIT cell roles in tissue repair and remodeling ^38,89–92^ should help resolve structural impairments that may elevate the risks of continued or *de novo* infections. For all of the above reasons, we propose that reinstating or reinforcing tissue MAIT cell compartments will protect against secondary infections and superinfections. Importantly, the above functions are typically, if not exclusively, dependent on the MR1/*i*TCR-dependent pathway, which is targeted by 5-OP-RU. Efforts to expand and store MAIT cells as donor MHC-unrestricted, off-the-shelf immunotherapeutics are underway ^43^. Whether our *in vivo* findings that poly (I:C) and type I IFN enhance the ability of 5-OP-RU to expand MAIT cells can be extended to *in vitro* platforms for future human cell therapies is a subject of ongoing work.

In summary, although MR1 ligands are not generated by viruses and virus-based vaccines, we have herein demonstrated that MR1 can be targeted by 5-OP-RU towards more efficacious antiviral immunization strategies. 5-OP-RU synergizes with multiple vaccines to expand pro-inflammatory MAIT1 cells in a TLR3/IFNAR-dependent manner, to enhance antiviral CD8^+^ T_conv_ cell responses, and to enable heterosubtypic anti-IAV immunity. Therefore, we propose 5-OP-RU as a potent and versatile vaccine adjuvant against respiratory viral pathogens.

## Abbreviations

7-AAD: 7-aminoactinomycin D
Ag: antigen
α-GalCer: α-galactosylceramide
AUP(s): animal use protocol(s)
B6: C57BL/6
BFA: brefeldin A
CFDA-SE: carboxyfluorescein diacetate succinimidyl ester
COVID-19: coronavirus disease 2019
Ct: cycle threshold
DC(s): dendritic cell(s)
DMSO: dimethyl sulfoxide
dsRNA: double-stranded ribonucleic acid
FBS: fetal bovine serum
6-FP: 6-formylpterin
GATA-3: GATA binding protein 3
gMFI: geometric mean fluorescence intensity
GZM: granzyme
HA: hemagglutinin
HI-PR8: heat-inactivated A/Puerto Rico/8/1934 (PR8) [IAV strain]
HK: A/Hong Kong/1/1968 [influenza A virus (IAV) strain]
HLA: human leukocyte antigen
HMNCs: [non-parenchymal] hepatic mononuclear cell(s)
ICS: intracellular cytokine staining
IFN: interferon
*Ifnar1*: IFN-α/β receptor subunit 1
IL: interleukin
*i*NKT: invariant natural killer T [cell]
*i.m.*: intramuscular(ly)
*i.n.*: intranasal(ly)
*i.p.*: intraperitoneal(ly)
*i*TCR: invariant T cell receptor
*i.v.*: intravenous(ly)
LCMV: lymphocytic choriomeningitis virus
LMNC(s): [non-parenchymal] lung mononuclear cell(s)
mAb: monoclonal antibody
MAIT: mucosa-associated invariant T [cell]
MAPK: mitogen-activated protein kinase
MDA5: melanoma differentiation-associated gene 5
MHC: major histocompatibility complex
MR1: MHC-related protein 1
MNC(s): mononuclear cell(s)
NA: neuraminidase
NK: natural killer [cell]
NKG2D: natural killer group 2, member D
NP: nucleoprotein
NT60: A/Northern Territory/60/1968 [IAV strain]
5-OP-RU: 5-(2- oxopropylideneamino)-6-D-ribitylaminouracil
PAMP: pathogen-associated molecular pattern
PBMC(s): peripheral blood mononuclear cell(s)
PBS: phosphate-buffered saline
PCR: polymerase chain reaction
PD-1: programmed death-1
pDC(s): plasmacytoid dendritic cell(s)
PD-L1: programmed death ligand-1
PE: phycoerythrin
PFN: perforin
PFU(s): plaque-forming unit(s)
PMA: phorbol 12-myristate 13-acetate
poly (I:C): polyinosinic-polycytidylic acid
PRR(s): pattern recognition receptor(s)
RIG-I: retinoic acid-inducible gene-I
rmIFNβ: recombinant mouse IFN-β
RORγT: retinoic acid receptor-related orphan receptor γt
rVSV(s): recombinant vesicular stomatitis virus(s)
SARS-CoV-2: severe acute respiratory syndrome coronavirus 2
S1PR1: sphingosine-1-phosphate receptor 1
T-bet: T-box expressed in T cells
TCID_50_: 50% tissue culture infectious dose
T_conv_: conventional T [cell]
T_H_: T helper [cell]
TLR: Toll-like receptor
TNF: tumor necrosis factor
VacV: vaccinia virus

## Acknowledgments

This work was funded by the Canadian Institutes of Health Research (CIHR) through a Project Grant (PJT 174984) to SMMH. CMW was and NIW is a recipient of a Frederick Banting and Charles Best Canada Graduate Scholarship – Master’s (CGS-M) from CIHR. We thank Dr. Olivier Lantz (Institut Curie) for critically reviewing this manuscript and all members of the Haeryfar Laboratory for helpful discussions. Dr. Patrick Rudak, Dr. Kate Parham, Connor Richer, Art Marzok and Alisha Kang provided expert technical assistance.

## Supplementary Figure Legends

**Supplementary Fig. 1:**
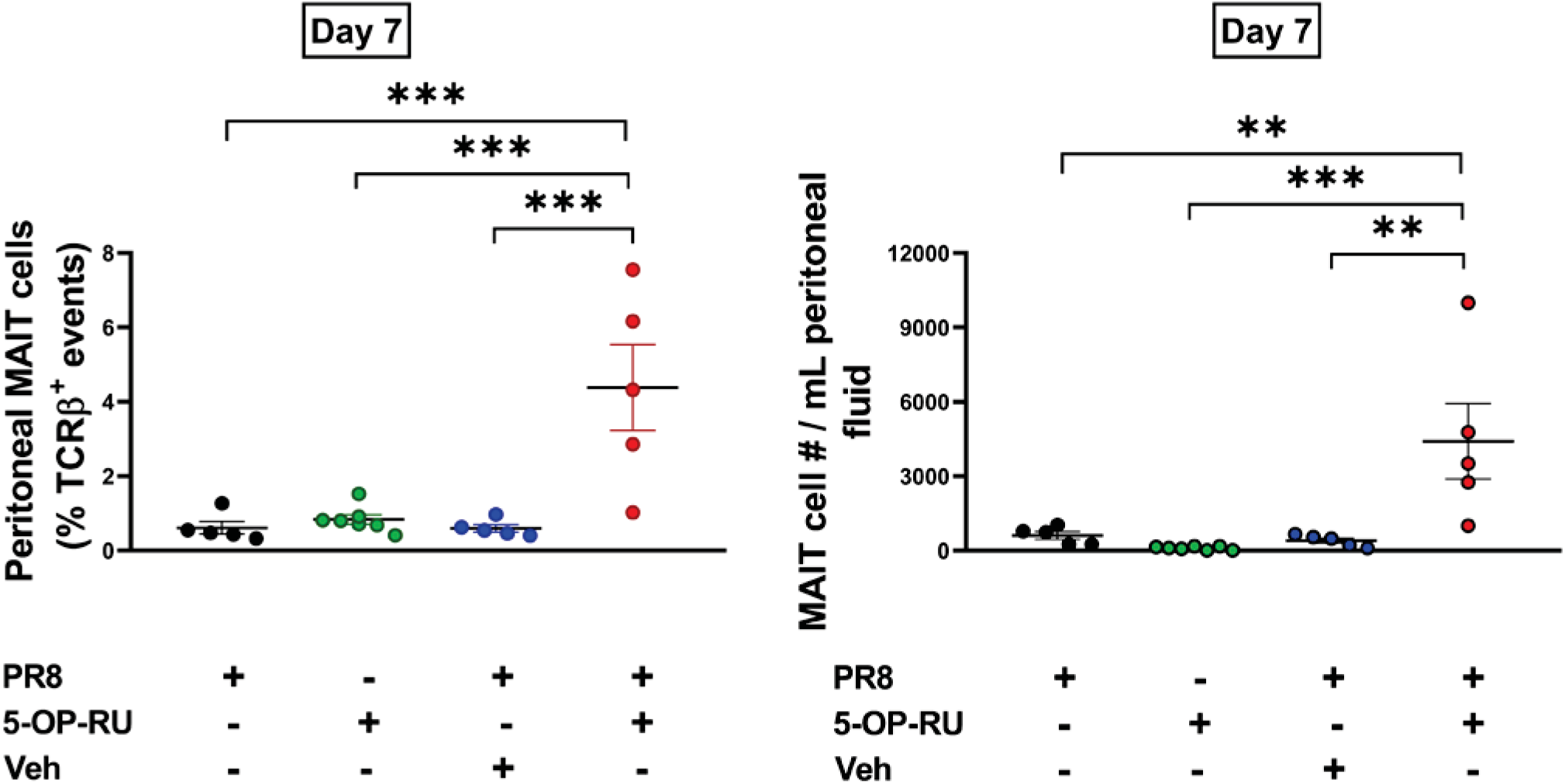
Intraperitoneal MAIT cell accumulation after immunization with PR8 and 5-OP-RU lasts longer than three days. B6-MAIT^CAST^ mice were inoculated *i.p.* with PR8, 5-OP-RU, or a combination of PR8 and 5-OP-RU (or vehicle) followed, 7 days later, by cytofluorimetric enumeration of MAIT cells in the peritoneal cavity. Peritoneal MAIT cell frequencies and absolute numbers are depicted. Each circle corresponds to an individual animal, and data are presented as mean ± SEM. Statistical comparisons were made using the one-way ANOVA followed by the Dunnett’s post-hoc Multiple Comparisons test. ** and *** signify differences with *p* ≤ 0.01 and *p* ≤ 0.001, respectively.

**Supplementary Fig. 2:**
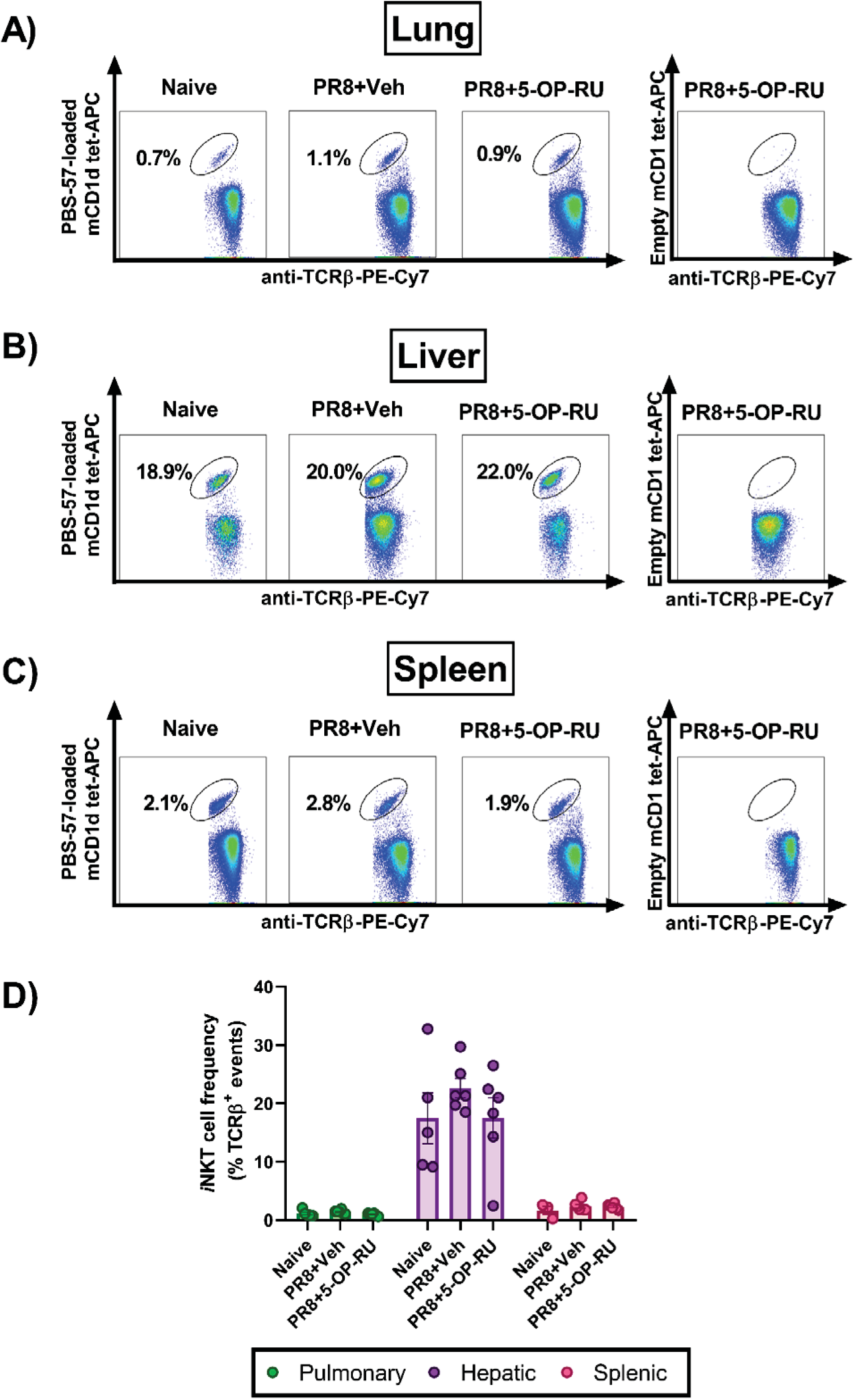
Immunization with 5-OP-RU-adjuvanted PR8 does not affect tissue *i*NKT cell frequencies. Naïve B6-MAIT^CAST^ mice (n=5) and mice that had received PR8 plus 5-OP-RU (or vehicle) three days earlier (n=6) were sacrificed for their lungs (**A and D**), liver (**B** and **D**) and spleen (**C-D**) in which TCRβ^+^ PBS-57-loaded mCD1d tetramer^+^ *i*NKT cells were enumerated. Empty mCD1d tetramers were used in parallel to draw cytofluorimetric gates (**A-C**). Representative plots (**A-C**) and summary data with mean ± SEM values (**D**) are shown. Group comparisons were made using the Kruskal-Wallis test followed by the Dunn’s post-hoc analysis.

**Supplementary Fig. 3:**
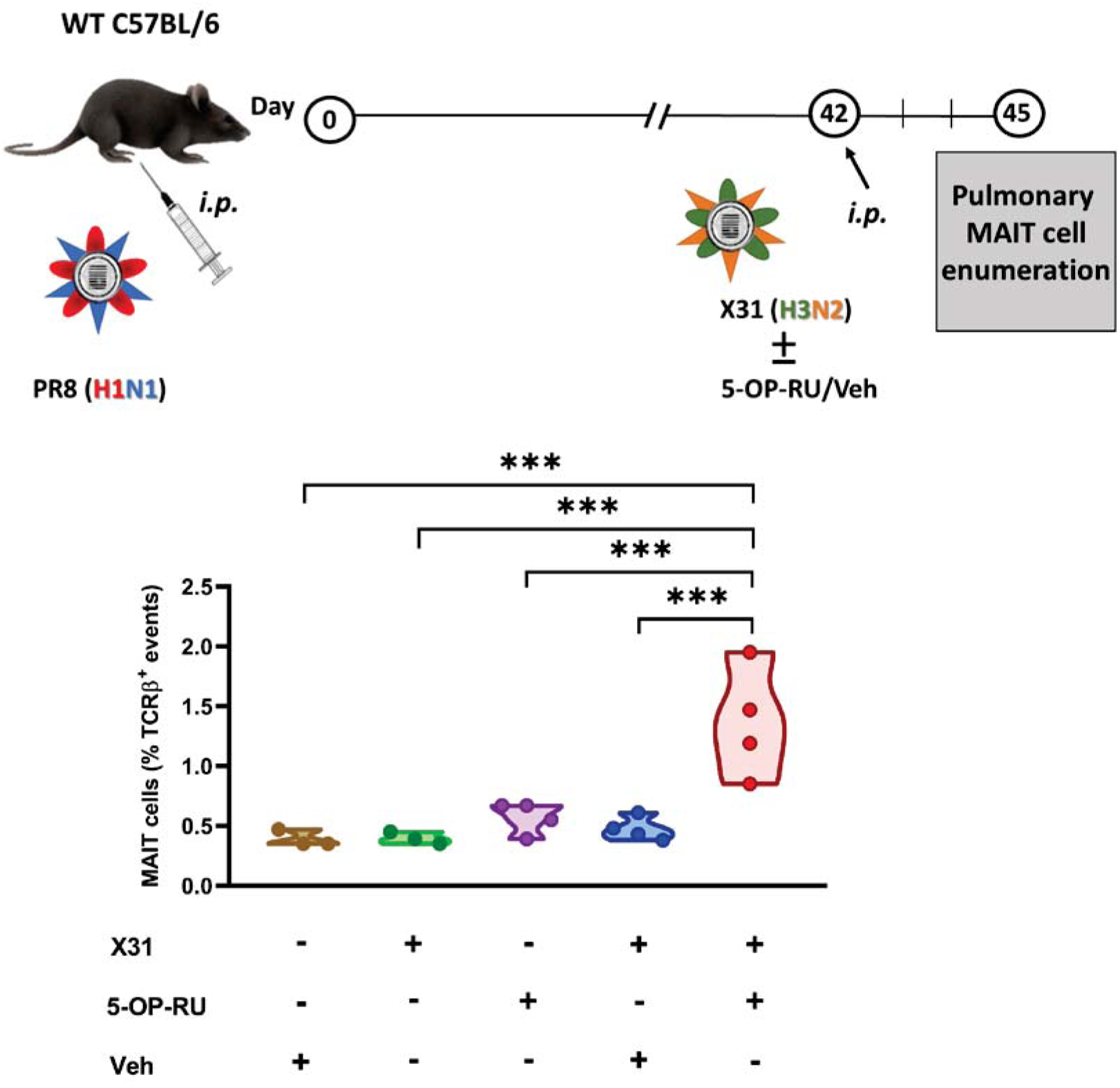
5-OP-RU elevates pulmonary MAIT cell numbers when administered during boosting immunization against IAV. B6 mice (n=3-4) were primed *i.p.* with PR8 and boosted, six weeks later, with an *i.p.* inoculum of the reassortant X31 strain plus 5-OP-RU (or vehicle). Additional cohorts received 5-OP-RU or vehicle alone as indicated. Three days after the secondary injection/immunization, MAIT cells were enumerated by flow cytometry in the lungs. Each circle represents an individual animal. *** denotes a statistically significant difference with *p* ≤ 0.001 by one-way ANOVA followed by the Holm-Šidék Multiple Comparisons test.

**Supplementary Fig. 4:**
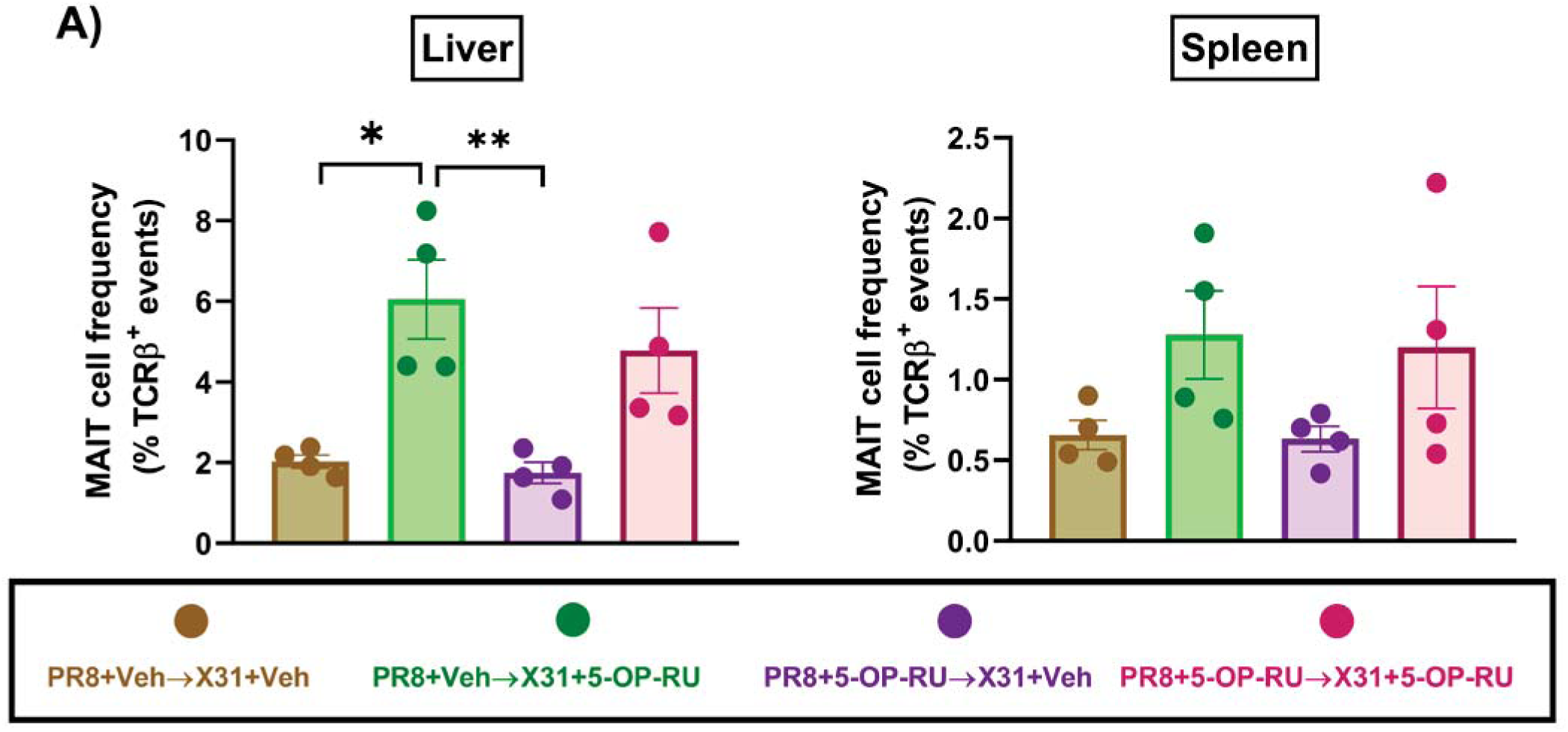
Incorporating 5-OP-RU into boosting anti-IAV immunization elevates the frequency of hepatic MAIT cells. B6-MAIT^CAST^ mice were primed *i.p.* with PR8 (H1N1) and boosted *i.p.*, four weeks later, with X31 (H3N2). 5-OP-RU (or vehicle) was co-administered in both phases. Three days after secondary immunization, hepatic (**A**) and splenic (**B**) MAIT cells were enumerated by flow cytometry. Each circle represents an individual mouse, and data are shown as mean ± SEM. * and ** denotes a significant difference with *p* ≤ 0.05 and *p* ≤ 0.01, respectively, using one-way ANOVA followed by the Tukey’s Multiple Comparisons test.

**Supplementary Fig. 5:**
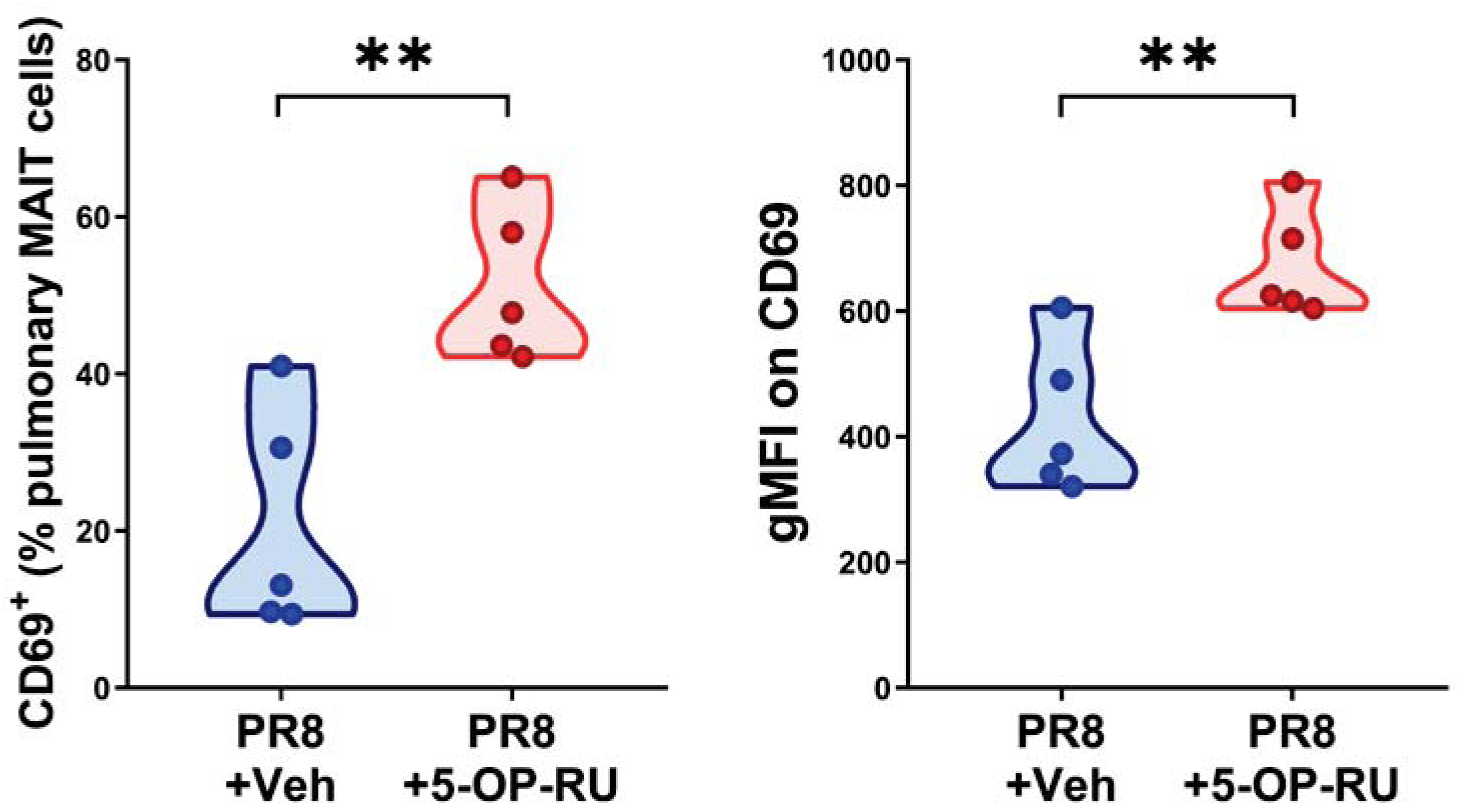
Intraperitoneal co-inoculation of PR8 and 5-OP-RU upregulates CD69 expression by pulmonary MAIT cells. B6-MAIT^CAST^ mice (n=5/group) were injected *i.p.* with PR8 plus 5-OP-RU or vehicle. Three days later, CD69^+^ MAIT cell frequencies in the lungs and the geometric mean fluorescence intensity (gMFI) of CD69 staining were determined by flow cytometry. ** denotes significant differences with *p* ≤ 0.01 by unpaired *t*-tests.

**Supplementary Fig. 6:**
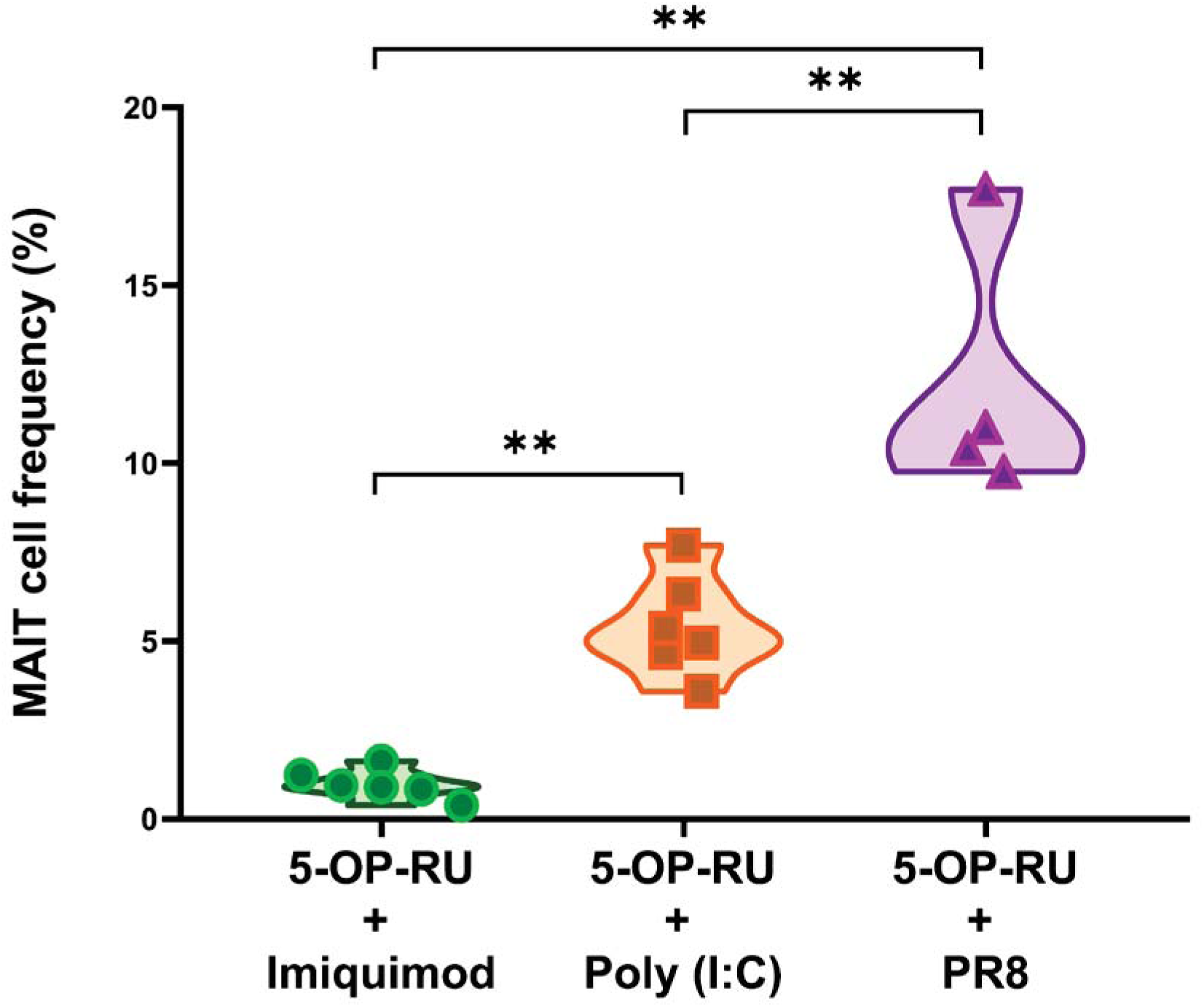
Poly (I:C), but not imiquimod, partially simulates PR8 when combined with 5-OP-RU to expand MAIT cells. B6-MAIT^CAST^ mice were injected *i.p.* with 5-OP-RU in combination with PR8, poly (I:C) (50 µg/mouse) or imiquimod (50 µg/mouse). Three days later, peritonealx MAIT cell frequencies were determined by flow cytometry. Each symbol represents an individual animal. ** denotes significant differences with *p* ≤ 0.01 by the Mann-Whitney *U* test.

**Supplementary Fig. 7:**
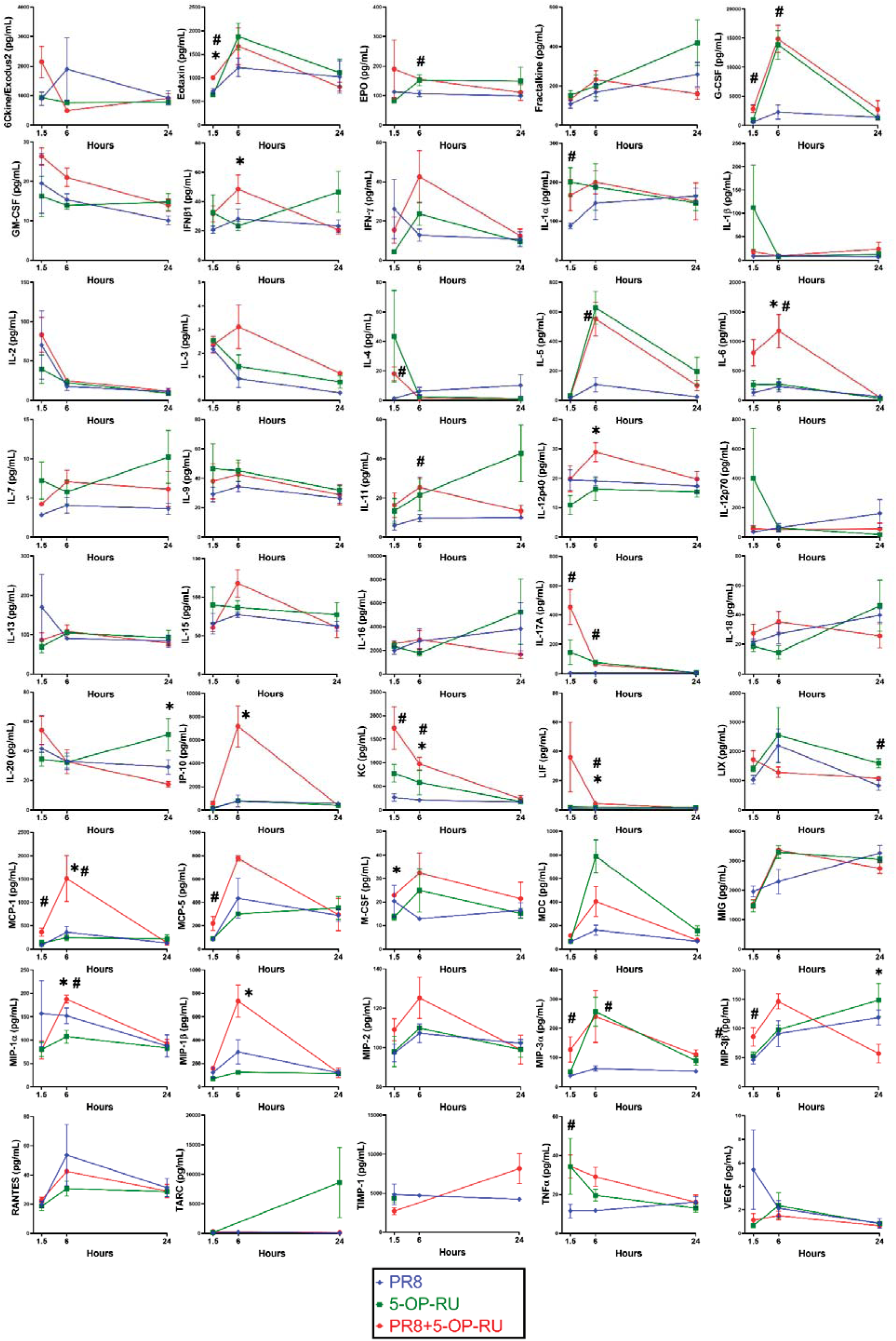
Cytokine serum levels after intraperitoneal administration of PR8 and/or 5-OP-RU. B6-MAIT^CAST^ mice were injected *i.p.* with PR8 (n=12), 5-OP-RU (n=12), or both (n=12). After 1.5, 6 or 24 hours, animals were sacrificed (n=4/timepoint) and their serum cytokine levels were measured. Statistical comparisons were made using unpaired *t*-tests. Asterisks (*) denote significant differences between sera from mice receiving PR8 plus 5-OP-RU and those injected with 5-OP-RU only. Hashtags (#) signify differences between animals receiving PR8 plus 5-OP-RU and those inoculated with PR8 only.

**Supplementary Fig. 8:**
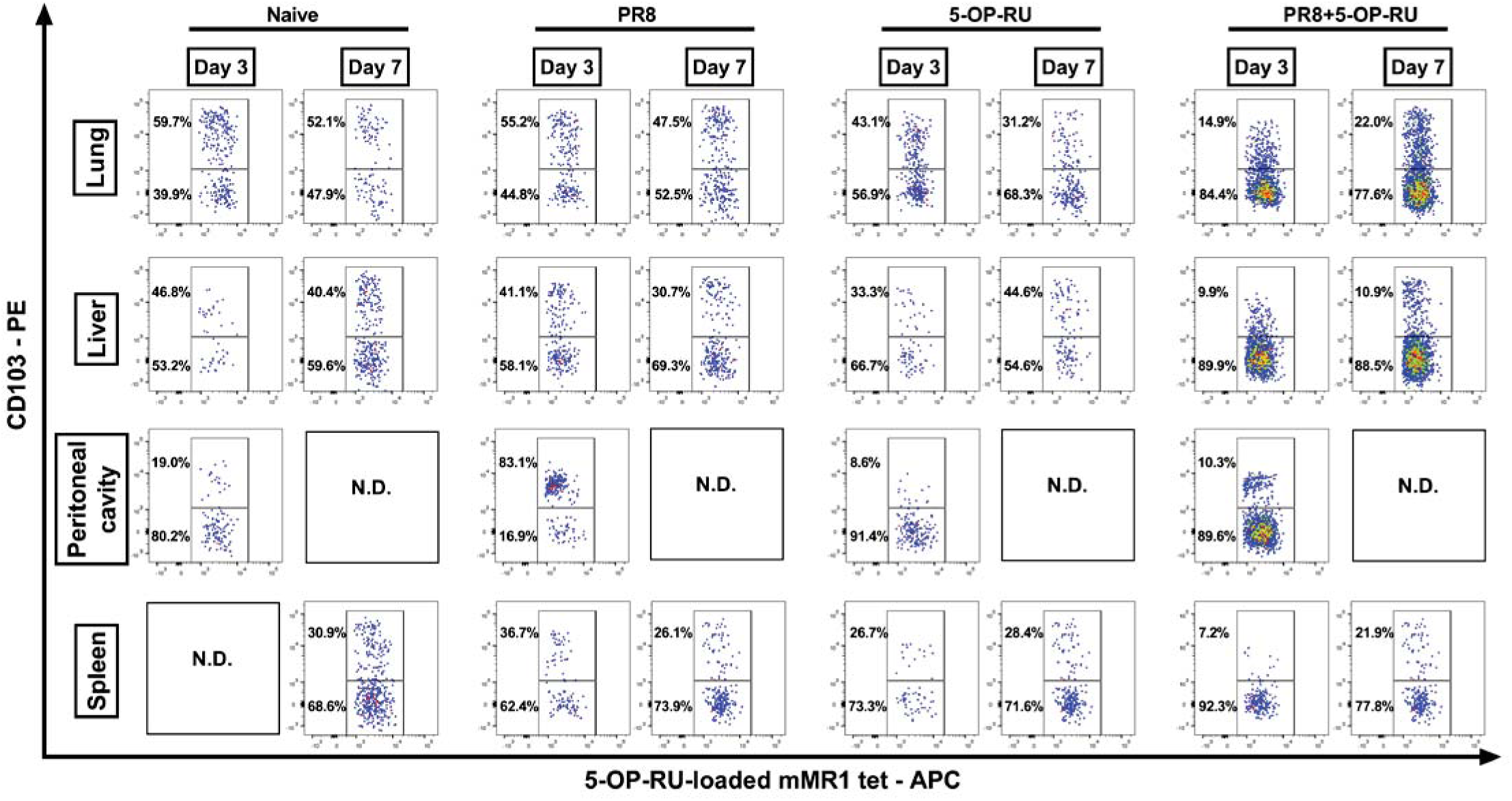
Tissue MAIT cells expanded via intraperitoneal immunization with PR8 and 5-OP-RU are low CD103 expressors. Naïve B6-MAIT^CAST^ mice and those receiving PR8, 5-OP-RU or both 3 or 7 days earlier were sacrificed, and pulmonary, hepatic, peritoneal and splenic MAIT cells were assessed by flow cytometry for CD103 expression. Representative plots after gating on TCRβ^+^ cells are illustrated. N.D. Not Determined.

**Supplementary Fig. 9:**
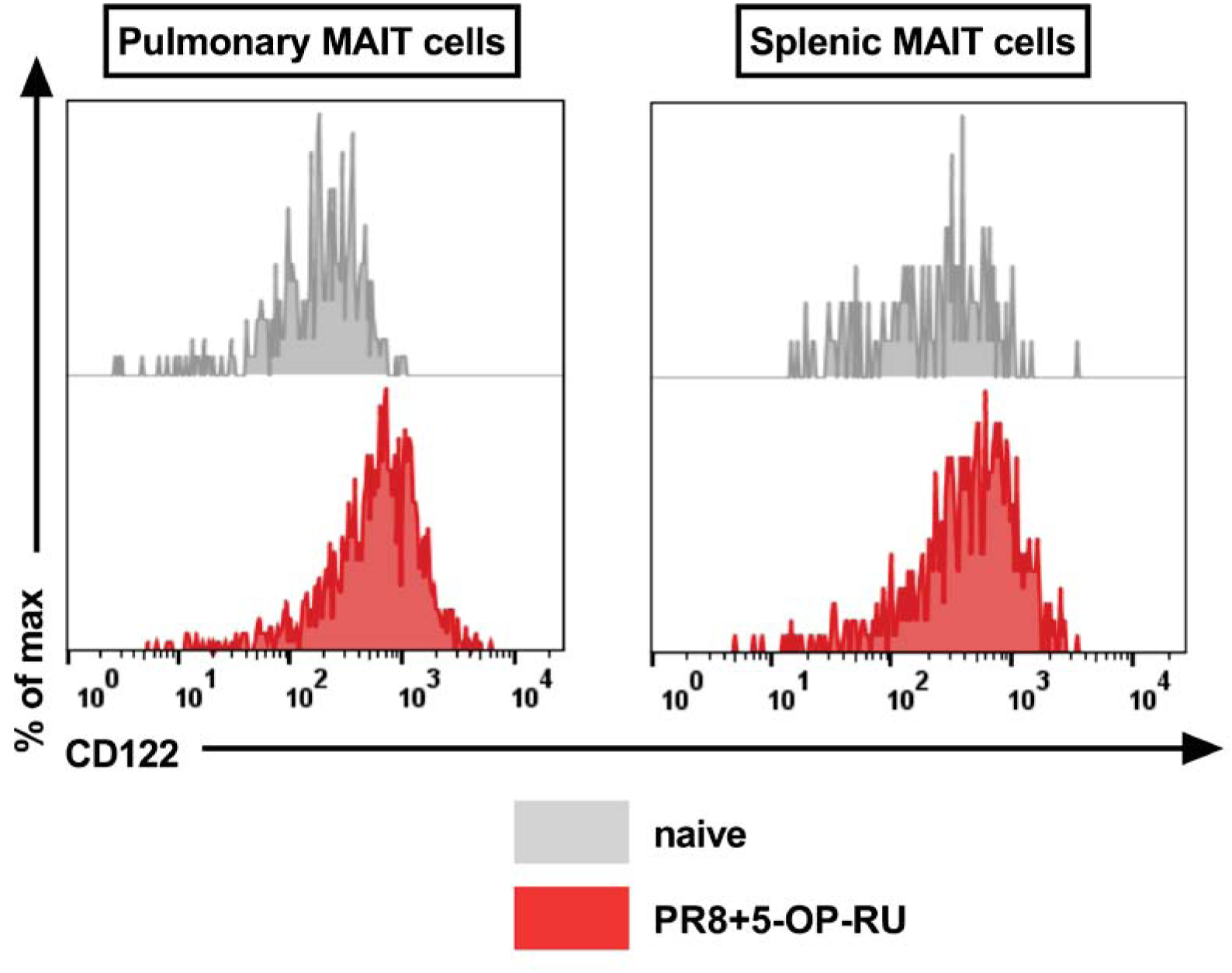
MAIT cells expanded after intraperitoneal co-administration of PR8 and 5-OP-RU abundantly express CD122 on their surface. Naïve B6-MAIT^CAST^ mice and animals receiving PR8 plus 5-OP-RU three days earlier were sacrificed, and pulmonary and splenic MAIT cells were examined cytofluorimetrically for their CD122 expression. Histograms corresponding to naïve mice are overlaid with those from immunized animals.

**Supplementary Fig. 10:**
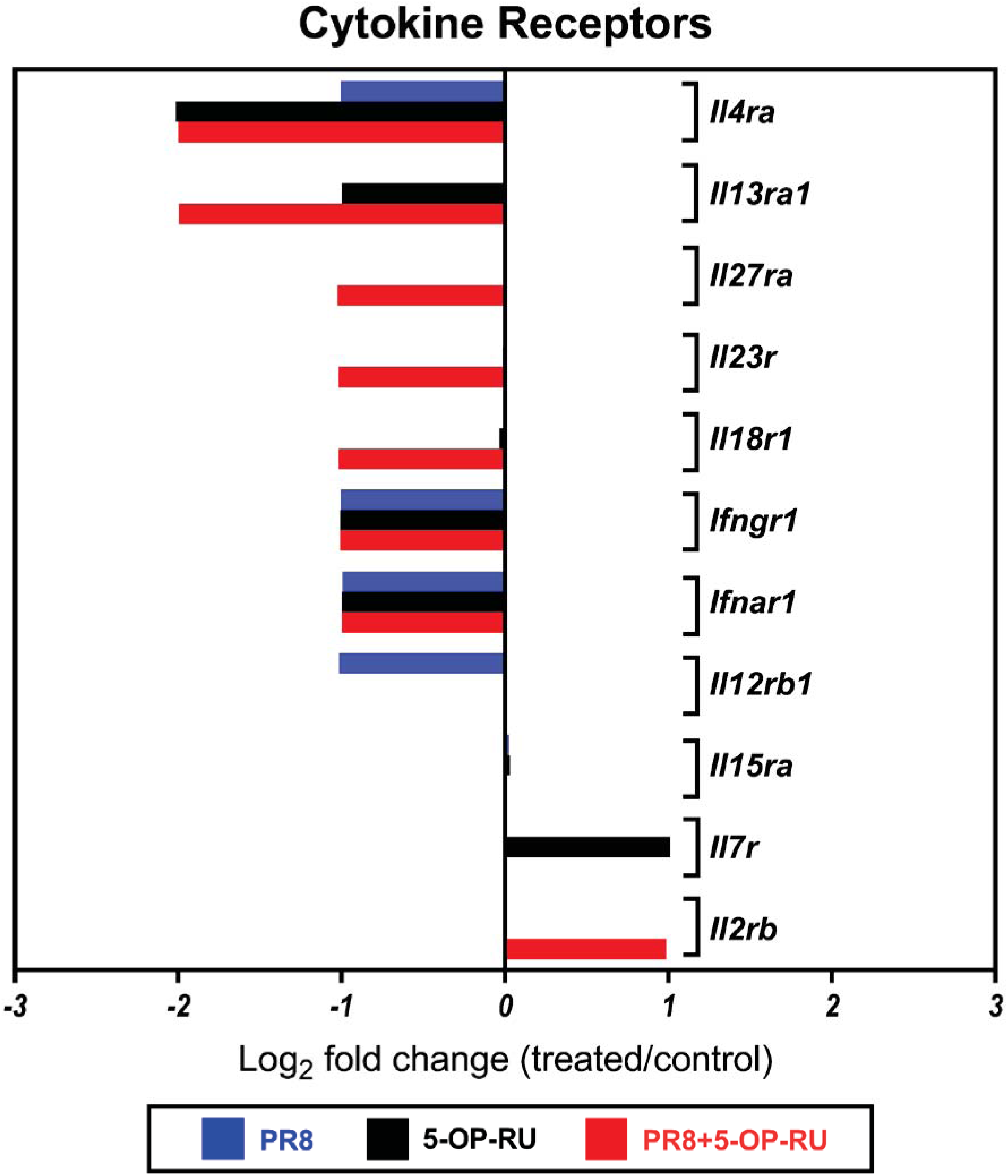
Cytokine receptor gene expression by pulmonary MAIT cells after intraperitoneal administration of PR8 and/or 5-OP-RU. B6-MAIT^CAST^ mice were given an *i.p.* injection of PBS (n=10), PR8 (n=10), 5-OP-RU (n=10), or PR8 plus 5-OP-RU (n=5). Three days later, pooled pulmonary MAIT cells were purified and assessed for their transcript levels of indicated cytokine receptors by quantitative PCR. Gene expression fold changes for each cohort relative to the control (PBS-injected) cohort were calculated using the 2^-(ΔΔCt)^ method.

**Supplementary Fig. 11:**
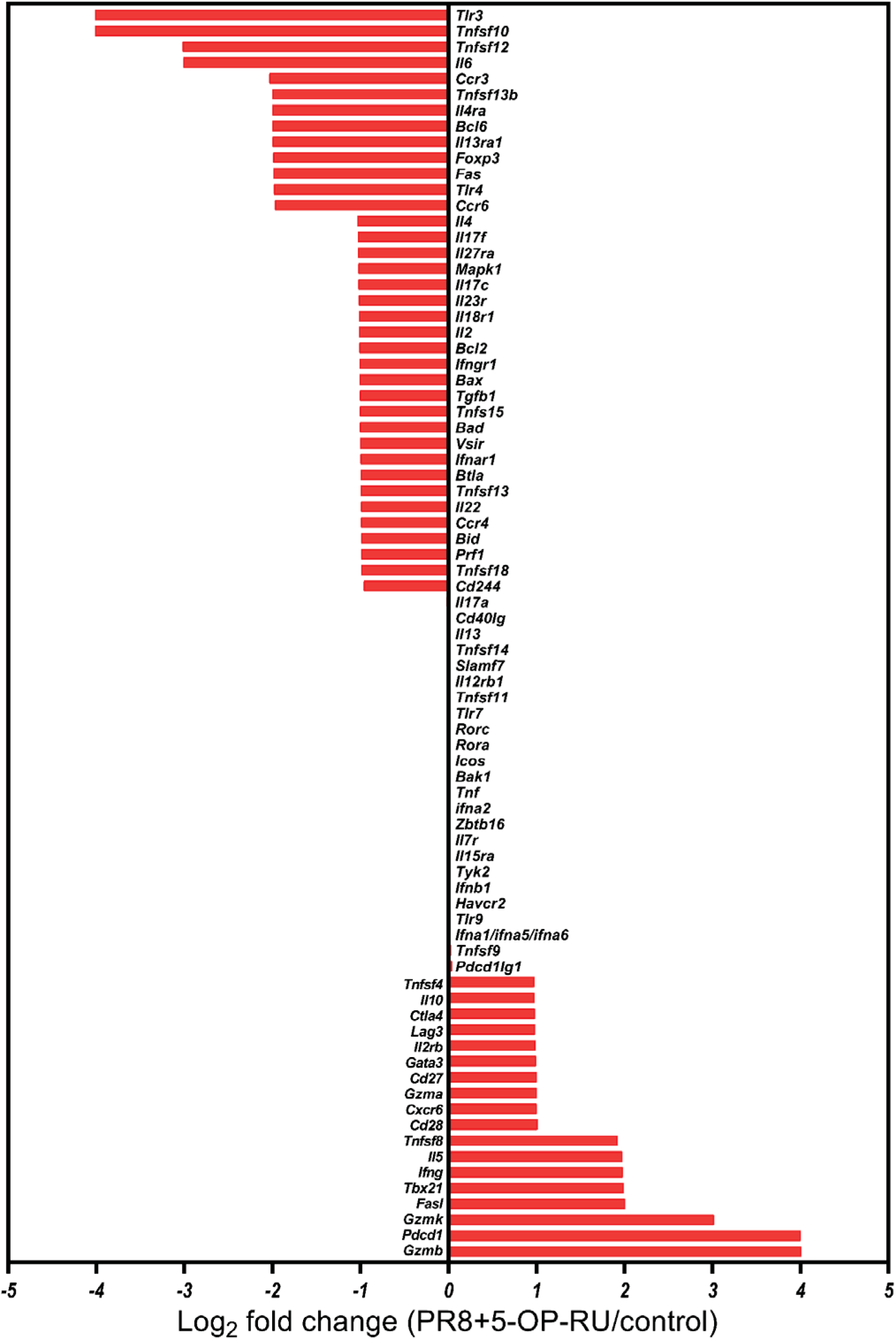
Comprehensive gene expression analysis of pulmonary MAIT cells after intraperitoneal administration of PR8, 5-OP-RU, or both. B6-MAIT^CAST^ mice were injected *i.p.* with PBS (n=10), PR8 (n=10), 5-OP-RU (n=10), or PR8 plus 5-OP-RU (n=5). Three days later, pooled pulmonary MAIT cells were isolated and examined by real-time PCR to quantitate the transcript levels of indicated molecules. Gene expression fold changes for each cohort were calculated relative to the control (PBS-injected) cohort using the 2^-(ΔΔCt)^ method.

**Supplementary Fig. 12:**
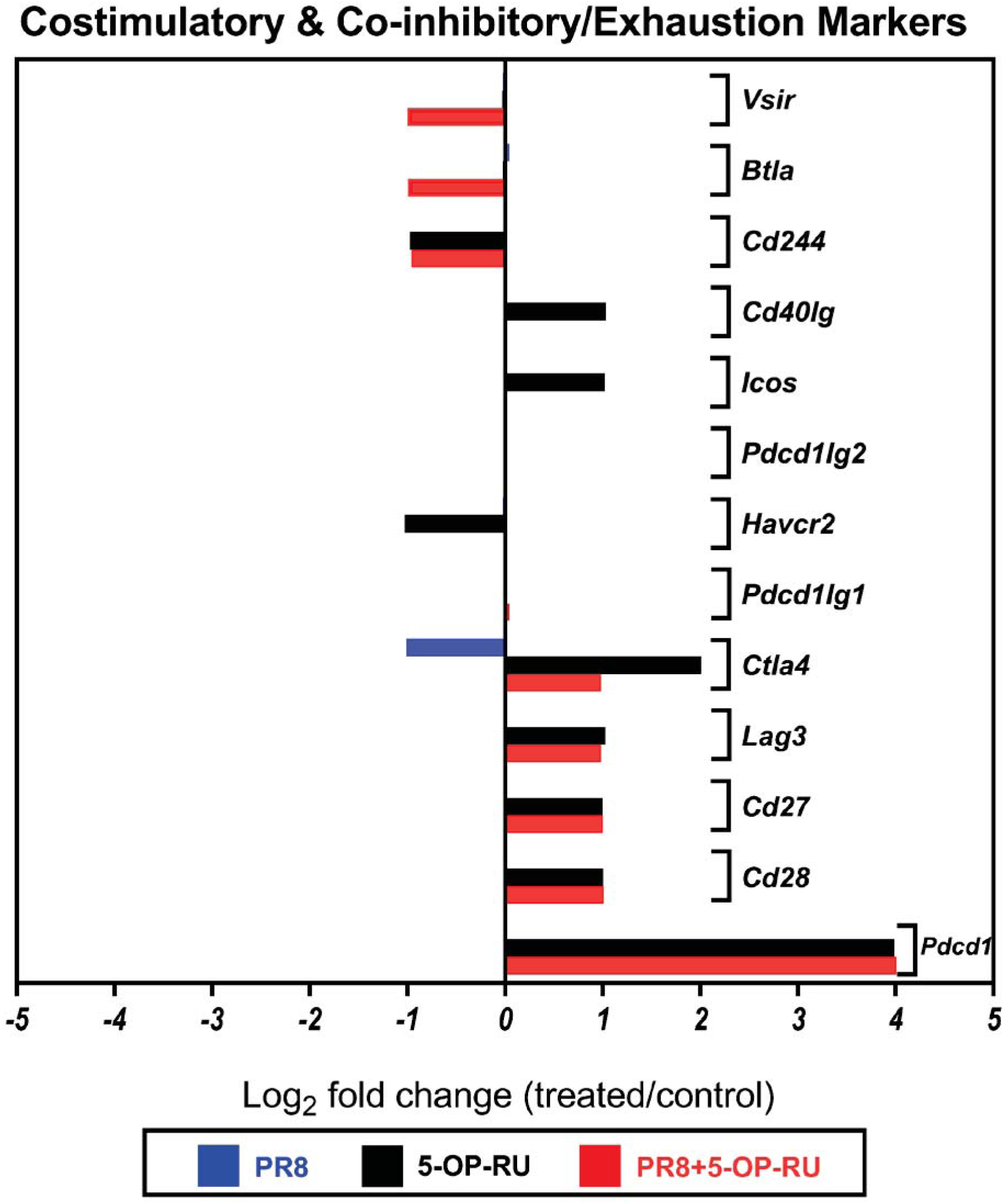
Transcriptional examination of classic costimulatory, co-inhibitory and exhaustion-associated molecules expressed by pulmonary MAIT cells after intraperitoneal administration of PR8 and/or 5-OP-RU. B6-MAIT^CAST^ mice were given an *i.p.* injection of PBS (n=10), PR8 (n=10), 5-OP-RU (n=10), or PR8 plus 5-OP-RU (n=5). After 3 days, pulmonary MAIT cells were purified, pooled and assessed by quantitative PCR for their transcript levels of indicated molecules. Gene expression fold changes for each cohort were calculated in comparison with the PBS-injected (control) cohort as described in Materials and Methods.

**Supplementary Fig. 13.**
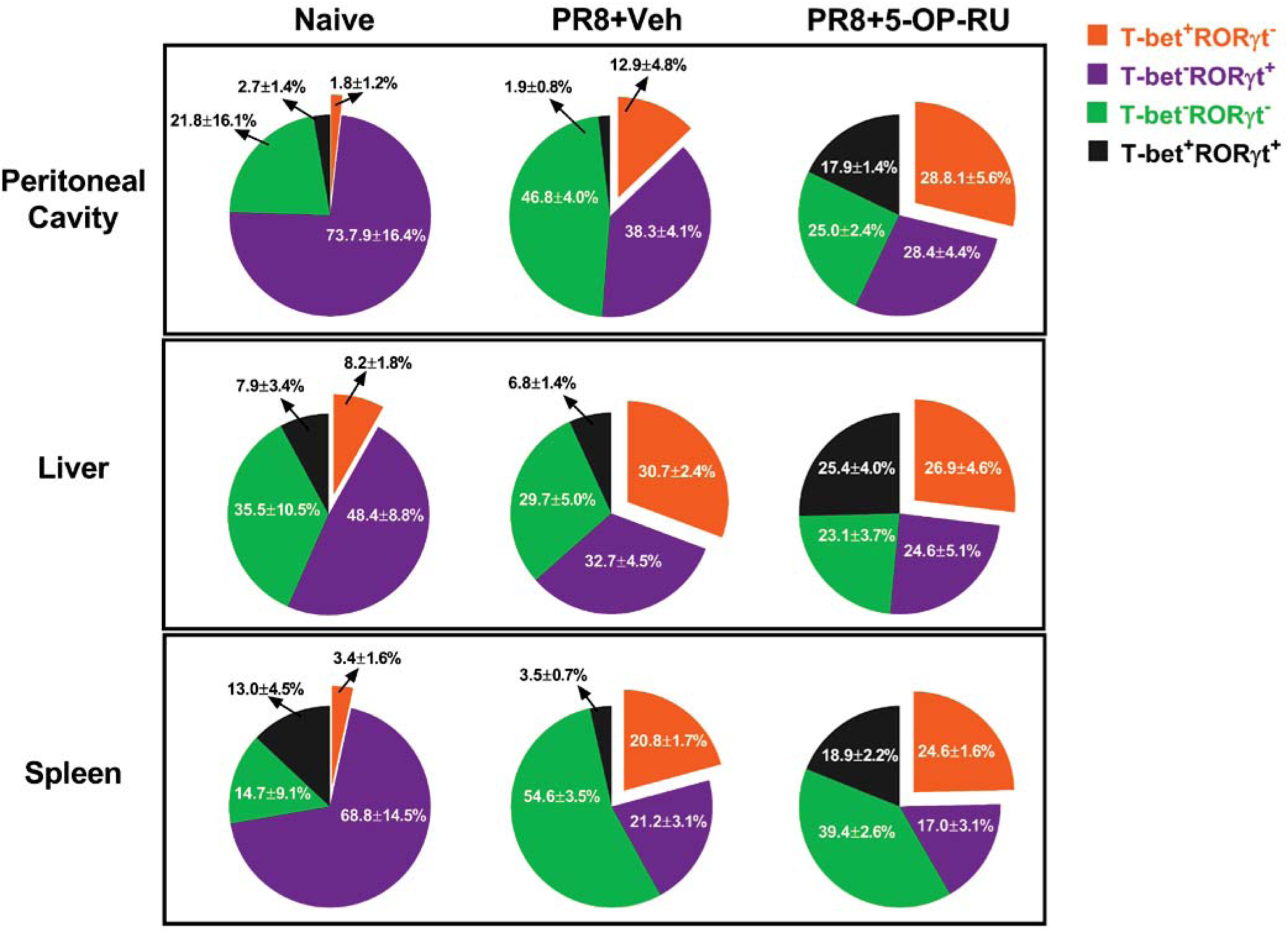
Intraperitoneal immunization with PR8 plus 5-OP-RU skews peritoneal, hepatic and splenic MAIT cells towards a T-bet^+^RORγt^-^ phenotype. Naïve B6-MAIT^CAST^ mice (n=5) and animals that had been inoculated 3 days earlier with PR8 plus 5-OP-RU (or vehicle) (n=6/group) were sacrificed. Peritoneal, hepatic and splenic MAIT cells were then stained for intracellular T-bet and RORγt. Pie charts visualize the frequencies (± SEM) of T-bet^+^RORγt^-^, T-bet^-^RORγt^+^, T-bet^+^RORγt^+^ and T-bet^-^RORγt^-^ MAIT cell subsets.

**Supplementary Fig. 14:**
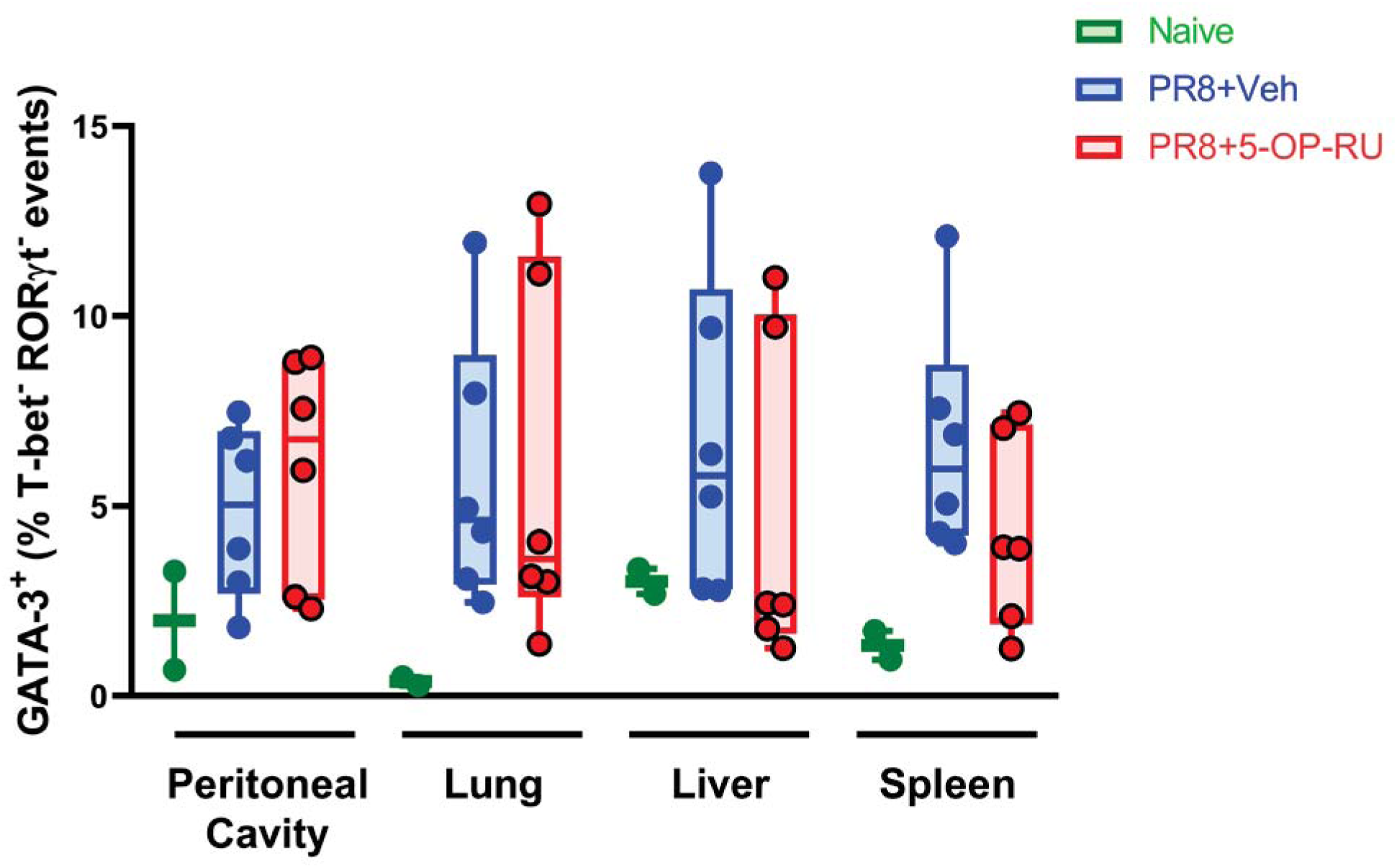
PR8 and 5-OP-RU co-administration does not alter GATA-3^+^ MAIT cell frequencies in different tissues. Two naïve B6-MAIT^CAST^ mice and 12 animals that had received an *i.p.* injection of PR8 plus 5-OP-RU (or vehicle) (n=6/group) three days earlier were sacrificed. Peritoneal, pulmonary, hepatic and splenic MAIT cells were interrogated by flow cytometry for their intracellular T-bet, RORγt and GATA-3 levels. Box-and-Whisker plots demonstrate GATA-3^+^ cell percentages among T-bet^-^RORγt^-^ MAIT cells.

**Supplementary Fig. 15:**
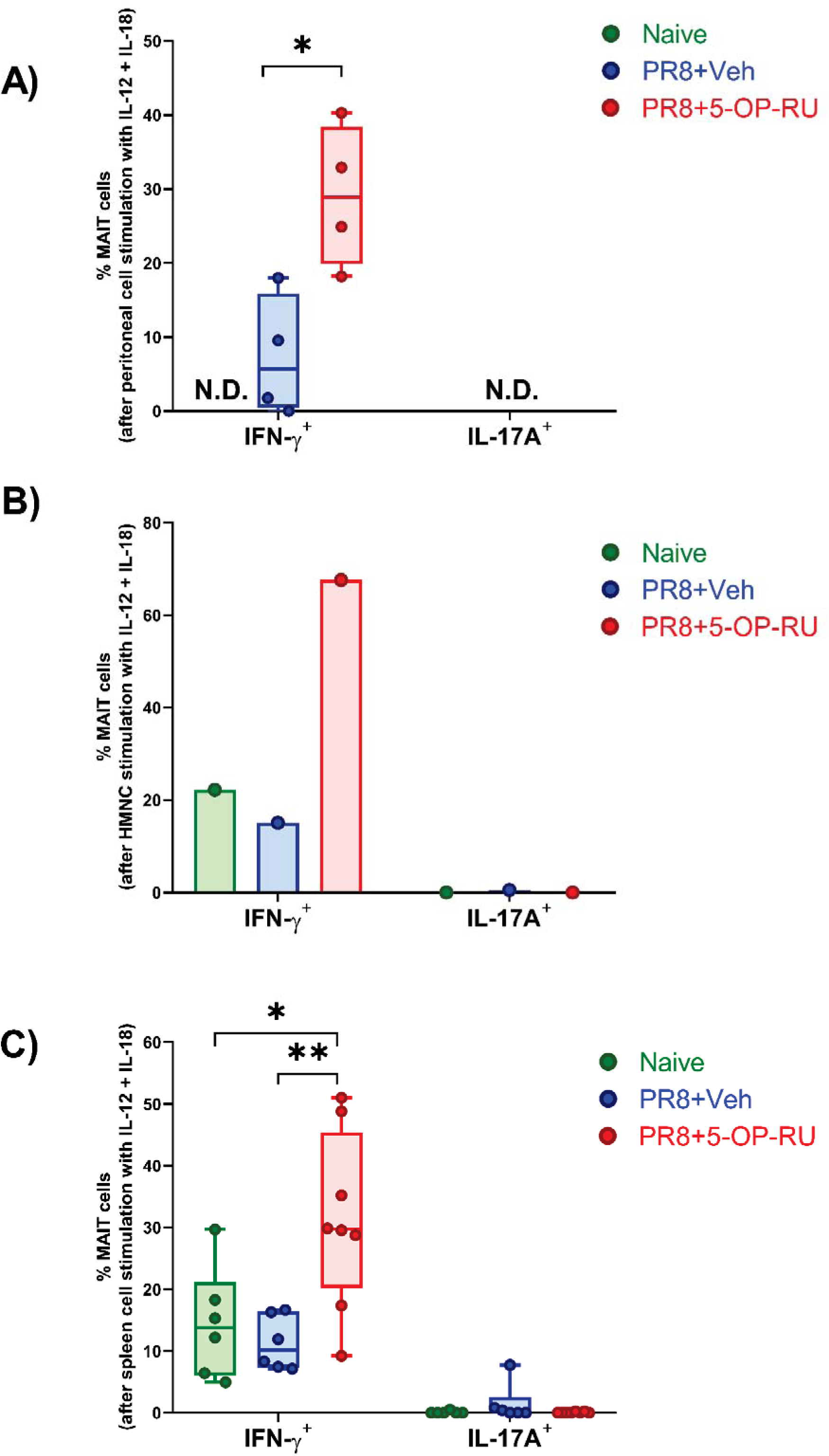
*In vivo* co-administration of PR8 and 5-OP-RU gives rise to a higher percentage of MAIT cells capable of producing IFN-γ, but not IL-17A, in response to *ex vivo* stimulation with IL-12 and IL-18. Unfractionated peritoneal cells (**A**), non-parenchymal hepatic mononuclear cells (HMNCs) (**B**) and spleen cells (**C**) were obtained from indicated numbers of naïve and immunized B6-MAIT^CAST^ mice and stimulated *ex vivo* with recombinant mouse IL-12 and IL-18. After 18 hours, IFN-γ^+^ and IL-17A^+^ cell frequencies among TCRβ^+^ MR1 tetramer^+^ MAIT cells were determined by flow cytometry. Each circle corresponds to an individual sample. * and ** denote differences with *p* ≤ 0.05 and *p* ≤ 0.01, respectively, by unpaired *t*-tests (**A**) and one-way ANOVA followed by the Tukey’s Multiple Comparisons test (**C**).

**Supplementary Fig. 16:**
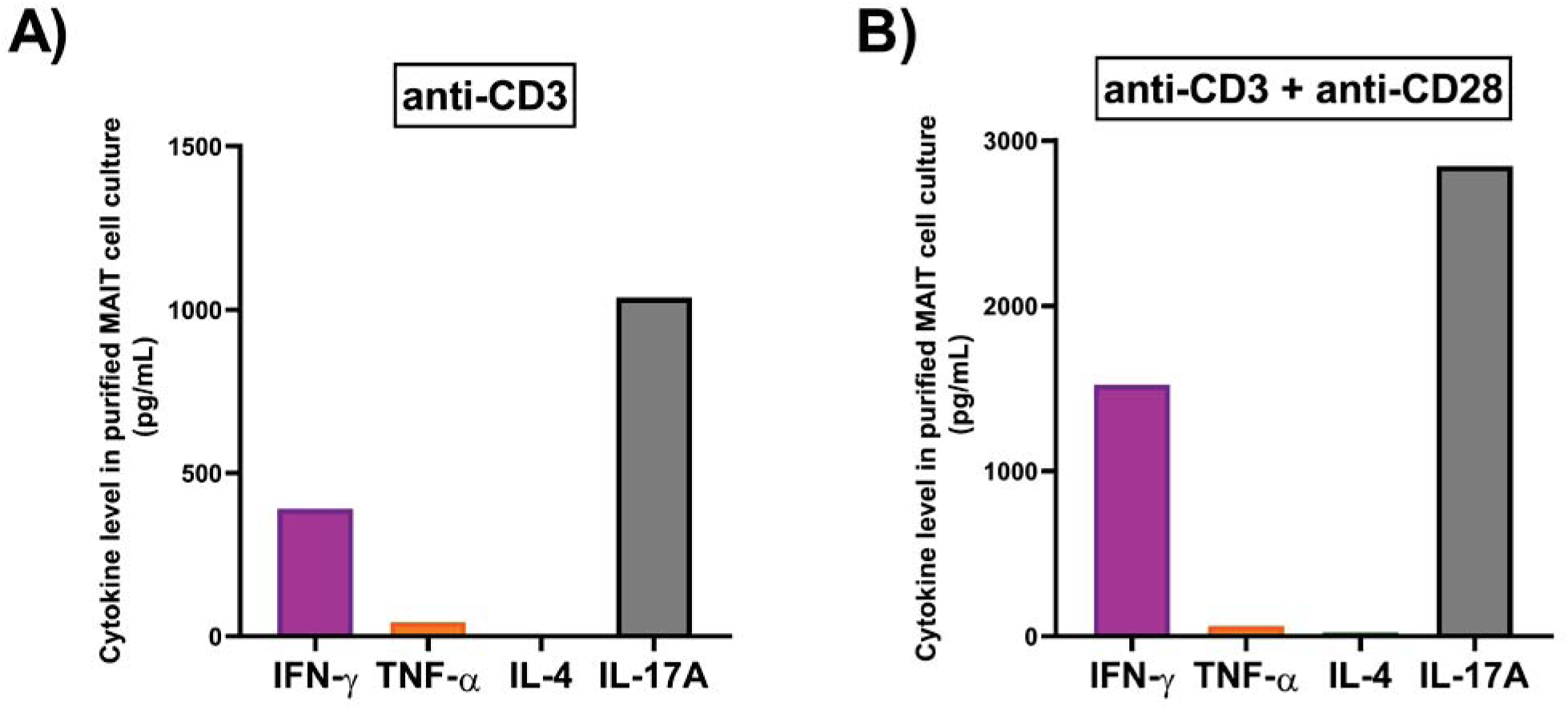
Prior immunization with PR8 and 5-OP-RU does not compromise MAIT cells’ ability to respond to anti-CD3 alone or in combination with anti-CD28. Purified pulmonary MAIT cells were pooled from 10 B6-MAIT^CAST^ mice that had been inoculated *i.p.* with PR8 and 5-OP-RU three days earlier. Cells were stimulated with plate-coated anti-CD3 in the absence (**A**) or presence (**B**) of soluble anti-CD28 as described in Materials and Methods. Eighteen hours later, indicated cytokines were measured in culture supernatants by ELISA.

**Supplementary Fig. 17:**
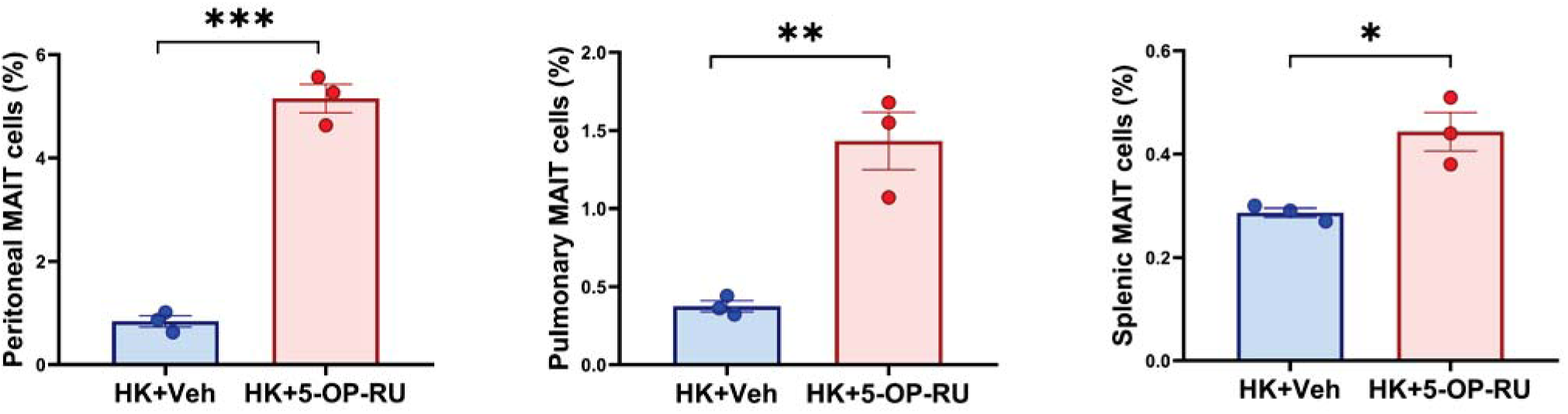
Intraperitoneal co-administration of A/Hong Kong/1/1968 and 5-OP-RU expands MAIT cells in several locations, including in the lungs. B6 mice (n=3/group) were injected *i.p.* with the A/Hong Kong/1/1968 (HK) strain of IAV plus 5-OP-RU or vehicle. Three days later, peritoneal (left panel), pulmonary (middle panel) and splenic (right panel) MAIT cells were enumerated by flow cytometry. Unpaired *t*-tests were used for statistical comparisons. *, ** and *** indicate differences with *p* ≤ 0.05, *p* ≤ 0.01 and *p* ≤ 0.001, respectively.

**Supplementary Fig. 18:**
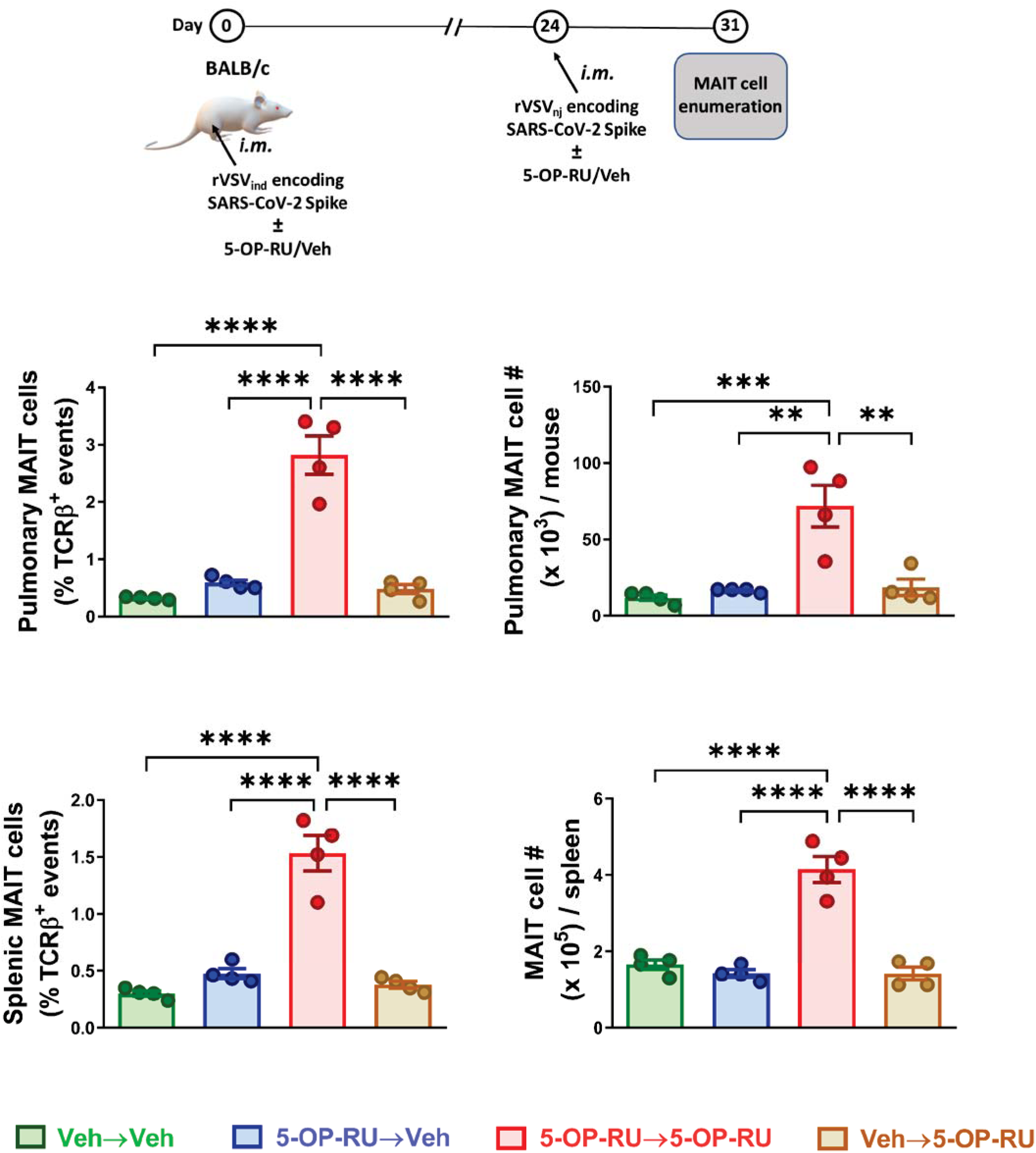
Robust MAIT cell expansion is achieved when 5-OP-RU is included during both primary and secondary immunizations with rVSV-based vaccines against SARS-CoV-2. BALB/c mice (n=4 per group) were primed intramuscularly (*i.m.*) with rVSV_Ind_- Spike and boosted 24 days later (also *i.m.*) with rVSV_NJ_-Spike. 5-OP-RU or vehicle was added to each vaccine inoculum. The frequencies (left panels) and absolute numbers (right panels) of pulmonary (upper panels) and splenic (lower panels) MAIT cells were determined on day 7 post-secondary immunization. Data are shown as mean ± SEM. **, *** and **** denote differences with *p* ≤ 0.01, *p* ≤ 0.001 and *p* ≤ 0.0001, respectively, by one-way ANOVA followed by Tukey’s post-hoc Multiple Comparisons analyses.

**Supplementary Fig. 19:**
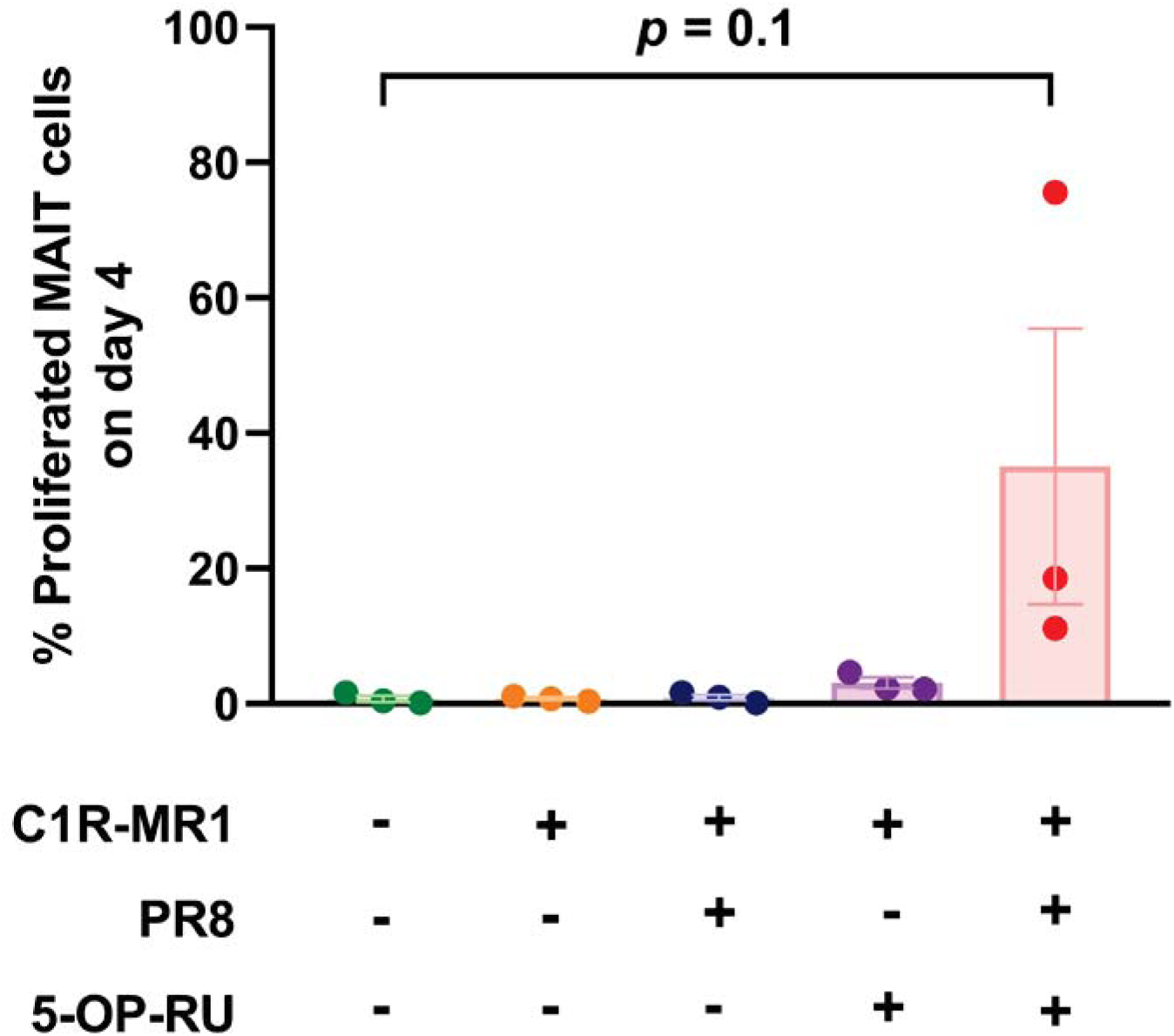
Simultaneous stimulation of human PBMCs for 4 days with PR8 and 5-OP-RU appears to expand MAIT cells. CFDA-SE-labeled human PBMCs from 3 healthy donor PBMCs were rested in complete medium or stimulated with C1R-MR1 cells that had been infected with PR8 and/or pulsed with 5-OP-RU. Four days later, the extent of CFDA-SE dye dilution among MAIT cells was examined by flow cytometry. Data are shown as mean ± SEM.

**Supplementary Fig. 20:**
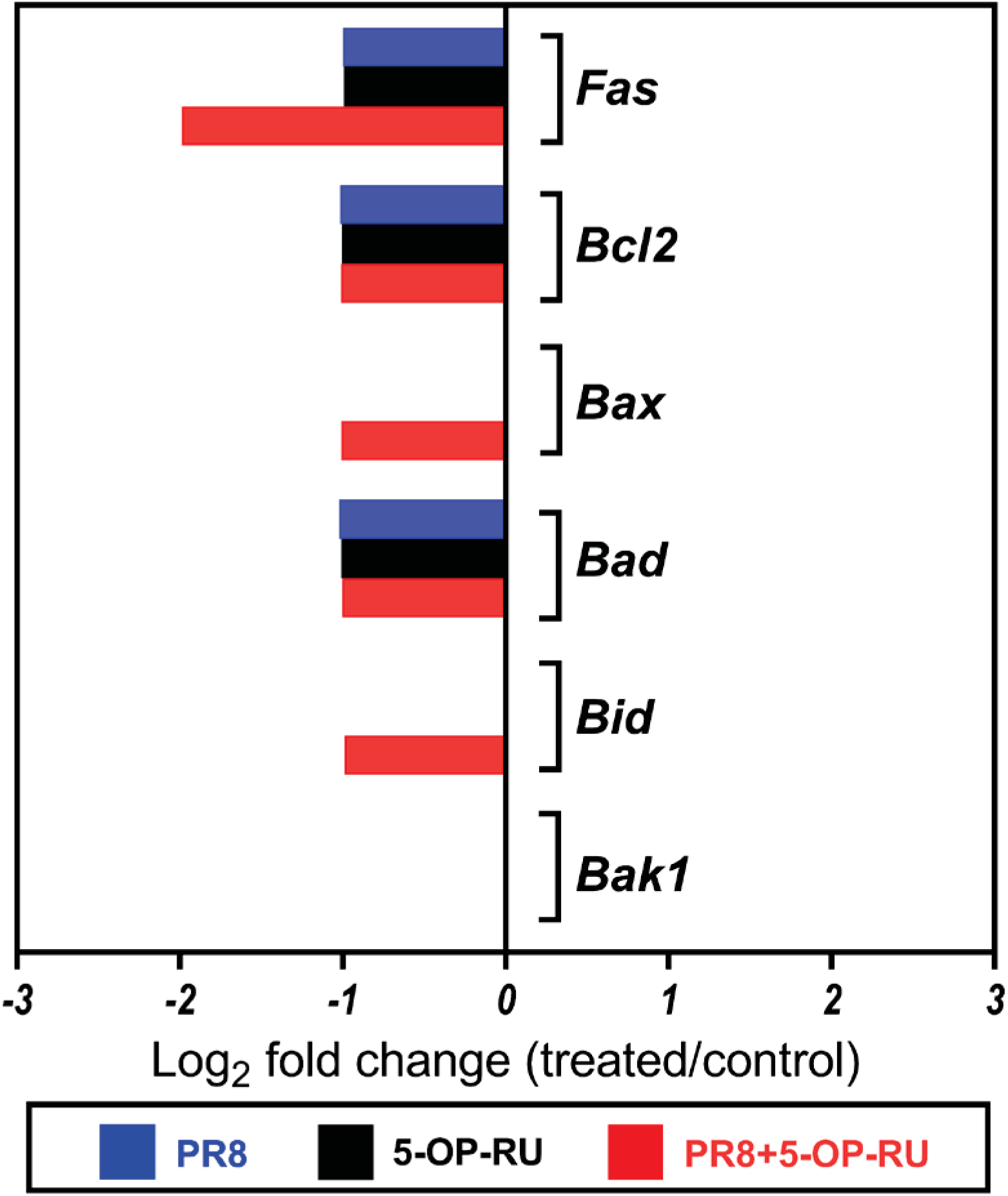
Transcriptional assessment of pro/anti-survival gene expression by pulmonary MAIT cells after intraperitoneal administration of PR8 and/or 5-OP-RU. B6-MAIT^CAST^ mice were injected with PBS (n=10), PR8 (n=10), 5-OP-RU (n=10), or PR8 plus 5-OP-RU (n=5). Three days later, pooled pulmonary MAIT cells were evaluated by real-time PCR for their transcript levels of indicated molecules. Gene expression fold changes for each cohort relative to the control (PBS-injected) cohort were calculated as described in Materials and Methods.

**Supplementary Fig. 21:**
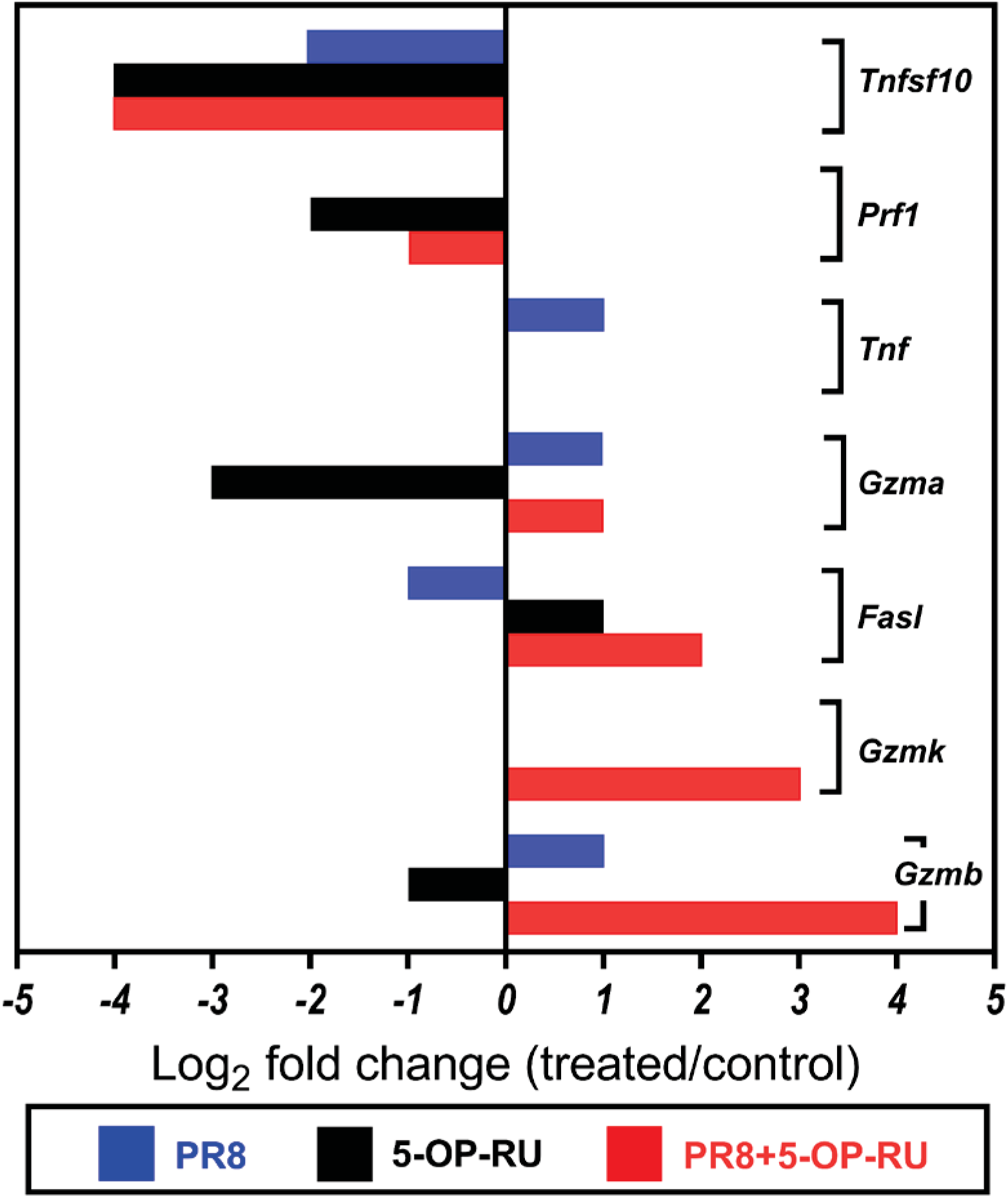
Gene expression analysis of cytolytic effector molecules expressed by pulmonary MAIT cells following intraperitoneal injection of PR8 and/or 5-OP-RU. B6-MAIT^CAST^ mice were injected with PBS (n=10), PR8 (n=10), 5-OP-RU (n=10), or PR8 plus 5-OP-RU (n=5). After 3 days, pulmonary MAIT cells were purified, pooled and assessed by real-time PCR for their transcript levels of listed molecules. Gene expression fold changes were determined for each cohort compared with the PBS-injected (control) cohort as described in Materials and Methods.

**Supplementary Table 1.** TaqMan-based quantitative PCR primer/probe sets used in this investigation.

| **Target Gene** | **Assay Identifier** |
| --- | --- |
| *Actb* | Mm00607939_s1 |
| *B2m* | Mm00437762_m1 |
| *Bad* | Mm00432042_m1 |
| *Bak1* | Mm00432045_m1 |
| *Bax* | Mm00432051_m1 |
| *Bcl2* | Mm00477631_m1 |
| *Bcl6* | Mm00477633_m1 |
| *Bid* | Mm00432073_m1 |
| *Btla* | Mm00616981_m1 |
| *Ccr3* | Mm00515543_s1 |
| *Ccr4* | Mm01963217_u1 |
| *Ccr6* | Mm99999114_s1 |
| *Cd27* | Mm01185212_g1 |
| *Cd28* | Mm00483137_m1 |
| *Cd244* | Mm00479575_m1 |
| *Cd40lg* | Mm00441911_m1 |
| *Ctla4* | Mm00486849_m1 |
| *Cxcr6* | Mm02620517_s1 |
| *Fas* | Mm01204974_m1 |
| *Fasl* | Mm00438864_m1 |
| *Foxp3* | Mm00475162_m1 |
| *Gapdh* | Mm99999915_g1 |
| *Gata3* | Mm00484683_m1 |
| *Gzma* | Mm01304452_m1 |
| *Gzmb* | Mm00442834_m1 |
| *Gzmk* | Mm00492530_m1 |
| *Havcr2* | Mm01294183_m1 |
| *Hmbs* | Mm00660262_g1 |
| *Hprt1* | Mm00446968_m1 |
| *Icos* | Mm00497600_m1 |
| *Ifna1/Ifna5/Ifna6* | Mm03030145_gH |
| *Ifna2* | Mm00833961_s1 |
| *Ifna4* | Mm00833969_s1 |
| *Ifnb1* | Mm00439552_s1 |
| *Ifng* | Mm01168134_m1 |
| *Ifnar1* | Mm00439544_m1 |
| *Ifngr1* | Mm00599890_m1 |
| *Il2* | Mm00434256_m1 |
| *Il4* | Mm00445259_m1 |
| *Il5* | Mm00439646_m1 |
| *Il6* | Mm00446190_m1 |
| *Il9* | Mm00434305_m1 |
| *Il10* | Mm00439614_m1 |
| *Il13* | Mm00434204_m1 |
| *Il17a* | Mm00439618_m1 |
| *Il17b* | Mm01258783_m1 |
| *Il17c* | Mm00521397_m1 |
| *Il17f* | Mm00521423_m1 |
| *Il22* | Mm01226722_g1 |
| *Il2rb* | Mm00434268_m1 |
| *Il4ra* | Mm00439634_m1 |
| *Il12rb1* | Mm00434189_m1 |
| *Il13ra1* | Mm00446726_m1 |
| *Il15ra* | Mm04336046_m1 |
| *Il18r1* | Mm00515178_m1 |
| *Il23r* | Mm00519943_m1 |
| *Il27ra* | Mm00497259_m1 |
| *Ipo8* | Mm01255158_m1 |
| *Lag3* | Mm00493071_m1 |
| *Mapk1* | Mm00442479_m1 |
| *Pdcd1* | Mm01285676_m1 |
| *Pdcd1lg1* | Mm03048248_m1 |
| *Pdcd1lg2* | Hs00228839_m1 |
| *Pgk1* | Mm00435617_m1 |
| *Prf1* | Mm00812512_m1 |
| *Rora* | Mm00443103_m1 |
| *Rorc* | Mm01261022_m1 |
| *Rplp2* | Mm00782638_s1 |
| *Slamf7* | Mm00513808_m1 |
| *Tbp* | Mm00446973_m1 |
| *Tbx21* | Mm00450960_m1 |
| *Tfrc* | Mm00441941_m1 |
| *Tgfb1* | Mm01178820_m1 |
| *Tlr3* | Mm01207404_m1 |
| *Tlr4* | Mm00445273_m1 |
| *Tlr7* | Mm04933178_g1 |
| *Tlr9* | Mm00446193_m1 |
| *Tnf* | Mm00443258_m1 |
| *Tnfsf4* | Mm00437214_m1 |
| *Tnfsf8* | Mm00437153_m1 |
| *Tnfsf9* | Mm00437155_m1 |
| *Tnfsf10* | Mm01283606_m1 |
| *Tnfsf11* | Mm00441906_m1 |
| *Tnfsf12* | Mm02583406_s1 |
| *Tnfsf13* | Mm03809849_s1 |
| *Tnfsf13b* | Mm00446347_m1 |
| *Tnfsf14* | Mm00444567_m1 |
| *Tnfsf15* | Mm00770031_m1 |
| *Tnfsf18* | Mm00839222_m1 |
| *Tyk2* | Mm00444469_m1 |
| *Ubc* | Mm01201237_m1 |
| *Vsir* | Mm00472312_m1 |
| *Xiap* | Mm01311594_mH |
| *Zbtb16* | Mm01176868_m1 |

**Supplementary Table 2.**
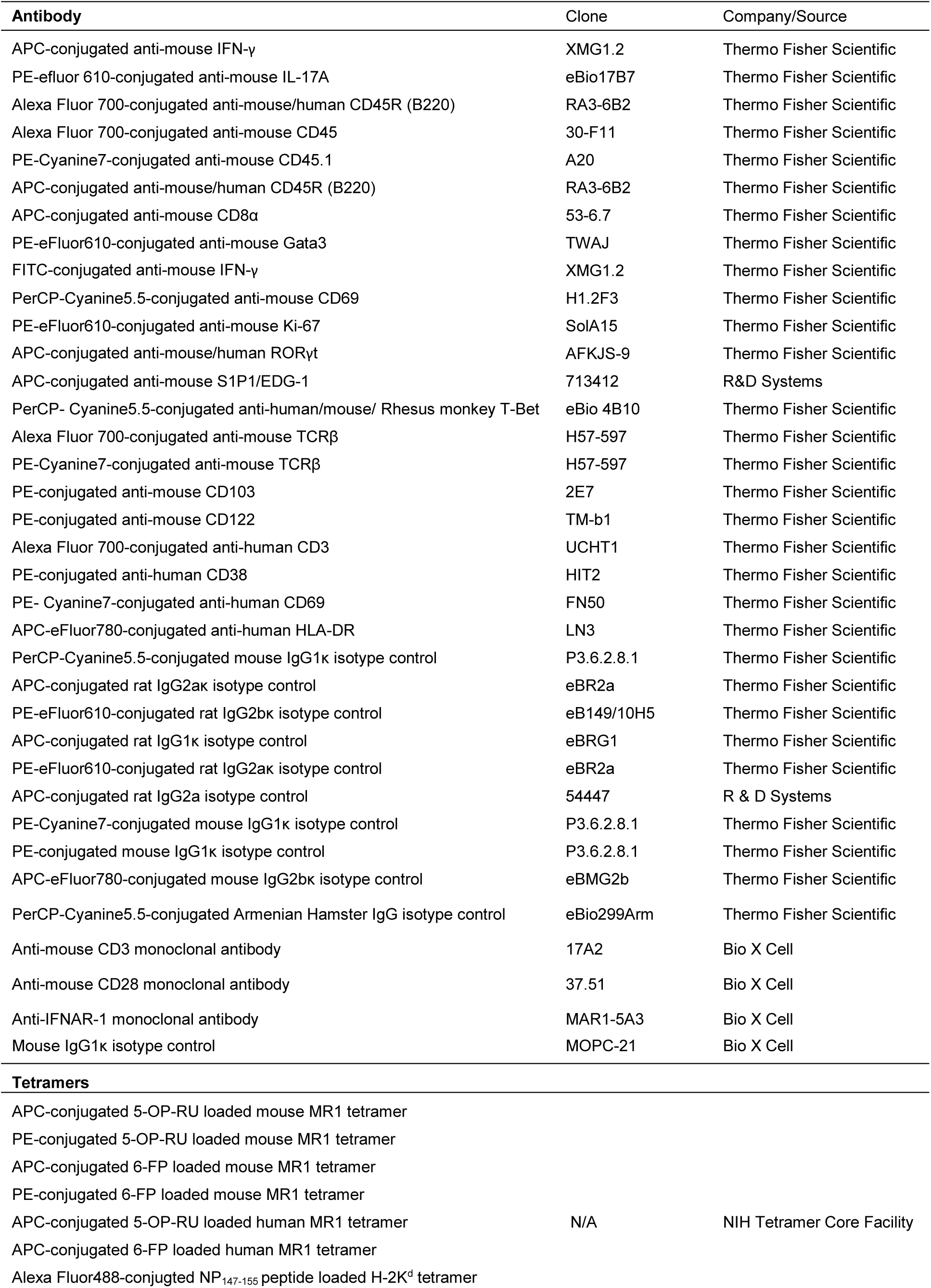
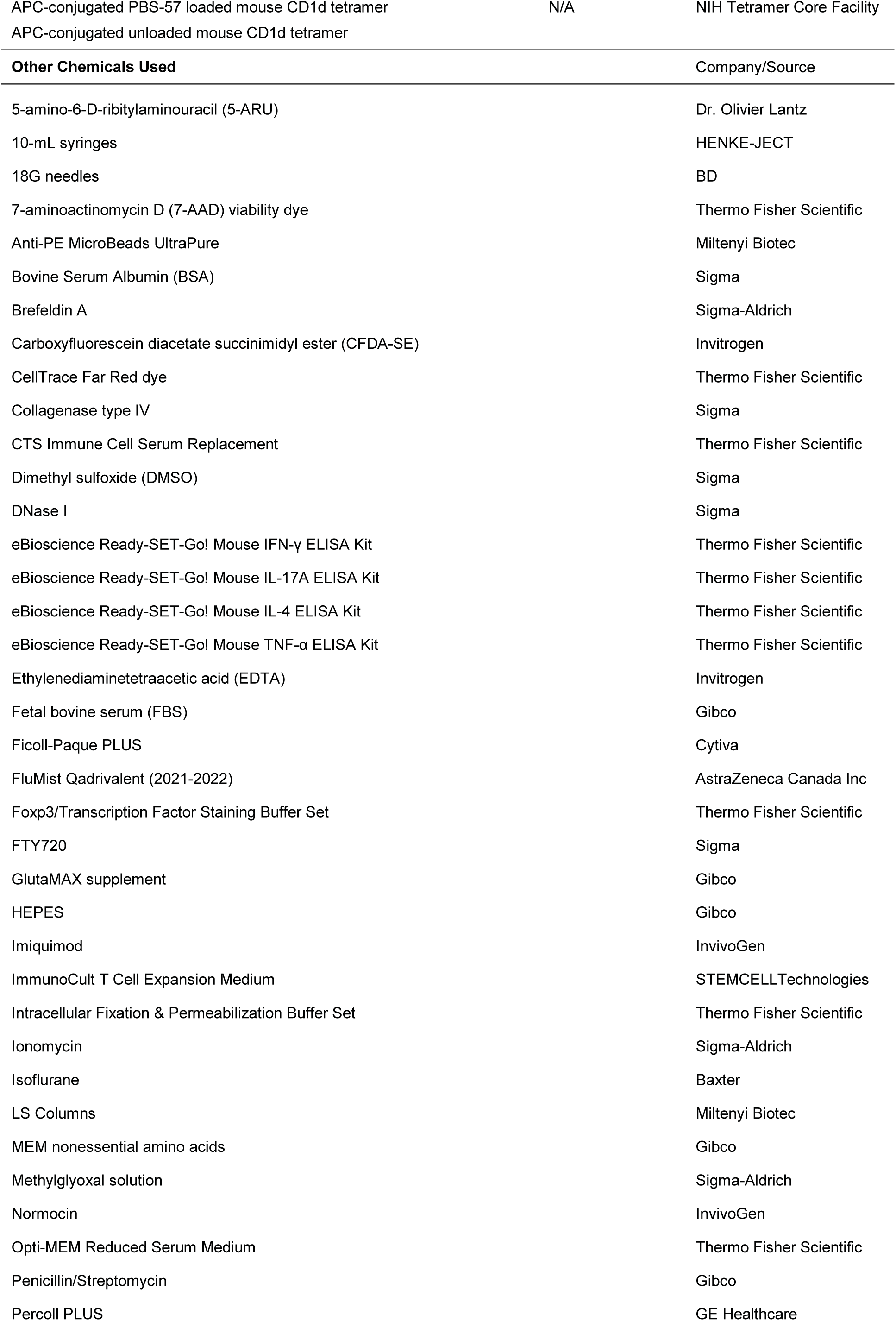
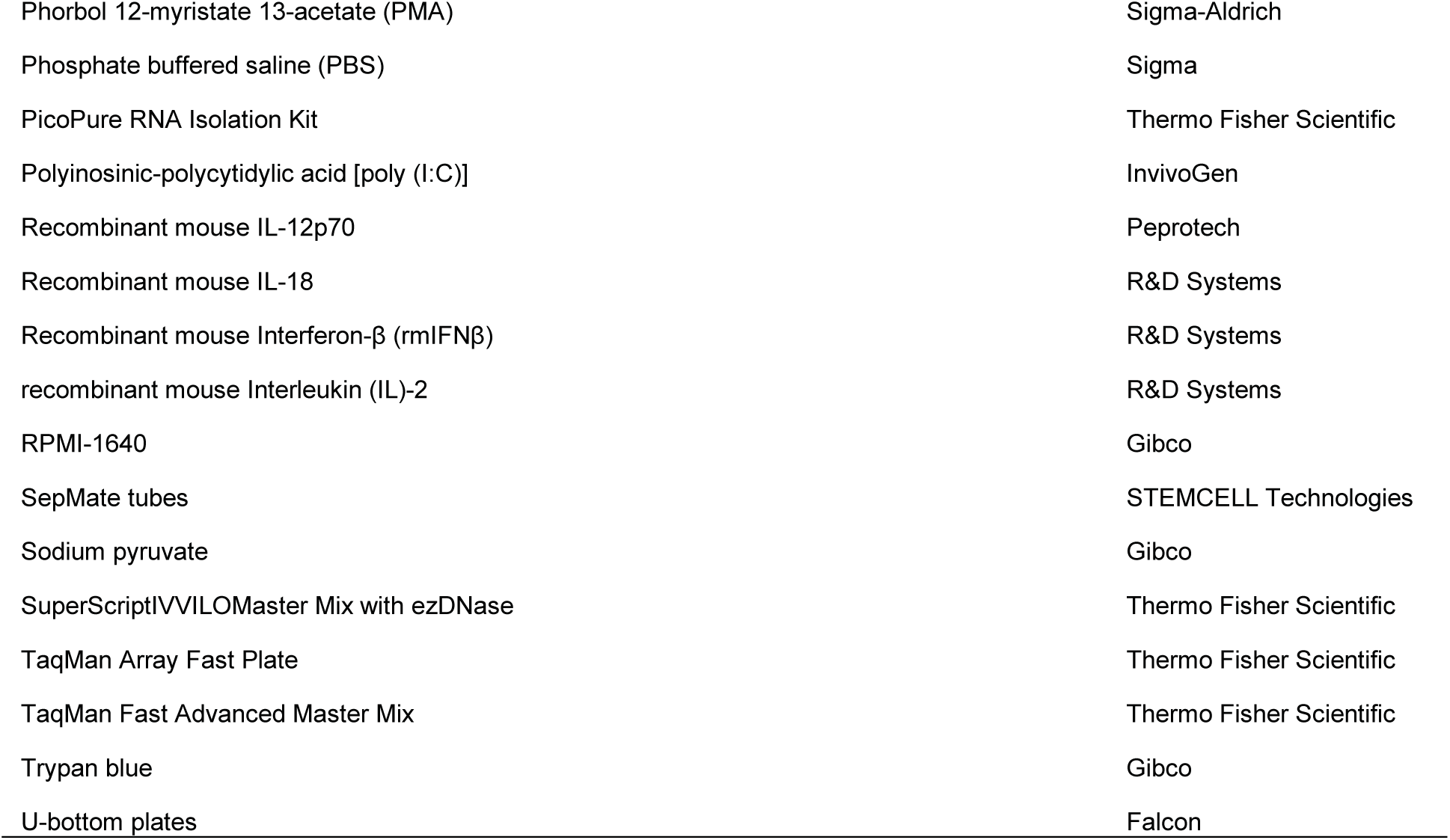
List of Antibodies, Tetramers and other reagents

